# Erythrocyte membrane protein 3 (EMAP3) is exposed on the surface of the *Plasmodium berghei* infected red blood cell

**DOI:** 10.1101/2024.05.28.596273

**Authors:** Sophia Hernandez, Ravish Rashpa, Thorey K. Jonsdottir, Martina S. Paoletta, Maria Rayón Diaz, Severine Chevalley-Maurel, Takahiro Ishizaki, Chris J. Janse, Blandine Franke-Fayard, Mathieu Brochet, Ellen SC Bushell

## Abstract

The human malaria parasite *Plasmodium falciparum* invades red blood cells (RBC) and exports parasite proteins to transform the host cell for its survival. These exported proteins facilitate uptake of nutrients and cytoadherence of the infected RBC (iRBC) to endothelial cells of small blood vessels, thus protecting the iRBC from splenic clearance. The parasite protein PfEMP1 and the host protein CD36 play a major role in *P. falciparum* iRBC cytoadherence. The murine parasite *Plasmodium berghei* is a widely used experimental model that combines high genetic tractability with access to *in vivo* studies. *P. berghe*i iRBC also sequesters in small blood vessels, mediated by binding to CD36. However, the parasite proteins binding to CD36 are unknown and only very few parasite proteins, including EMAP1 and EMAP2, have been identified that are present at the iRBC membrane. We have identified a new protein named EMAP3 and demonstrated its export to the iRBC membrane where it interacts with EMAP1, with only EMAP3 exposed on the outer surface of the iRBC. Parasites lacking EMAP3 display no significant reduction in growth or sequestration, indicating that EMAP3 is not the major CD36-binding protein. The outer-surface location of EMAP3 offers a new scaffold for displaying *P. falciparum* proteins on the surface of the *P. berghei* iRBC, providing a platform to screen *in vivo* putative inhibitors of *P. falciparum* cytoadherence.

## Introduction

The etiological agents of malaria are unicellular *Plasmodium* parasites that cause disease by invading and replicating within red blood cells (RBC), which ultimately leads to the destruction of the infected cells^1^. Upon invasion of the RBC the parasite proliferates within the protective parasitophorous vacuole and remodels the RBC to support its rapid growth by exporting a large range of parasite proteins across the parasitophorous vacuole membrane (PVM) into the infected RBC (iRBC)^2^. A number of malaria parasite species, including the human parasite *P. falciparum*, export proteins onto the surface of the iRBC that facilitate cytoadherence to vascular endothelium of small blood vessels and thereby evasion of clearance of the iRBC by the spleen. This causes parasites to accumulate in blood vessels of inner organs and contributes to disease pathology through blockage of blood capillaries, vascular leakage and endothelial inflammation, which correlates with severe malaria in infections with *P. falciparum*^3^. In *P. falciparum* endothelial cytoadherence and the resulting organ sequestration is mediated by the parasite protein PfEMP1 (erythrocyte membrane protein 1), which is encoded by the *var* multigene family. This protein is the major parasite surface ligand binding the host protein CD36 that is expressed on the vascular endothelium. In addition, it binds to the host proteins iCAM-1 (intercellular adhesion molecule 1), as well as to CSA (chondroitin sulphate A) in the placenta of pregnant women^4^. The identification of new therapies blocking cytoadherence has the potential of being translated into much needed life-saving interventions that can be deployed to interrupt the rapid and often fatal disease progression characterising *P. falciparum* severe malaria. Experimentally studying *P. falciparum* cytoadherence and sequestration *in vivo* is challenging due to the lack of appropriate whole organism models.

*Plasmodium berghei* is a widely used murine experimental model that benefits from high genetic tractability and facilitates *in vivo* studies. The *P. berghei* iRBC also sequesters in small blood vessels, mediated by binding to CD36. Importantly, *P. berghei* infections can be monitored by whole body imaging, where both the parasite and host can be manipulated and the effect on sequestration and virulence can be directly assessed^5,6^. Despite being a commonly used model, the molecular mediators of *P. berghei* sequestration are not fully understood. There is no orthologue of *pfemp1* in *P. berghei*^7^. However, many fundamental aspects of parasite biology and virulence are conserved between *P. falciparum* and *P. berghei.* Significantly, P. *berghei* also exports a large and diverse array of parasite proteins into the infected RBC, including proteins encoded by multigene families^8,9^. The molecular machinery for protein export across the PVM is also shared between *P. falciparum* and *P. berghei* and experimental deletion of several of these conserved proteins in *P. berghei* results in reduced blood-stage growth and sequestration *in vivo*^10–15^. In addition, *P. berghei* induces the generation of membranous structures in the cytoplasm of iRBC, termed intra-erythrocytic *P. berghei*-induced structures (IBIS), akin to Maurer’s clefts in *P. falciparum* that mediate trafficking of proteins to the surface of the infected cell^16^. Importantly, some of the key molecular players of these membrane structures are also shared between *P. falciparum* and *P. berghei* ^17–19^.

None of the *P. berghei* exported proteins that are known to play a role in CD36-mediated sequestration are presented on the surface of the iRBC. Instead, they are located in the iRBC cytoplasm and therefore likely play a role in trafficking CD36-binding ligand(s) to the surface^5,17,20^. For example, deletion of the *P. berghei* gene encoding SMAC (schizont membrane-associated cytoadherence protein) results in slow growth of blood stage parasites *in vivo* and impaired CD36-mediated sequestration of iRBC. SMAC is exported into the iRBC but does not localise to the iRBC membrane^5,17^. In addition, exported proteins encoded by several different multigene families, including the large *pir* (*Plasmodium* interspersed repeat) family are located in the PV or in the cytoplasm of the iRBC with no clear evidence for an iRBC surface localisation, neither in *P. berghei* ^21^ or in another rodent malaria parasite *Plasmodium chabaudi*^21,22^. The *P. berghei* proteins EMAP1 and EMAP2 (erythrocyte membrane associated protein 1/2) are exported to the iRBC membrane of rodent malaria parasites. Although these two proteins have been identified based on their absence in iRBCs of a the non-sequestering laboratory strain of *P. berghei* (K173), they do not play a critical role in growth of blood stage parasites or iRBC sequestration and thus are not *P. berghei* surface ligands that mediates binding to CD36^8^.

PfEMP1 binds CD36 via its extracellular cysteine-rich interdomain region (CIDR) domain. There are no known *P. berghei* proteins with CIDR domains^5^. We therefore set out to identify by bioinformatic analyses *P. berghei* genes encoding novel exported proteins that are putatively expressed on the surface of the iRBC and that could mediate iRBC cytoadherence. These analyses resulted in the identification of a single copy gene coding for a small triple transmembrane domain protein that we named EMAP3. We demonstrate that in the sequestering schizont stage EMAP3 interacts with EMAP1, where EMAP3 is anchored to the iRBC membrane and displayed on the outer side of the iRBC membrane. Deletion of the gene encoding EMAP3 does not significantly affect asexual blood stage growth nor iRBC sequestration or disease characteristics. We conclude that EMAP3 is exposed on the outer side surface of the iRBC, but it is not the major ligand for CD36 mediated sequestration of *P. berghei* iRBC. Nevertheless, its extracellular exposure facilitates the use of EMAP3 as a scaffold to display *P. falciparum* proteins on the surface of the *P. berghei* iRBC for screening *in vivo* putative inhibitors of *P. falciparum* cytoadherence.

## Results

### EMAP3 is exported onto the iRBC surface during asexual blood stage development

The *emap3* gene (PBANKA_0825900) encodes a small protein (352 amino acids) that lacks functional domain prediction, has three predicted transmembrane domains at the N-terminus and is conserved among different human, simian, avian and rodent malaria parasites (https://plasmodb.org/)^23^. The N-terminal positioning of the three transmembrane domains is preserved in syntenic orthologues of the human malaria parasites *Plasmodium vivax*, *Plasmodium knowlesi* and *Plasmodium ovale*^23^. EMAP3 (but not the two other EMAP proteins) is predicted to have transmembrane domains but it lacks a signal peptide motif. Only EMAP1 has a known functional domain, a pyst-A domain that has been implicated in lipid binding and transport^8,24^ (**Fig 1A**).

**Figure 1.**
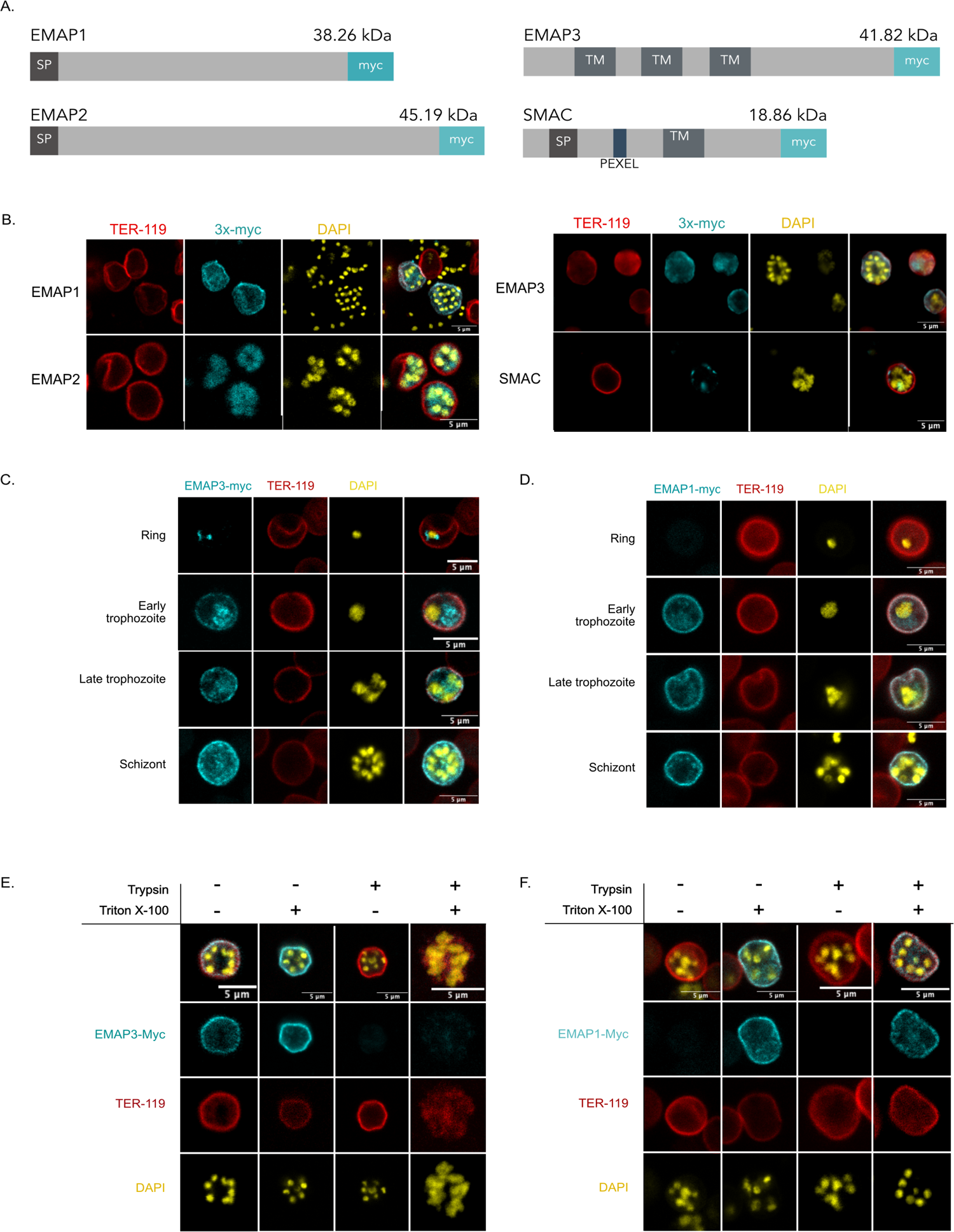
EMAP3 is exposed on the surface of *P. berghei* iRBC during the schizont stage. **(A)** Schematic presentation of genes encoding EMAP1, EMAP2, EMAP3 and SMAC with C-terminal 3x-myc tag (myc). *Plasmodium* export element (PEXEL), transmembrane domain (TM) and signal peptide (SP). Molecular weight of proteins given in kilodalton (kDa), excluding myc-tag. **(B)** Immunofluorescence assays (IFA) of schizonts expressing 3x-myc tagged EMAP1, EMAP2, EMAP3 and SMAC. **(C-D)** Time course IFA of (C) EMAP3-myc and (D) EMAP1-myc expression in different blood stages (ring, early trophozoite, late trophozoite, schizonts). **(E-F)** The effect of permeabilisation using Triton X-100 and trypsin treatment on the detection of (E) EMAP3-myc and (F) EMAP1-myc in *P. berghei* schizonts by IFA. In all microscopy panels the myc-tagged protein is visualised in cyan, together with co-staining for TER-119 to label the red blood cell membrane (red) and DNA labelled by 4′,6-diamidino-2-phenylindole (DAPI), (yellow). Scale bar = 5 µm.

To determine the subcellular location of EMAP3 we generated a *P. berghei* mutant with a C-terminal triple (3x-myc) tag in the endogenous *emap3* locus by standard genetic modification technology^25^. In addition, we generated *P. berghei* mutants with 3x-myc tagged EMAP1 (PBANKA_0836800) or EMAP2 (PBANKA_0316800) that are known to be exported to the iRBC membrane and SMAC (PBANKA_0100600), which is known to be involved in *P. berghei* iRBC sequestration^5,8^. Correct 3x-myc tagging was confirmed by diagnostic genotyping PCR (**Fig. S1**). Expression of the tagged proteins in blood stage parasites (schizonts) was verified by immunofluorescence assays (IFA) detecting the 3x-myc tagged proteins (Fig. 1B). In schizonts EMAP3-myc and EMAP1-myc show a location largely concentrated to the iRBC membrane, while EMAP2-myc displayed a more diffuse localisation throughout the iRBC cytoplasm. SMAC-myc was also concentrated at the iRBC periphery but, in contrast to EMAP3 and EMAP1, it is present in distinct foci seemingly lining the inside of the iRBC (Fig. 1B).

To further investigate the location and timing of expression of EMAP3 we took samples from cultures containing roughly synchronised blood stages (ring, early trophozoite, late trophozoite, and schizont) and performed IFA to detect EMAP3-myc and EMAP1-myc. EMAP3 is expressed already in rings, where it appears closely associated with the nucleus (likely still residing in or close to the endoplasmic reticulum, ER). Already in early trophozoites EMAP3-myc is found at the iRBC membrane and EMAP3 remains located at the iRBC membrane in schizonts (Fig. 1C). The secretion of EMAP3 and the specificity of the EMAP3-myc signal at the iRBC membrane was verified by treatment with brefeldin A (BFA), which traps parasite proteins during transport between the ER and Golgi apparatus^26^. As expected, BFA treatment inhibited export of EMAP3-myc and abolished EMAP3-myc iRBC surface staining in mature parasite stages (schizonts / trophozites). In earlier parasite stages (rings), EMAP3-myc accumulated in close proximity to the parasite ER (visualised by co-staining of the ER-resident BiP (binding immunoglobulin protein), in both control and BFA treated parasites (**Fig. S2**). EMAP3-myc expression and location closely mirrors that of EMAP1-myc, which is also exported to the iRBC membrane in trophozoites and schizonts. A notable difference is that EMAP1-myc is not detectable in rings as has been previously reported^8^ (Fig. 1D).

EMAP3 is predicted to have three transmembrane domains at its N-terminus, with the bulk of the protein downstream of the final transmembrane domain (Fig. 1A). We hypothesised that the C-terminus contains functional domain(s) that may play a role in interacting with host cell proteins outside the iRBC when exposed on the surface of the iRBC. We therefore set out to identify how the protein is oriented on the iRBC membrane through surface shaving using trypsin and permeabilisation with Triton X-100. EMAP3-myc is detectable without prior permeablisation, demonstrating that the C-terminal end of the protein containing the 3x-myc tag is exposed on the iRBC surface. This is supported by the observation that the EMAP3-myc signal is lost upon treatment with trypsin (Fig. 1E). In contrast, EMAP1-myc could only be detected when the cell is permeabilised (Fig. 1F), indicating that the C-terminus of EMAP1 is not exposed on the surface of the iRBC. Despite its association with the iRBC membrane, EMAP1 lacks predicted transmembrane domains (Fig. 1A) and the orientation of the protein at the iRBC membrane was not further studied.

To examine the location of EMAP1, EMAP3 and SMAC more closely, we performed ultrastructure expansion microscopy (UEx-M). To analyse the orientation and location at the iRBC membrane we stained the iRBC with BODIPY TR ceramide to visualise membranes and/or N-hydroxysuccinimide (NHS)-ester that binds to protein dense regions. UEx-M analysis confirms that EMAP3-myc localises to the iRBC membrane in mature schizonts, where it co-localised with the BODIPY-stained RBC membrane (Fig. 2A). EMAP1-myc could also be observed in close proximity to the iRBC membrane, however the EMAP1-myc staining in UEx-M was more widespread and was present more widely across the host cell cytoplasm (Fig. 2B). In addition, in agreement with IFA results, SMAC-myc was concentrated in distinct patches, seemingly located just under the iRBC membrane (Fig. 2C). Taken together, these analyses show that EMAP3 is exported into the iRBC where it is anchored as a multi-pass transmembrane domain protein with its C-terminus exposed on the surface of the iRBC.

**Figure 2.**
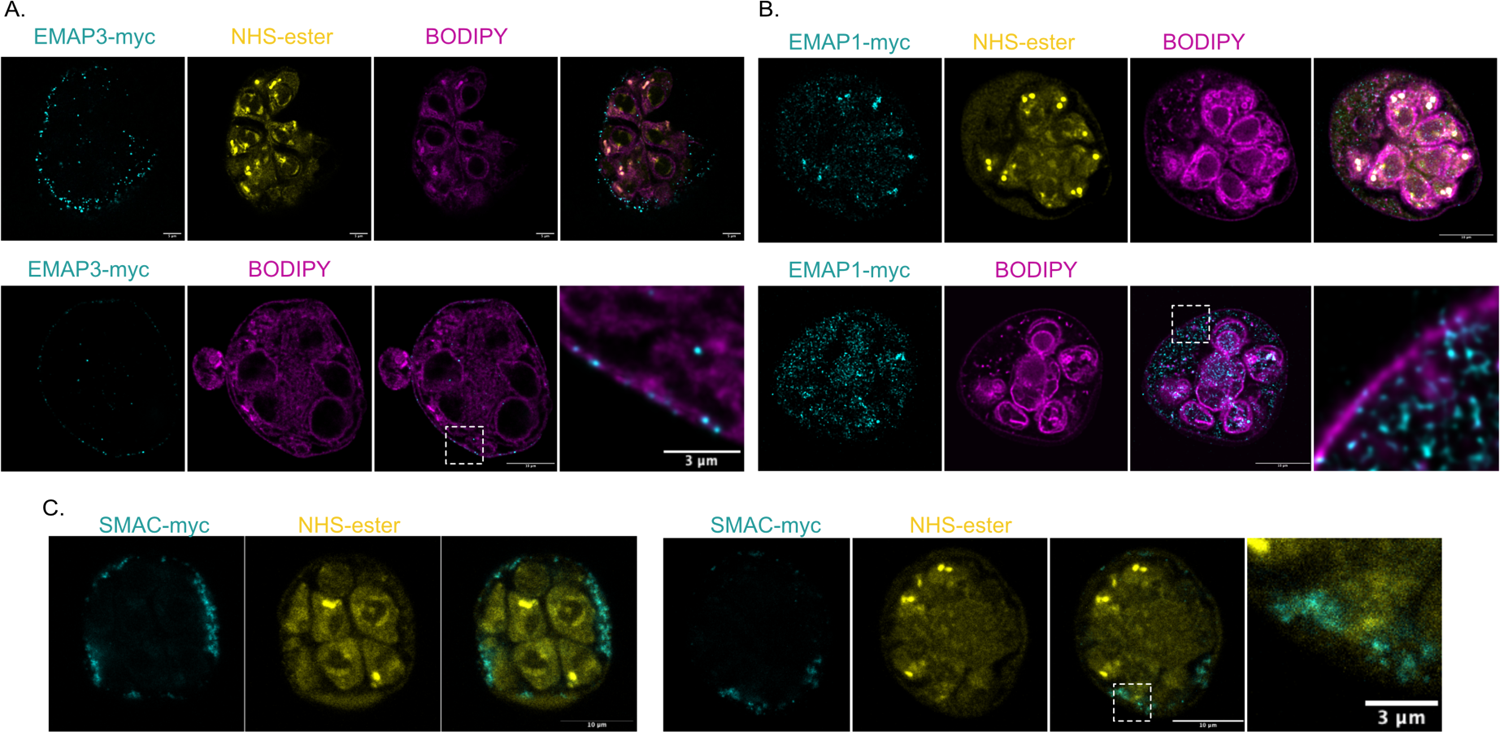
Ultrastructure expansion microscopy showing EMAP3 localisation to the iRBC membrane. Ultrastructure expansion microscopy (UEx-M) of **(A)** EMAP3-myc and **(B)** EMAP1-myc in schizonts with the c-myc tagged proteins in cyan with BODIPY ceramide to visualise membranes (magenta) and NHS-ester marking protein dense regions (yellow). **(C)** UEx-M of SMAC-myc schizonts with c-myc tagged proteins in cyan and NHS-ester stain in yellow. Scale bar = 10 µm or 3 µm. The images are maximum intensity Z-projections of three to four slices.

### EMAP3 interacts with EMAP1 but not with SMAC at the iRBC membrane

To determine putative parasite proteins that interact with the EMAPs and SMAC and to identify novel exported proteins we performed cross-linking co-immunoprecipitation (Co-IP) assays using antibodies against the 3x-myc tagged proteins. We performed Co-IPs of iRBC containing schizonts without and with saponin treatment of the iRBC. Proteins that are soluble in the iRBC will be lost with saponin treatment but those that are bound to the iRBC membrane or cytoskeleton will be retained. The saponin treatment is expected to remove soluble exported proteins and weak interactors of the myc-tagged bait proteins. A list of putative interacting proteins was generated for each 3x-myc tagged protein and visualised in heatmaps. Only proteins with at least four times the peptide abundance compared to the control, in at least two Co-IP replicates (under the same conditions), were included (**Table S1**, Fig. 3A). The control used was schizonts prepared from a PbCas9-FLAG_Tir1-myc mutant expressing the non-exported, 3x-myc tagged TiR1 (transport inhibitor response 1) protein, referred to as Tir1-myc. Correct insertion and expression of the 3x-myc tagged Tir1 protein was confirmed by diagnostic PCR-genotyping and Western blot analysis (**Fig. S1**).

**Figure 3.**
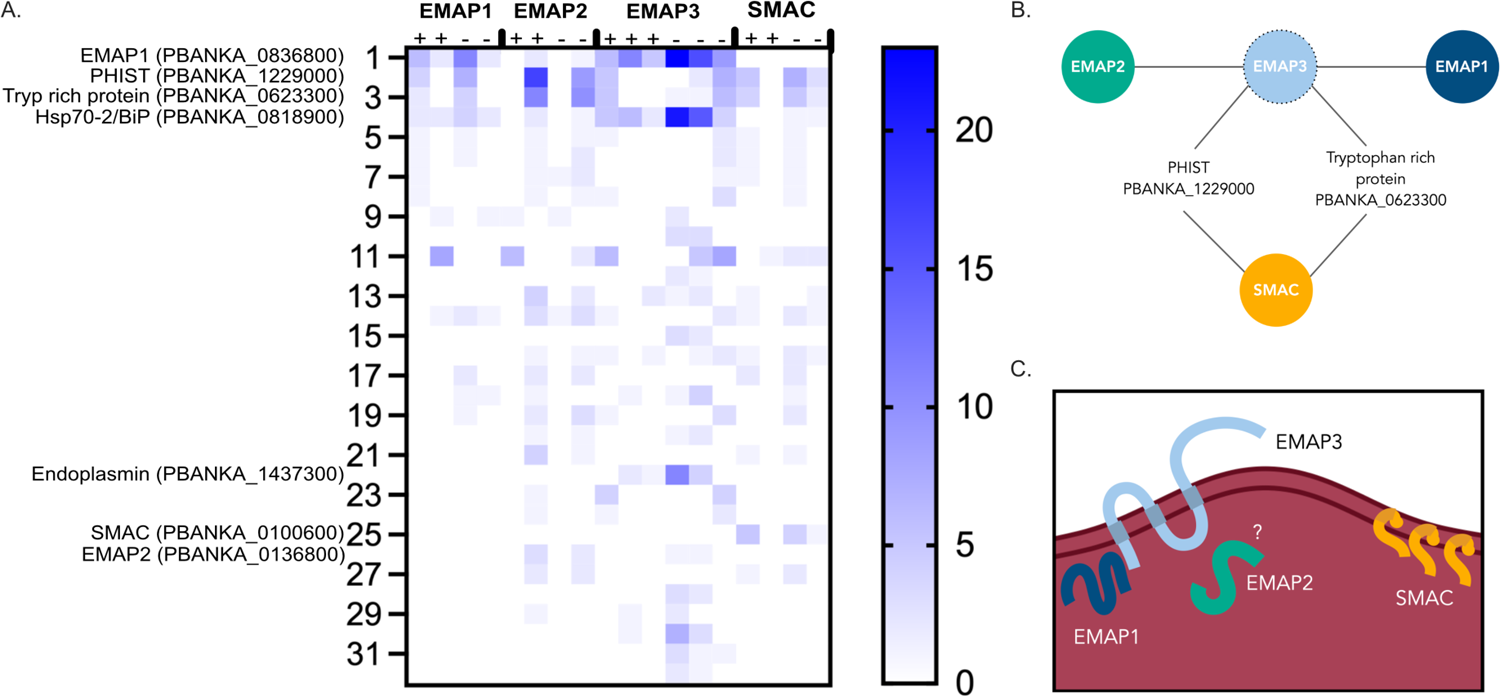
EMAP3 interacts with EMAP1 at the iRBC membrane but not with SMAC. **(A)** Heatmap summary of results from co-immunoprecipitation (Co-IP) using EMAP1-myc, EMAP2-myc, EMAP3-myc or SMAC-myc as bait proteins and analysed by nanoscale liquid chromatography coupled to tandem mass spectrometry (nano LC-MS/MS). Co-IPs were performed with (+) and without (-) saponin treatment followed by formaldehyde cross-linking. Results are shown only for proteins detected in a minimum of two replicas (under same conditions) at abundances a minimum of four-fold above TIR-myc control. **(B)** Schematic summary of selected shared interactions between the EMAPs and SMAC at the iRBC membrane, where only proteins detected in a minimum of two replicas are shown as solid lines indicating their interaction. EMAP3 has a dotted line indicating its absence from mass spectrometry detection. **(C)** Proposed model of orientation and interactions of EMAP3 at the iRBC membrane.

Western blot analysis of the Co-IP samples prior to mass spectrometry analysis confirmed the presence of the myc-tagged EMAPs and SMAC bait proteins with the expected sizes (**Fig. S3**). EMAP3-myc robustly co-immunoprecipitates EMAP1 in all biological replicates, with stronger interaction observed for the samples without saponin lysis. EMAP3-myc also co-precipitates EMAP2 but this interaction is detected less robustly (in two out of three replicas for non-saponin treated samples only). In contrast, there was no evidence of EMAP3-myc interacting with SMAC (Fig. 3A**, Table S1**). Next, we performed Co-IPs using EMAP1-myc, EMAP2-myc and SMAC-myc as bait proteins. The EMAP2-myc Co-IPs show evidence that EMAP2 interacts with EMAP1 (but not in all replicas), but EMAP2 was not present in the reciprocal Co-IP of EMAP1-myc. This is possibly due to a low abundance of EMAP2, as is evident from the low amount of EMAP2 peptide hits in the EMAP2-myc Co-IPs, and therefore the possible interaction between EMAP1 and EMAP2 remains uncertain. Unexpectedly, EMAP3 is not detectable in any of the Co-IPs, including the EMAP3-myc Co-IP. It is possible that EMAP3 is not easily detectable by mass spectrometry, which would explain its absence in the EMAP1-myc and EMAP2-myc co-IPs. In agreement with EMAP3-myc Co-IPs, SMAC was not identified in the EMAP1-myc or EMAP2-myc Co-IPs, which suggests that SMAC is not interacting with any of the EMAPs (Fig. 3A**, Table S1**).

Overall, only few proteins were detected above the set thresholds in the EMAPs and SMAC Co-IPs. This may indicate low numbers of interaction proteins or may result from the approach used where we combined cross-linking with RIPA solubilisation. For this reason, we also performed EMAP3-myc Co-IPs without crosslinker and lysed the samples with NP-40 detergent. While the interaction with EMAP1 was confirmed, only a few additional interactions were identified (**Fig. S4**). All EMAPs and SMAC show evidence of interacting with two shared proteins, a tryptophan-rich protein (PBANKA_0623300) and a *Plasmodium* helical interspersed subtelomeric (PHIST) protein (PBANKA_122900). In addition, all bait proteins, but in particularly EMAP1-myc and EMAP3-myc, also co-precipitate BiP (PBANKA_0818900), which is the ER heat shock protein 70 (HSP70) chaperone, also known as HSP70-2. In addition, EMAP3-myc co-precipitates PBANKA_1437300, another ER resident heat shock protein 90 (HSP90), also named endoplasmin or glucose response protein 94 (GRP94)^27^. These interactions likely take place in the ER and might indicate that EMAP3 is actively exported also in schizonts. The shared interactions between the EMAPs and SMAC at the iRBC membrane are summarised in a schematic figure (**Fig 3B**).

Furthermore, we investigated the mouse host proteins that were present in the EMAP3-myc and SMAC-myc crosslinked Co-IPs. We found that both EMAP3 and SMAC appear to interact with the mouse RBC membrane skeleton protein ankyrin-1 as well as spectrin (alpha and beta chain) and Band 3 protein, which link the RBC membrane to the membrane skeleton via ankyrin^28^, (**Fig. S4, Table S1**). These interactions seem specific to SMAC and EMAP3 and most likely not resulting from contamination due to their sheer abundance since they are absent or present at very low levels in Co-IPs with the non-exported Tir1-myc control (**Fig. S4**). In addition, EMAP3-myc but not SMAC-myc was found to bind mouse IgG immunoglobulin (peptide hits mapping to gamma (heavy) and kappa (light) chain variable and constant regions). This again is consistent with an iRBC surface exposure of EMAP3 where it would be exposed to mouse IgG antibody. EMAP3-myc also co-precipitated Protein 4.2 which is part of the RBC cytoskeleton. In summary, the data presented here indicates that EMAP3 interacts with EMAP1 at the iRBC membrane where it likely also interacts with EMAP2 and the RBC membrane skeleton. Based on the Co-IP interaction data, the three transmembrane domains of EMAP3 and the demonstration that the C-terminus of EMAP3 is exposed on the surface of the iRBC, we suggest a model of the interactions and localisation of the EMAP3 at the iRBC surface (Fig. 3C).

### EMAP3 neither mediates organ sequestration nor influences virulence

Based on the timing of expression and its location on the surface of the iRBC, we hypothesised that EMAP3 could mediate cytoadherence and tissue sequestration of the iRBC. To test this hypothesis we generated a gene-deletion mutant (*emap3* ko) lacking expression of EMAP3 using standard genetic modification technology^25^. The *emap3* gene was deleted in a dual reporter *P. berghei* (ANKA) reference line that constitutively expresses mCherry and luciferase, driven by the *hsp70* and *eef1a* promoter, respectively^29^. The dual reporter line hsp70:mcherry-eef1a:luciferase (mCherry-Luc) was used as a control in all experiments. Synchronised blood-stage infections were established in Balb/c mice by intravenous injection of histodenz-purified schizonts of the *emap3* ko or mCherry-luc line. The blood stage development of *emap3* ko parasites in the mice was monitored using Giemsa-stained smears of tail blood. The blood stage growth rate of *emap3* ko parasites was comparable with mCherry-Luc blood stages. Although *emap3* ko parasites appear to have a mildly reduced growth rate during the later days of the blood stage infection compared to the mCherry-Luc control, this reduction was not statistically significant (p-value = 0.183467), (Fig. 4A).

**Figure 4.**
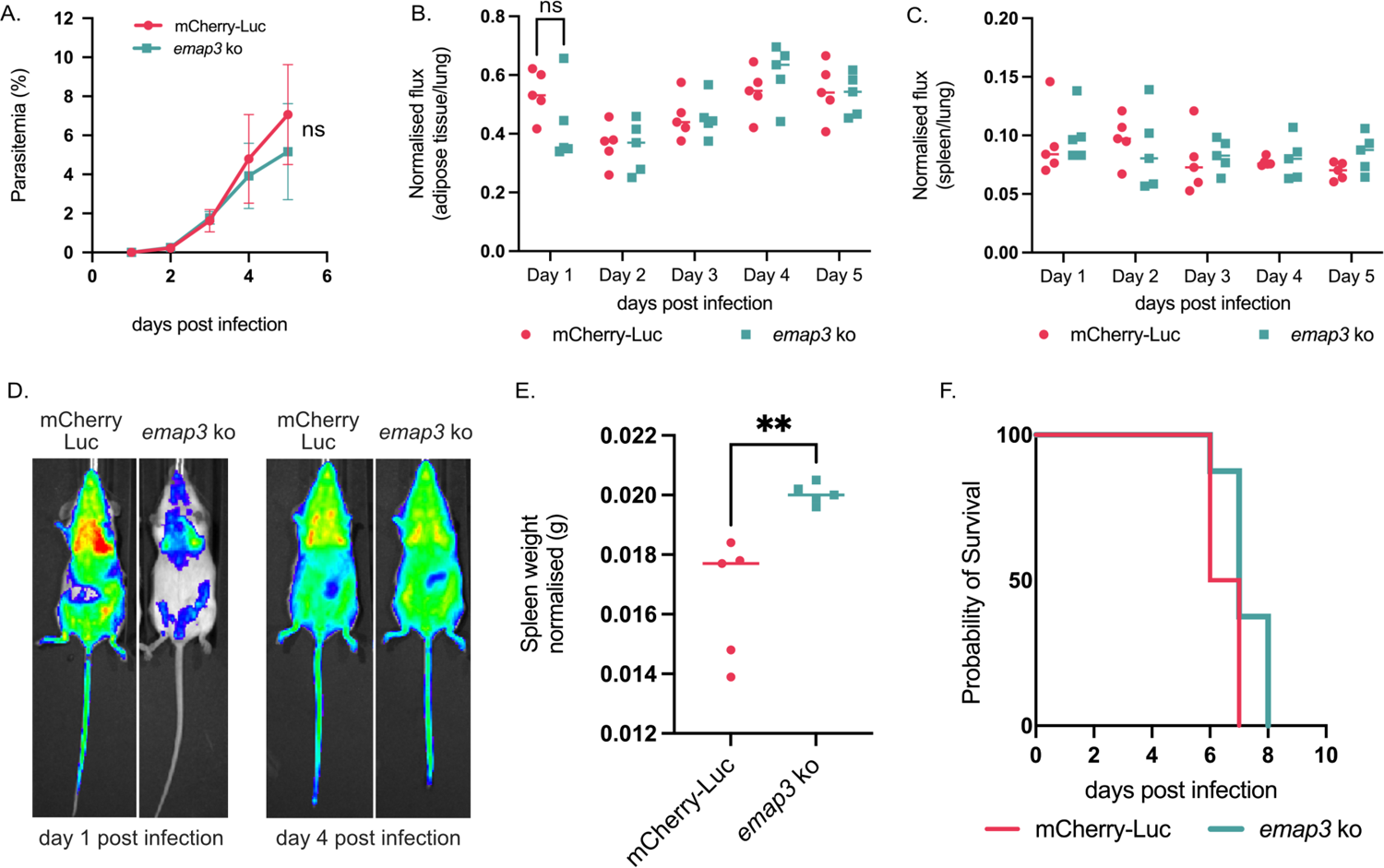
EMAP3 does not play a critical role in parasite sequestration and virulence. **(A)** *In vivo* blood stage growth assay of *emap3* ko compared to the mCherry-Luc *P. berghei* background line. Balb/c mice were injected with 1 x 10^6^ parasites intravenously (IV) on day 0. Parasitaemia was determined by Giemsa-stained smears where each day in the graph represents the average of eight Balb/c mice from two independent biological experiments. Day five Student’s t-test p-value = 0.183467. **(B-C)** Whole body imaging of *emap3* ko and mCherry-Luc parasites in Balb/c mice using the Spectrum *in vivo* imaging system (IVIS) to visualise luciferase expressing parasites in (B) adipose tissue and (C) spleen on day one to five after injecting 10-30 x 10^6^ purified schizonts IV. Day one Student’s t-test on adipose tissue signal p-value= 0.162349. Bioluminesence signal in each organ is normalised to the signal in lungs and expressed as normalised flux. Representative experiment with five Balb/c mice per parasite line is shown in (B-C) together with **(D)** representative whole-body images from day one and four of infection. Analysed IVIS data for a second independent experiment together with all IVIS images are available in **Fig. S5. (E)** Spleen weight of Balb/c mice infected with *emap3* ko and mCherry-Luc parasites on day five of infection where spleen weight is normalised to mouse body weight. Student’s t-test p-value = 0.0050. **(F)** Evaluation of virulence of *emap3* ko and mCherry-Luc parasites in C57BL/6 mice, expressed as probability of mouse survival where mice are scored for malaria symptoms and culled when they reach moderate symptoms. Eight (*emap3* ko) or ten (mCherry-Luc) C57BL/6 mice were assayed in two independent experiments. Symptom scoring data is available together with parasitaemia and spleen weight for C57BL/6 infections in **Fig. S6**.

In parallel, we investigated whether EMAP3 plays a role in tissue sequestration by performing whole-body *in vivo* imaging of the infected mice to measure parasite-expressed luciferase as a measure of parasite load in organs using the Spectrum *in vivo* imaging system (IVIS), (Fig. 4B-D, **Fig. S5**). As expected, in all *emap3* ko and mCherry-Luc infected mice the lungs show the highest parasite load, which is a major site of parasite accumulation along with abdominal adipose tissue^6^. During early stages of infection (day one post-infection), the *emap3* ko infected mice displayed a mildly reduced luciferase signal in adipose tissue, compared to the mCherry-Luc infected mice, however this difference was not statistically significant (p-value = 0.162349), and this small difference disappears as the infection progresses (Fig. 4B). It has been shown previously that *P. berghei* sequestration deficient blood stages commonly display reduced sequestration in adipose tissue but an increased luciferase signal in the spleen as a result of splenic clearance of the non-sequestering iRBC^17^. We did not observe an increase in luciferase signal in the spleen of *emap3* ko infected mice (Fig. 4C). However, the weight of isolated spleen from sacrificed *emap3* ko infected mice was significantly increased compared to spleens of mCherry-Luc infected mice (p-value = 0.0050), (Fig. 4E). Consistent with our observations that EMAP3 does not play a significant role in blood stage growth or sequestration in Balb/c mice, we observed no difference between emap3 ko and mCherry-Luc infected C57BL/6 mice with respect to virulence in mice. The *emap3* ko and mCherry-Luc infected mice developed moderate symptoms (as determined by scoring piloerection, mobility and hunching) at a comparable time (day six to eight post-infection), at which point they were culled (Fig. 4F**, Fig. S6**). The growth rate of *emap3* ko blood stages in C57BL/6 mice was not significantly lower compared to mCherry-Luc blood stages, and thereby mirrored results obtained in Balb/c infected mice (**Fig. S6**). Curiously no difference in spleen weight was observed between *emap3* ko and mCherry-Luc infected C57BL/6 mice (**Fig. S6**). We thereby conclude that EMAP3 does not play a significant role in parasite growth or sequestration and does not influence parasite virulence characteristics.

## Discussion

We here identify EMAP3, a novel *P. berghei* protein that is exported by blood stage parasites into the iRBC and transported to the iRBC membrane where its C-terminal is exposed on the outer side of the iRBC membrane. So far, the location of relatively few putative *P. berghei* exported proteins has been analysed. This includes proteins encoded by the multigene families *pir*, *fam-a*, *fam-b*, *etramp* (early transcribed membrane protein) and *phist* (*Plasmodium* helical interspersed subtelomeric), and the tryptophan-rich protein family ^8,20,21,30,31^. It was proposed that proteins encoded by the *pir* multigene family are located at the iRBC surface^32–34^, like the PfEMP1 protein of *P. falciparum* encoded by the *var* multigene family^4^. However, several recent studies analysing multiple different PIR proteins by tagging with fluorescent markers demonstrate a location in the iRBC cytoplasm and no evidence was found for a location at the surface membrane^5,21,22^.

Several proteins encoded by single copy genes including IBIS1 (intra-erythrocytic *P. berghei*-induced structures 1), SBP1 (skeleton binding protein 1), MAHRP1a (membrane-associated histidine-rich protein 1a), SMAC and exported proteins lacking functional annotation have also experimentally been shown to be exported into the iRBC. However, these proteins have either a diffuse location pattern in the iRBC cytoplasm or have a discrete punctate distribution associated with IBIS membranous structures^5,8,17^. For two *P. berghei* exported proteins, EMAP1 and EMAP2, a location at the iRBC surface membrane has been shown. Nevertheless, no strong evidence was found for exposure of these proteins at the outer side of the iRBC surface ^8^. To our knowledge, only one other protein, the merozoite surface protein PbTiP (*P. berghei* T-cell immunomodulatory protein) has been demonstrated to be exported to the iRBC membrane and exposed on the outer surface of the iRBC during asexual development ^35^.

For the EMAP3 protein we describe here, we demonstrate an outer surface location and provide evidence for an interaction with the two other iRBC surface membrane located *P. berghei* proteins, EMAP1 and EMAP2^8^. We did not find direct evidence for interaction with the exported protein SMAC that is located to the iRBC cytoplasm and mediates *P. berghei* iRBC sequestration^5,8^. Deletion of EMAP3, has no significant effect on blood stage development, sequestration of schizonts nor virulence. This is similar to the absence of an effect on blood stage development or iRBC sequestration of mutants lacking either EMAP1 or EMAP2^8^. We therefore conclude that EMAP3 is not a major mediator of iRBC sequestration and is thereby not the so far unidentified parasite surface ligand that binds to CD36.

EMAP3-myc reproducibly co-precipitated EMAP1 in all Co-IP conditions tested, and to a lesser extent also EMAP2. We performed reciprocal Co-IPs but EMAP3 was not detected neither in EMAP1-myc or EMAP2-myc Co-IPs, nor when EMAP3-myc itself was used as the bait. This is despite the presence of residues for tryptic digestion in the EMAP3-myc amino acid sequence, confirmation of EMAP3-myc in the Co-IP lysates by Western blot and previous report of its detection by mass spec^36^. Two proteins showed evidence of interacting with the three EMAPs (in particular EMAP3) and SMAC. These were tryptophan-rich protein (PBANKA_0623300) and a PHIST protein (PBANKA_122900). The PHIST protein PBANKA_122900 has previously been identified by proteomics, found to be dispensable to blood stage growth and to localise to IBIS in the iRBC^8,30^. There is no knockout phenotype or localisation data available for the tryptophan-rich protein PBANKA_0623300, however it was previously detected in a proteomic study as a putative liver merosome protein^36^.

The role of SMAC in *P. berghei* iRBC sequestration and the observation that SMAC-myc localises to dense and large patches in the iRBC cytoplasm supports the hypothesis that SMAC might serve a trafficking role to present a CD36-binding parasite ligand on the surface of the iRBC^5^. SMAC does not interact with any of the EMAPs and is therefore unlikely to play a role in anchoring EMAP3 at the iRBC membrane. Instead SMAC likely plays a role in presenting a yet to be identified parasite ligand binding to host CD36 that was not captured in our Co-IP set up. There was overall a paucity in interactions identified and relatively few interaction partners were identified for all bait proteins, and in particular for SMAC-myc. This indicates that conditions we have used in our Co-IPs might have enabled detection of only the most robust or abundant interactions such as those between EMAP3 and EMAP1. Further efforts to optimise Co-IP conditions might identify more SMAC-interacting proteins, including a potential *P. berghei* surface ligand that binds to host CD36.

The lack of direct evidence for EMAP3 interacting with SMAC indicates that EMAP3 is trafficked onto the iRBC surface in a mechanism independent of SMAC. Analysis of transport and location of EMAP3 in a mutant lacking expression of SMAC would conclusively prove this. Interestingly, it has been shown that EMAP1 and EMAP2 are still expressed at the iRBC surface in the absence of SMAC^8^, which supports the hypothesis that EMAP3 is also transported to the iRBC surface membrane independently of SMAC. These observations now pave the way for the generation of *P. berghei* mutants where the CD36-binding of iRBC is abolished by deletion of the *smac* gene and where EMAP3 is used as a scaffold presenting *P. falciparum* proteins on the outside surface of the iRBC by tagging them to EMAP3. For example, EMAP3 could be tagged in these mutants with cytoadherence proteins or domains including PfEMP1 variants. The dispensable nature of *emap3* for blood stage growth is an advantage as it indicates that is likely possible to truncate the endogenous *emap3* gene after the last transmembrane domain to attach the *P. falciparum* proteins/domains onto the iRBC surface.

EMAP1 has previously been used as the scaffold to display Var2CSA on the surface of *P. berghei* infected iRBC as a model to study pregnancy associated malaria experimentally *in vivo*. However, only a very low proportion (<6%) of iRBC were found to express EMAP1-Var2CSA at the outer side of the iRBC^37^. This is in agreement with previous observations that only a low percentage (around 7%) of iRBC exposed EMAP1 on the their outside^8^ and with our observations of the lack of evidence for the C-terminus of EMAP1-myc being exposed on the outer surface of the iRBC. With no predicted transmembrane domain in EMAP1, we propose that the association of EMAP1 with the RBC-membrane is by its interaction with the N-terminus of the transmembrane protein EMAP3. An improved platform, where EMAP3 is used as an expression scaffold in a mutant lacking SMAC expression, could be combined with humanised mice expressing for example human CD36^38^ offering a much needed platform to test iRBC sequestration and disease modulating therapies aimed at interrupting *P. falciparum* cytoadherence and sequestration *in vivo*.

## Materials and Methods

All antibodies, key reagents and primer sequences are available in **Table S1**.

### Identification of *emap3 in silico*

To identify *Plasmodium* exported proteins, we applied previously reported selection criteria of genes for fluorescent tagging based on existing data for transcription and/or protein expression in *P. berghei* blood-stage parasites and presence of protein export motifs^8,21^. By refining these selection criteria for expression in both *P. berghei* blood- and liver-stage parasites^39^ and annotation as conserved *Plasmodium* protein, with unknown function we identified *emap3* (PBANKA_0825900).

### Animal Work

The parasitaemia of infected animals was determined by microscopy of methanol-fixed and Giemsa-stained thin blood smears from small-volume blood samples. Infected blood was collected by heart puncture on terminally anaesthetised mice and collecting the blood into a syringe containing heparin (Sigma-Aldrich).

### Animal work at Umeå University

Animal experiments were performed under and according to ethics permits A34-2018 and A24-2023 as approved by the Swedish Board of Agriculture (Jordbruksverket). For routine propagation of parasites BALB/c mice were used and for pathology experiments C57BL/6 mice. Female-only mice were purchased from Charles River Europe and used for experiments from six weeks of age. Housing was in groups of four mice in individually ventilated cages (IVC) with autoclaved wood chips and nesting material and a temperature of 21 ± 1 °C under a 12:12 h light-dark cycle with a relative humidity of 55 ± 5%. Specific pathogen-free conditions were maintained and biannual Exhaust Air Dust (EAD) monitoring and analysis was performed. Fresh water and commercial dry rodent diet was freely available (*ad libitum*) at all times. Visual health checks were performed daily to monitor animal health.

### Animal work at Leiden University Medical Centre

All animal experiments were approved and performed under a license given by competent authority after ethical evaluation by the Animal Experiments Committee Leiden (AVD1160020171625). All Animal experiments were carried out in agreement with the Experiments on Animals Act (Wod, 2014), which is the applicable legislation in The Netherlands that follows European guidelines (EU directive no. 2010/63/EU). All animal experiments were performed in Leiden University Medical Center; LUMC, a licensed establishment for animal experimentation. Experiments were performed with female 6-7 week old OF1 mice, which were purchased from Charles River Laboratories. Mice were housed in ventilated cages supplied with autoclaved aspen wood chips, wood chew block, fun tunnel and nestlets (at 21±2°C; 12-h light–dark cycle; relative humidity of 55±10%). Mice were fed and supplied with water *ad libitum*. They were fed commercially available autoclaved dry rodent diet pellets.

### Animal work at University of Geneva

Animal experiments were conducted with the authorisation numbers GE102 and GE-58-19, according to the guidelines and regulations issued by the Swiss Federal Veterinary Office. *P. berghei ANKA* strain clone 2.34 together with derived transgenic lines, were grown and maintained in CD1 outbred mice. Six to twelve-week-old mice were obtained from Charles River laboratories, and females were used for all experiments. Mice were specific pathogen free (including *Mycoplasma pulmonis*) and subjected to regular pathogen monitoring by sentinel screening. They were housed in individually ventilated cages furnished with a cardboard mouse house, tunnel and Nestlet, maintained at 21 ± 2°C under a 12 hours light/dark cycle, and given commercially prepared autoclaved dry rodent diet and water *ad libitum*.

### Plasmids and parasite lines

The *em*ap3 KO (PBANKA_0825900) plasmid was generated by amplifying 0,5 kb (5’) and (3’) homology arms flanking the *emap3* open reading frame (ORF) from *P. bergehi* genomic DNA and subsequently cloned into the pL0001 plasmid (www.mr4.com) carrying the mutated *tgdhfr/ts* (*Toxoplasma gondii* dihydrofolate reductase-thymidylate synthase) selection marker expressed under control of the *pbdhfr/ts* (*P. berghei* dihydrofolate reductase-thymidylate synthase) 5’ and 3’ UTRs (untranslated regions). The resulting *emap3* KO pL2296 plasmid was linearised (using KpnI, NotI, ScaI) and transfected into the *P. berghei* mCherry.Luc background line (hsp70p:mCherry eef1ap:Luciferase, 1868Cl1)^29^ and cloned by standard limited dilution cloning^25^ to generate the PbEMAP3 KO line (3076cl1).

The emap3-myc (PBANKA_0825900) tagging vector was generated by amplifying a 0,7 kb 3’ fragment of *emap3* that was cloned in-frame with the coding sequence for 3x-myc, which upon integration of the vector into the *emap3* target locus inserts a C-terminal 3x-myc tag. To this end the homology arm fragment was amplified from *P. berghei* genomic DNA and cloned into the the pL1672 vector^40^ containing the selection marker *tgdhfr/ts* using BamHI/EcoRI restriction enzymes to generate the emap3-myc vector (pL2302), which was linearised with NdeI prior to transfection. The emap1-myc (PBANKA_0836800, pL1594), emap2-myc (PBANKA_0316800, pL2358) and smac-myc (PBANKA_0100600, pL1435) tagging vectors were generated by replacing the mCherry tag with a 3xMyc tag in existing C-terminal tagging vectors carrying the *tgdhfr/ts* selection marker^5,8^. The vectors were linearised prior to transfection using NdeI (pL1594), ClaI (pL2358) or NdeI (pL1435). The resulting 3x-myc tagging vectors were transfected either into PbANKA ANKA cl15cy1 reference line^41^, (emap1-myc, line 1583 and emap3-myc, line 3084), mCherry.Luc (emap2-myc, line 3296) or GFP.Luc^6^, (smac-myc, line 1413).

The PbCas9-FLAG_Tir1-myc line was generated by cloning the auxin inducible *osTir1* (transport inhibitor response 1 gene from *Oryza sativa*)^42^, fused to a C-terminal 3x-myc tag and flanking it by 260 bp (5’) and 288 bp (3’) homology arms targeting insertion into the p230p locus, downstream of the GIMO locus^43^. The 5’ p230p and 3’ p230p targeting sequences were cloned into the pPbU6-hdhfr/yfcu plasmid (Addgene #216422),^44^ carrying the dual positive and negative selection marker *hdhfr-yfcu* (human dihydrofolate reductase/yeast cytosine deaminase and uridyl phosphoribosyl transferase) under control of *pbeef1a* (*P. berghei* elongation factor alpha) 5’UTR and *pbdhfr/ts* 3’UTR. The selection marker was made recyclable by addition of an additional *pbdhfr/ts* 3’UTR motif upstream of the *pbeef1a* 5’UTR. Finally a gRNA targeting p230 were designed using EupaGDT and ligated using T4 ligase (New England Biolabs, Ipswich, MA, USA), resulting in the pYCs-PbU6-hDHFR/yFCU-NS-p230p plasmid. The *eef1a* promoter used to drive osTir1 was amplified from pYCs plasmid^45^ and the *P. falciparum* histidine rich protein 2 (*pfHrp2)* terminator was amplified from pDC2-Cam-Cas9-hDHFR^46^. The *eef1a* promoter, osTir1-3x-myc, and *hrp2* terminator were seamlessly ligated into pSKA-001 vector using NEBuilder HiFi DNA Assembly Master Mix (NEB) and the intermediate fragment was cloned into pYCs-PbU6-hDHFR/yFCU-NS-p230p vector. The resulting vector was transfected into the PbCas9 line where spCas9 (*Streptococcus pyogenes* CRISPR associated protein 9) is integrated in the GIMO (gene in marker out) locus^43,44^.

### Transfections

The parasite lines described above were generated by transfection of *P. berghei* schizonts and transgenic parasites were selected for by pyrimethamine, all as previously described^25^. In brief, *P. berghei* parasites were cultured *ex vivo* from infected blood (approximately 1-5% parasitaemia) at 37^о^C with 120-150 rpm shaking for 22 hours. The culture was then checked by Giemsa staining for the presence of schizonts. Schizonts were harvested by using a 15.2% w/v Histodenz gradient (Sigma-Aldrich), (from 27.6% w/v Histodenz buffered stock solution), collecting the brown parasite layer on the interface. Collected schizonts were washed and then re-suspended in P3 buffer (Lonza) and mixed with DNA. The schizont-DNA mixture was then loaded onto a Lonza transfection cuvette and electroporated. Fresh media is added to the cuvette and aspirated for direct intravenous injection into a mouse. Mice are treated with pyrimethamine in drinking water a day after transfection to select for transgenic parasites. The transgenic lines obtained were genotyped by diagnostic PCR with all genotyping primers available in **Table S1**. DNA for genotyping was extracted from whole blood (>1% parasitemia) typically using the Qiagen DNAeasy or GeneJet Genomic DNA purification kit (Thermofisher).

### Immunofluorescence Assays

Immunofluorescence assays (IFA) were carried out by allowing 20-100 µL of cells fixed with 4% paraformaldehyde (diluted in RPMI or PBS) to settle on poly-D-lysine (GIBCO) coated slides for 15 minutes. The cell suspension was then removed and the slide was washed with 200 µL of 1X PBS. To permeabilize cells, 200 µL of 0.2% Triton X-100 was added and allowed to incubate for 5 minutes. The cells were then washed 3 times with 200 µL of 1X PBS. Blocking buffer was then added (200 µL of, 3% BSA in 1X PBS) and incubated at room temperature (RT) for 30-40 minutes. The blocking solution was removed and 100 µL of primary antibody solution (diluted in blocking buffer) was added to the slide that was placed in a humid chamber and allowed to incubate at 4^о^C overnight. The primary antibody was then removed and the slide was washed three times with 1X PBS before adding the secondary antibody (diluted in blocking buffer). The slides were incubated in a dark humid chamber for 1-2 hrs at RT. The secondary antibody was removed and the slides were washed three times with 200 µL of 1X PBS. The cells were mounted with 5 µL of Vectashield with DAPI and covered by a sealed coverslip. Slides were allowed to dry overnight before imaging. Widefield images were taken using a Zeiss Apotome microscope at 63X objective and confocal images were taken using a Leica SP8 microscope at 63X objective.

### Surface shaving and permeabilisation

Surface shaving and permeabilisation was performed by adding 200 µL of ice-cold 0.003% trypsin to 50 µL of gradient-purified schizonts. This was allowed to incubate at RT for 50 minutes. After incubation, the tube was spun at 500 g for 1 minute, the supernatant was removed and 200 µL of RPMI was added to stop trypsin activity. In parallel, cells with 200 µL of RPMI were allowed to incubate at RT for 50 minutes as a negative control for trypsin treatment. The aforementioned IFA protocol was then performed on the cells where the permeabilisation (0.2% Triton X-100) step was omitted for non-permeabilised set-ups.

### Brefeldin A treatment

Treatment with Brefeldin A (BFA, Sigma) was performed by taking infected mouse blood (mostly rings and early trophozoites) and setting up 3 mL *ex-vivo* cultures in 6-well plates. BFA was added to a final concentration of 5 µM (stock concentration 5 mg/mL). As a control, the same volume of DMSO was added to control wells. An aliquot (T=0) of this sample was taken and fixed with 4% PFA. The plate was placed in an airtight container following the candle jar method^47^ and incubated at 37^о^C. After three hours, an aliquot of the sample was taken and fixed with 4% PFA. The standard IFA protocol was then followed to determine the localisation of the 3x-myc tagged protein with and without BFA treatment.

### Ultrastructure Expansion Microscopy

Ultrastructure expansion microscopy (UEx-M) was performed on schizonts separated from uninfected erythrocytes on a density gradient (3.6 mL Percoll, Cytiva, 1 mL RPMI and 400 µL 10X PBS). Schizonts were fixed with 4% PFA according to protocols adapted to *Plasmodium* parasites^48^. Cells were allowed to settle on 12 mm round coverslips coated with poly-d-lysine for 10-15 minutes. The coverslips with cells were then incubated in a 1.4% formaldehyde (FA) and 2% acrylamide (AA) solution to add protein anchors. This was done by taking the coverslips and placing them in individual wells of a 24 well plate and adding the FA/AA solution, the plate was incubated at 37^о^C for 5 hours, water was added to the empty wells to prevent dehydration of the plate. The gel solution was prepared by adding 5 µL of 10% ammonium persulfate (APS) and 5 µL of 10% TEMED to an aliquot of monomer solution (23% Sodium acrylate; 10% AA; 0.1% BIS-AA in PBS). 35 µL of the gel solution was then placed on parafilm and the coverslips were inverted onto the gel (cell side down) to perform gelation. Coverslips embedded in gels were allowed to gelate for 1 hour at 37^о^C. Denaturation was performed by taking the gels and placing them in 1.5 mL tubes with denaturation solution. The tubes were then incubated at 95^о^C for 1 hour. Gel expansion was then performed by placing the gels in 200 mL of double distilled (ddH20) at RT for 30 minutes. This step was repeated two more times, replacing the water each time. The gels were then left to expand overnight at room temperature. The following day, the gels were shrunk by incubating them twice in 1X PBS for 15 minutes each time. The shrunk gels were transferred to 6 well plates and incubated in 1 mL of primary antibody solution at 37^о^C for 2 hrs with 125-150 rpm shaking. The gels were then washed three times with 1 mL of 0.1% PBS-Tween. The gels were then incubated in a secondary antibody solution at 37^о^C for 2 hrs with 125-150 rpm shaking, followed by three washes with 1 mL of 0.1% PBS-Tween. Gels were subsequently incubated in stains (NHS ester, Merck and BODIPY, Thermofisher) for 1 hr at RT shaking at 125-150 rpm. The gels were then washed three times with 1 mL of 0.1% PBS tween. Final gel expansion was performed by placing the gels in 200 mL of ddh20 and incubating for 30 minutes, this was repeated three times. The gels are allowed to incubate in ddh20 overnight in the dark. Finally the gels were cut into small cubes and placed on a poly-d-lysine coated 24mm coverslip in an O ring. The expanded cells were visualised on a confocal Leica SP8 microscope using 63X and 100X objectives.

### Growth Assays

To evaluate parasite blood stage growth mice were injected with 1×10^6^ parasites intravenously. Parasitemia was tracked by smearing daily starting three days after initial infection. Thin blood smears were made by small volume tail bleeds collected by lancet and smeared on glass slides. The smears were fixed with methanol and stained with 10% Giemsa for 10 minutes. The parasitemia was calculated by counting the number of *P. berghei* infected cells per 1000 RBC. The parasitemia over five days of infection of the *emap3* KO was compared to the mCherry-Luc background line.

### In vivo imaging

To evaluate parasite sequestration to organs, in vivo imaging (IVIS) was performed following the established protocol for *P. berghei* ^5,6^. In brief, to establish a roughly synchronous infection, mice were injected with 10-30 x 1×10^6^ Percoll-purified schizonts at 12 noon, intravenously. After 24 hrs (12 noon the next day), mice were anaesthetised with isoflurane and injected with 30 µL of 80 mg/mL D-luciferin (Synchem) subcutaneously. After 1-2 minutes of injecting D-luciferin, imaging with the IVIS Spectrum imager was performed. Mice were imaged at the same time (between 11-12 noon) each day, until day five post infection. On the last day of imaging, mice were anaesthetized and dissected to collect spleens for determination of spleen weight. IVIS images were analysed using the Living Image software, taking only images with no saturated pixels. Regions corresponding to the lungs, adipose tissue and spleen were drawn for each mouse using the region of interest (ROI) tool in the software.

### Co-immunoprecipitation

A formaldehyde cross-linking co-immunoprecipitation (Co-IP) protocol was performed to fix protein complexes together and co-precipitating any proteins interacting with the 3-myc tagged bait protein according to protocols adapted from established methods^49^. Co-IPs were performed on Percoll purified schizonts from a 100 mL schizont culture with 5 mL of infected mouse blood with a 1-5% parasitemia. The schizonts were harvested from the gradient and pelleted at 300 g for 3 minutes. The schizont pellet was washed two times with 1 mL of fresh RPMI. The pellet was then resuspended in 500 µL of RPMI, this was then split equally between two tubes. To one pellet, 500 µL of 0.1% ice cold saponin was added then incubated on ice for 5 minutes. The other pellet was kept untreated with saponin, instead 500 µL of RPMI was added. After saponin treatment, 500 µL of RPMI was added to stop the reaction. To crosslink the sample, formaldehyde was added to a final concentration of 1% and incubated at room temperature for 10 minutes with gentle agitation. To stop the crosslinking reaction, 10 mL of 0.125 M glycine was added to the samples and incubated at RT with gentle agitation for 5 minutes. The sample was spun at 1000 g for 10 minutes at 4^о^C and the supernatant was removed. The pellet was then used to perform the next IP steps on the same day or stored at −80^о^C for future use.

To immunoprecipitate 3x-myc tagged proteins, Protein G dynabeads (Thermofisher) were coated with rabbit α-myc antibody (Cell Signalling Technologies) following manufacturer’s protocol for coating beads for immunoprecipitation. In brief, magnetic beads (60 µL per sample) were placed on a magnet and allowed to separate from storage liquid for 10-30 seconds. The beads were then washed with 1 mL of 0.01% PBST. After washing, 7 µg of rabbit α-myc antibody was added. The beads were allowed to bind to the antibody for 15 minutes at room temperature with gentle rotation. To the cross-linked pellet, 1 mL of ice-cold RIPA lysis buffer was added. The tubes were then incubated on ice for 10 minutes. The pellet-RIPA solution was then passed through an insulin syringe 20x to solubilize the pellet. The sample was then spun at 14000 g for 15 minutes, 4^о^C. A 30 µL aliquot from this solution was saved as the ‘total protein’ sample. The supernatant was then mixed with 60 µL of the α-myc antibody coated magnetic beads and allowed to bind overnight at 4^о^C with gentle rotation. The bead-sample slurry was transferred to a fresh tube, then placed on the magnetic rack for 30 seconds to one minute. The liquid was collected as the ‘flow through (FT)’ sample. The tube was then separated from the magnetic rack and 1 mL of ice-cold RIPA was added to resuspend the beads. The tubes were placed back on the magnetic rack and the liquid was removed. The RIPA wash was repeated two times. After the last wash with RIPA, the beads were resuspended with 1 mL of ice cold 1X PBS with protease inhibitors and placed in a fresh tube. This was then allowed to settle on the magnet for 30 seconds to 1 minute. The wash with 1X PBS with protease inhibitors was repeated two times. The beads were resuspended in 500 µL of 1X PBS with protease inhibitors after the last wash. An aliquot of the beads was collected as the ‘immunoprecipitation (IP)’ sample and analysed by Western blot together with the total protein and flow-through samples.

Mass spectrometry data was analysed using the Scaffold software. A 1% protein false discovery rate (FDR) with at least 2 peptides identified per protein and a 0.1% peptide FDR was set as thresholds. To generate a list of putative protein interactions for each 3x-myc tagged *P. berghei* line (Table S1), only host and parasite proteins that were 4x more abundant in the samples compared to the Tir1-myc control were included.

### Western Blotting

Samples for western blotting were mixed with a 4X Laemmli buffer with β-mercaptoethanol (BME). These were incubated at 70^о^C for 10 minutes. Samples were run on either a 4-12% polyacrylamide pre-cast gel (NuPage, Thermofisher) or a 4-20% tris-glycine polyacrylamide gel (Mini-PROTEAN TGS, BioRad) at constant voltage (90-150V), until the running dye has reached the bottom of the gel. Transfer to a PVDF or nitrocellulose membrane was performed by activating the membrane in methanol then soaking in cold transfer buffer. The transfer stack was assembled following Thermofisher or Bio-Rad guidelines for Western blotting. Wet transfers were performed at 30V for 1 hr. The membrane was removed and stained with Ponceau S staining solution to visualise total protein amount and confirm successful transfer to membrane. The membrane was blocked in 5% Milk in 0.05% PBS-Tween for at least 1 hour on a shaker. The blocking solution was removed and primary antibody diluted in blocking solution was added three times with 0.05% PBS-Tween, 10-15 minutes each wash. The secondary antibody diluted in blocking solution was added and allowed to incubate for 2-3 hrs on shaking. The membrane was washed three times with 0.05% PBS-Tween, 10-15 minutes each wash. Detection solution was prepared (Amersham, ECL) and added to the blot, followed by imaging with the BioRad Chemidoc.

### Virulence assay

Virulence assays were performed by injecting 1×10^6^ parasites intravenously into six week old female C57BL/6 mice. Mice were health checked daily and given a symptom score following a scoring matrix. Briefly, a mouse was given a score from 1 (not affected) to 3 (significantly affected) for each category: piloerection (fur condition), movement and hunching (body posture). Infection was terminated when a mouse reached a score above six, which is equivalent to moderate symptoms, or scored six at two subsequent scoring occasions. Moderate symptoms equate to substantial piloerection, reduced mobility even under stimulation and persistent hunching. Symptoms of cerebral malaria were also checked for by testing reflexes, self-preservation, balance, motor skills and muscle tonus. Each day, the weight of the mouse was recorded, and thin blood smears collected to enumerate parasitemia.

## Figures

All figures and schematics were made and arranged using Affinity designer.

## Reagent availability

Plasmids and parasite lines are available upon request from Leiden University Medical Centre (contact).

## Acknowledgements

We acknowledge the Biochemical Imaging Center at Umeå University and the National Microscopy Infrastructure, NMI (VR-RFI 2019-00217) for providing assistance in microscopy. We also acknowledge the University of Geneva Proteomics Facility for providing assistance with proteomics and analysis of proteomic data.

## Author contributions

S.H. and R.R. planned experiments and analysed data. S.H., R.R., T.K.J., M.S.P., S. C-M., T.I. and M.R. conducted experiments. S.H. and E.S.C.B wrote the original draft. S.H. generated the figures. E.S.C.B, B.F.F. and C.J.J conceptualised the study. E.S.C.B., M.B., B.F.F. and C.J.J. supervised. All authors participated in the review and editing of the final manuscript.

