## Supplemental figures for "Erythrocyte membrane protein 3 (EMAP3) is exposed on the surface of the *Plasmodium berghei* infected red blood cell"

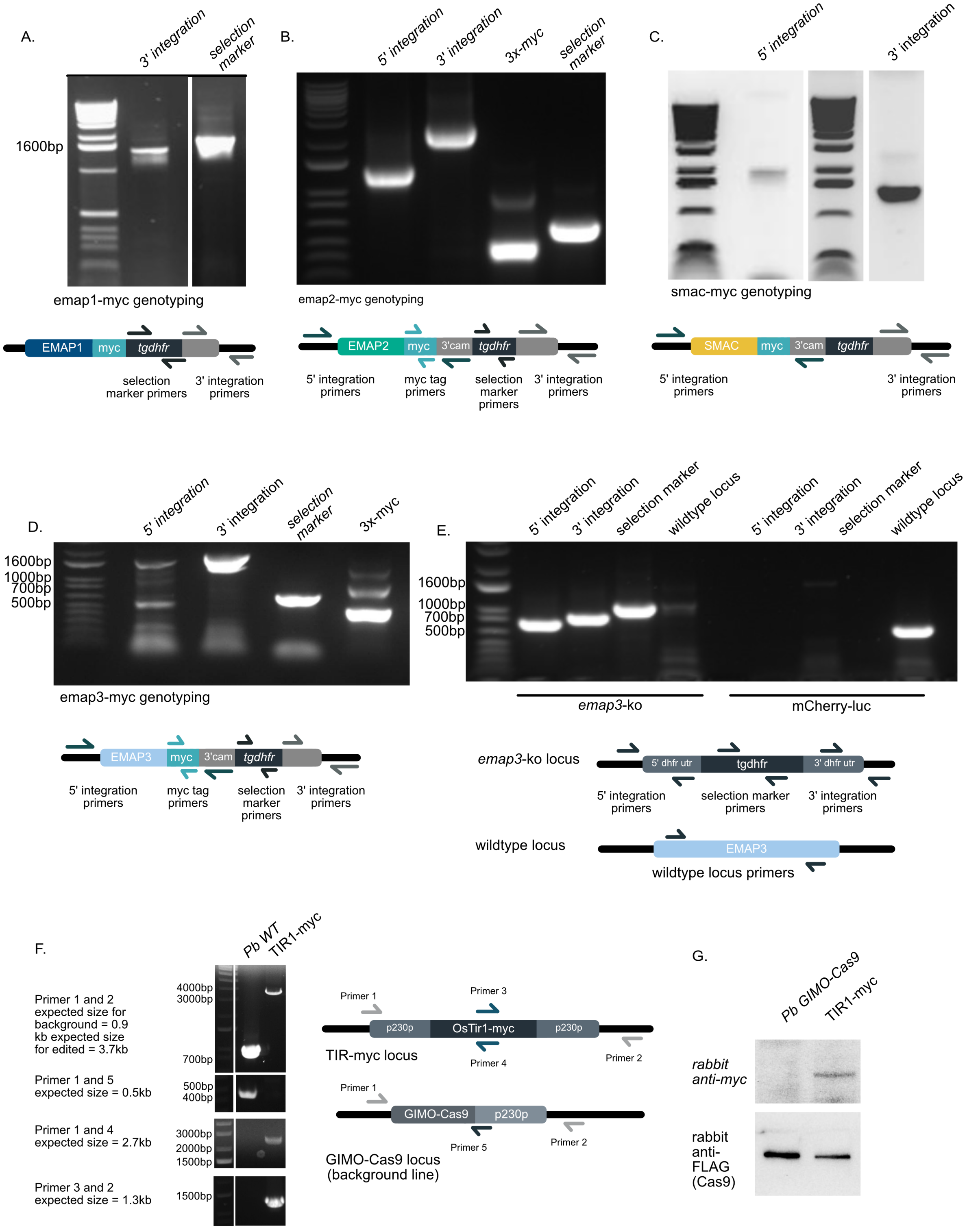


**Figure S1. Diagnostic genotyping of *P. berghei* myc-tagged and knockout (ko) lines. (A-D)** PCR-based genotyping of the (A) emap1-myc, (B) emap2-myc and (C) smac-myc and (D) emap3-myc tagged lines. To confirm 5' integration, primers annealing upstream of the homology region were used together with primers annealing within the calmodulin 3' untranslated region (UTR) used as terminator for the 3x-myc tagged gene. To confirm 3' integration, primers annealing downstream of the vector insertion site in the genome were used together with primers annealing within the vector backbone. The dihydrofolate reductase-thymidylate synthase (*tgdhfr/ts*) selection marker or 3x-myc tag were detected by primers specific to those sequences and included as additional controls. **(E)** To verify 5' and 3' integration of the knockout cassette in the *emap3 ko* line, primers that sit up- (5') or down- (3') stream of the homology arms were used together with primers annealing within the (*tgdhfr/ts*) selection marker. The presence of the selection marker in the knockout line was verified with primers amplifying the *tgdhfr/ts*. Primers against the wildtype *emap3* locus were used to verify the absence of wildtype in the emap3 *ko* line. **(F)** To verify integration of the TIR1-myc tagging cassette, primers that sit outside the homology regions were used together with primers that sit in the UTRs of the edited locus. Primers against the wildtype GIMO-Cas9 p230p locus were used to verify the absence of wildtype in the knockout line. **(G)** Western blot confirming the expression of TIR1-myc using a rabbit α-myc primary antibody. Expression of Cas9-FLAG using a rabbit α-flag antibody was used as a loading control.

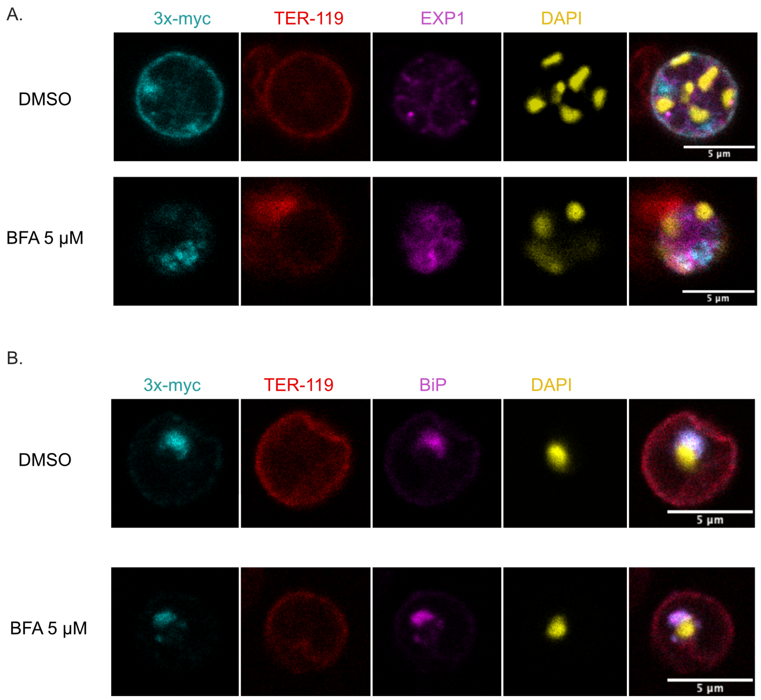


**Figure S2. Brefeldin A treatment of EMAP3-myc parasites.** Treatment of *P. berghei* EMAP3-myc mixed blood stage (rings and trophozoites) culture with 5µM Brefeldin A (BFA) was carried out for 3 hours. Staining with rabbit α-myc was performed to detect EMAP3-myc. An antibody against TER-119 was used to detect the RBC membrane. Exported protein 1 (EXP1) was used as a marker of the parasitophorous vacuole (PVM), binding immunoglobulin protein (BiP) as a marker for the ER and 4′,6-diamidino-2-phenylindole (DAPI) was used to stain DNA. Treatment with dimethylsulfoxide (DMSO) only was used as a control. The effect of BFA treatment on EMAP-3 localisation in (**A**) mature (schizont / trophozoite stage) and (**B**) young (ring stage) parasites were observed. No fully segmented, mature schizonts were detected in the BFA treated sample. Scale bar = 5 µm.

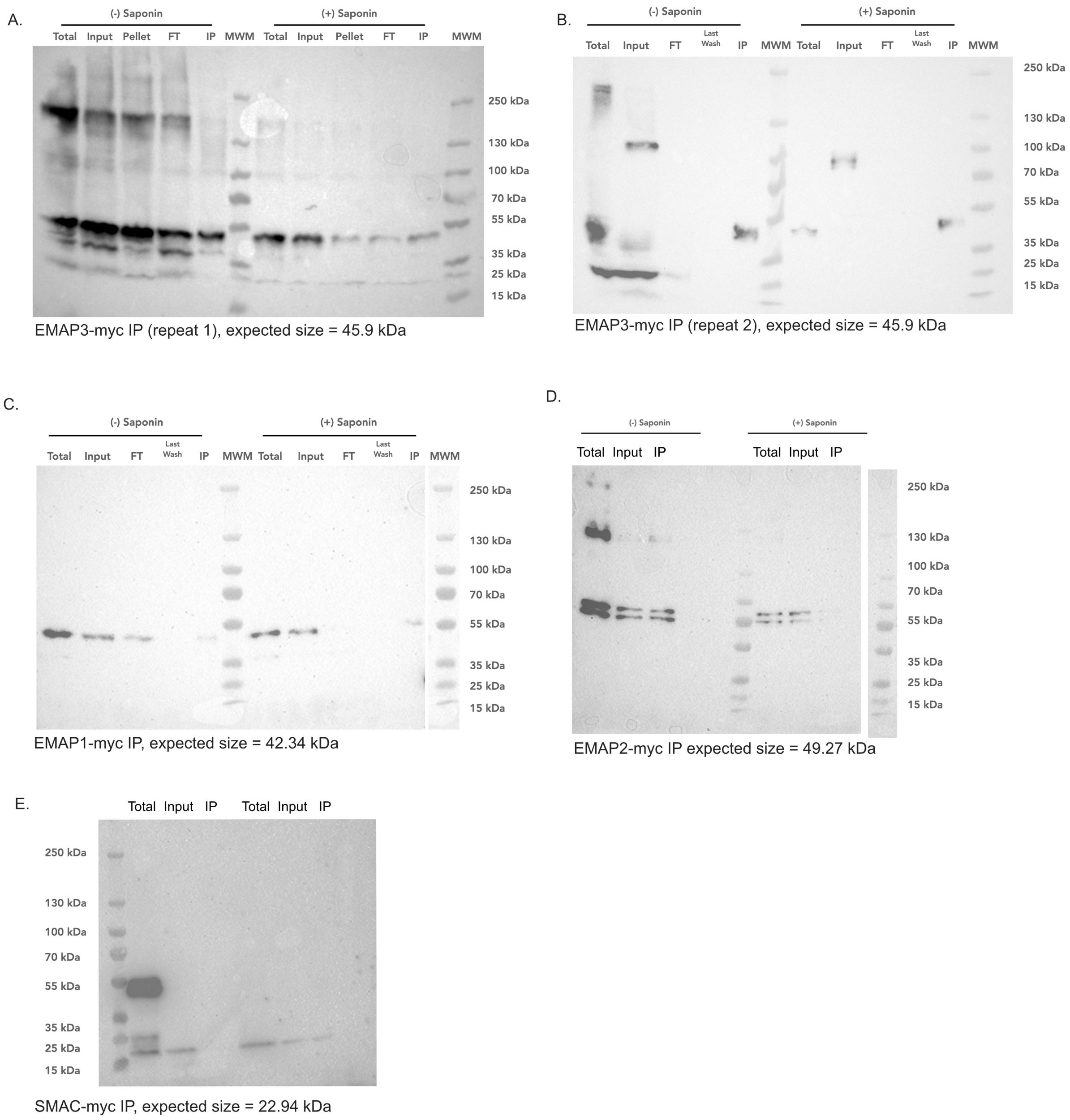


**Figure S3. Western blot analysis of co-immunoprecipitation (Co-IP) samples with 3x-myc tagged *P. berghei* proteins. (A-E)** Representative western blot analysis of 3x-myc tagged proteins in Co-IP samples with (+) and without (-) saponin. (A-B) EMAP3-myc (repeat 1 and 2), 45.9 kDa. (C) EMAP1-myc, 42.34 kDa. (D) EMAP2-myc, 49.27 kDa. (E) SMAC-myc, 22.94 kDa. The expected molecular weights indicated are including the 3x-myc tag. Immunoprecipitated sample (IP), flow-through (FT) and molecular weight marker (MWM).

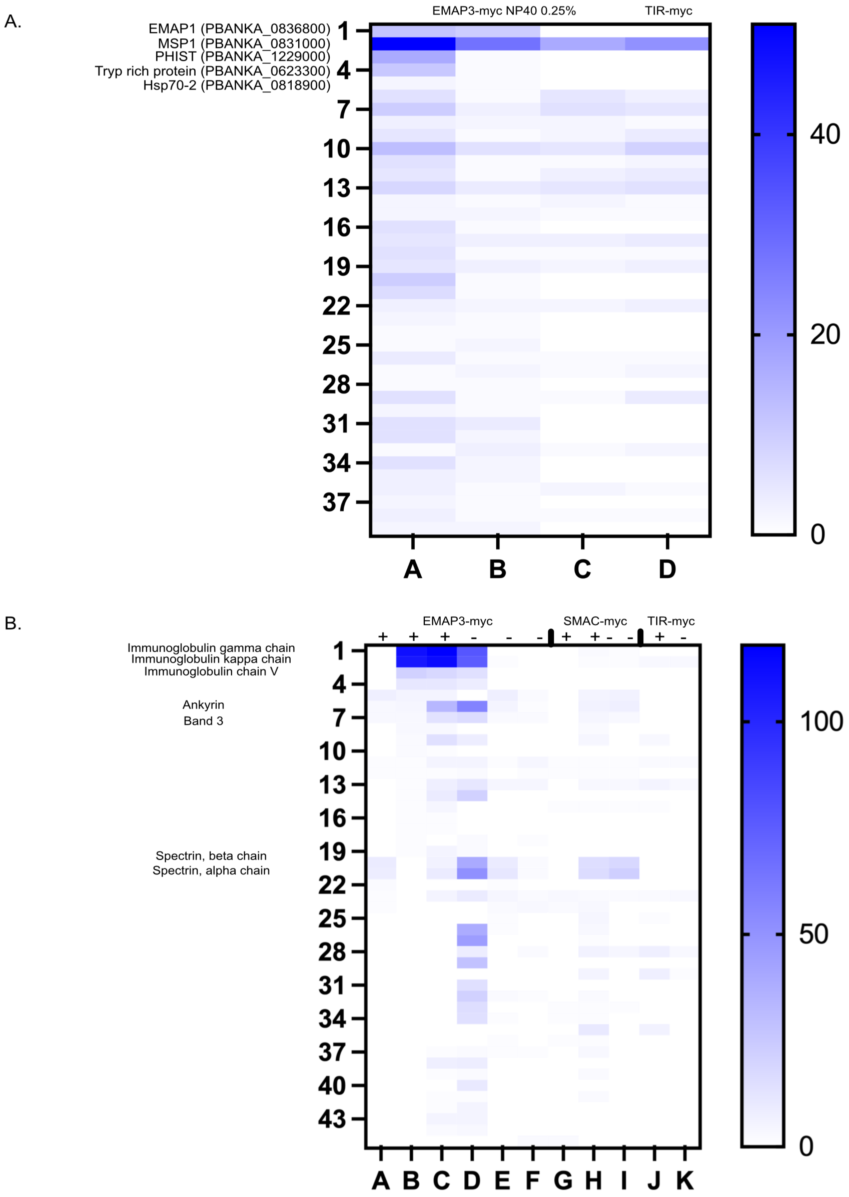


**Figure S4. Extended analysis of EMAP3-myc and SMAC-myc co-immunoprecipitation (Co-IP).** Heat maps showing: **(A)** Proteins co-precipitated in EMAP3-myc Co-IP using 0.25% NP40 detergent. **(B)** Host (mouse) proteins co-precipitated in EMAP3-myc and SMAC-myc cross linking Co-IPs with (+) and (-) without saponin. TIR1-myc was used as a non-exported control.

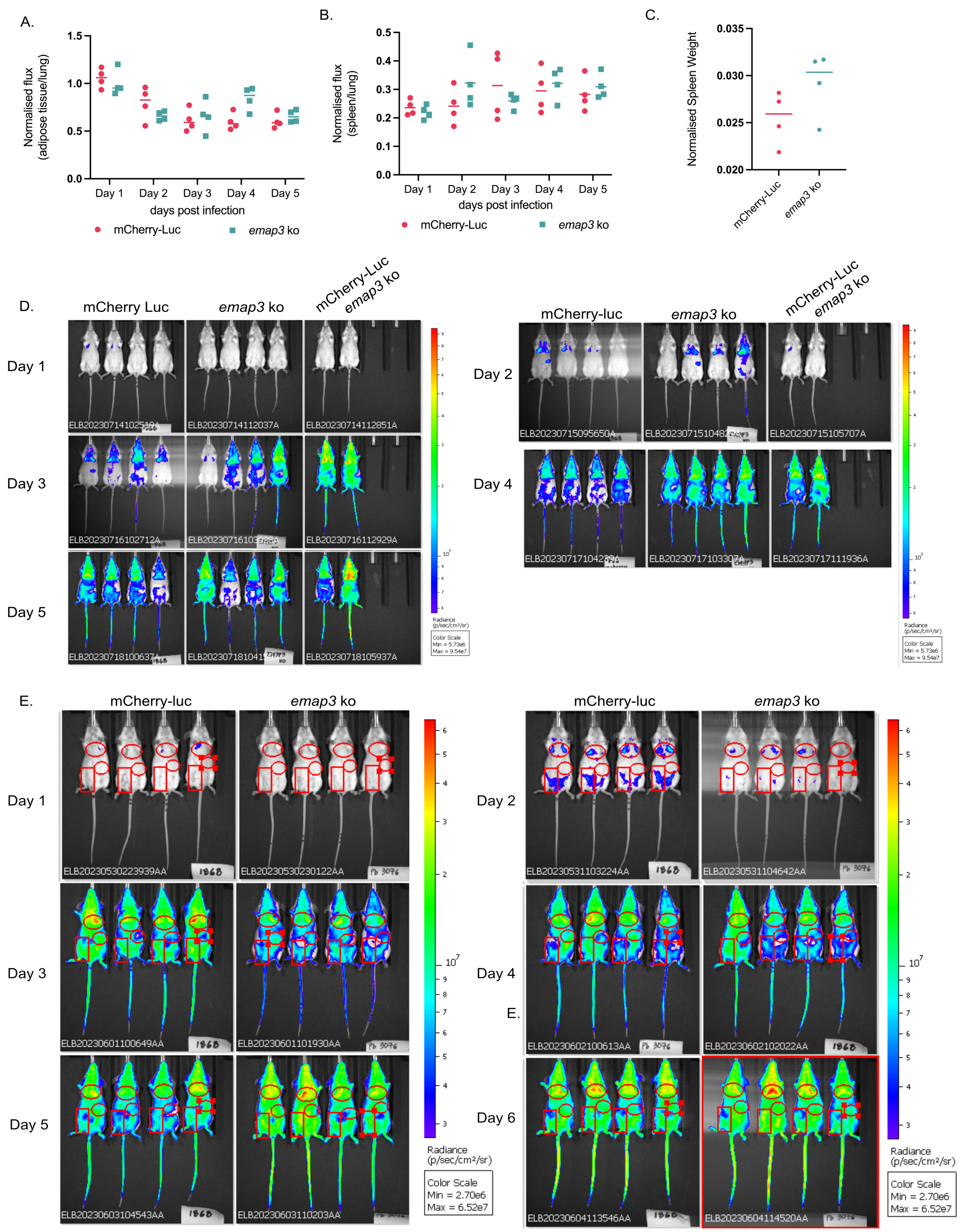


**Figure S5. Evaluation of *emap3* ko sequestration. (A-B)** Whole body imaging of *emap3 knockout (*ko) and mCherry-Luc parasites in Balb/c mice using the Spectrum *in vivo* imaging system (IVIS) to visualise luciferase expressing parasites in (A) adipose tissue and (B) spleen after injecting 10-30 x $10^6$ purified schizonts intravenously (IV). Bioluminescence signal in each organ is normalised to the signal in lungs and expressed as normalised flux. Data shown here is from a second independent biological repeat. **(C)** Spleen weights of BALB/c mice from second biological repeat of IVIS experiment (obtained day 5 post infection). Spleen weight is normalised to whole mouse body weight. **(D)** IVIS images for *emap3* ko and mCherry-Luc used to measure adipose tissue and spleen bioluminescence signals in main **Fig. 4B-C**, corresponding to the first biological repeat. **(E)** IVIS images for *emap3* ko and mCherry used to measure adipose tissue and spleen bioluminescence signals in supplemental **Fig. S5A-B**, corresponding to the second biological repeat. For the second biological repeat four mice were assayed for both parasite lines.

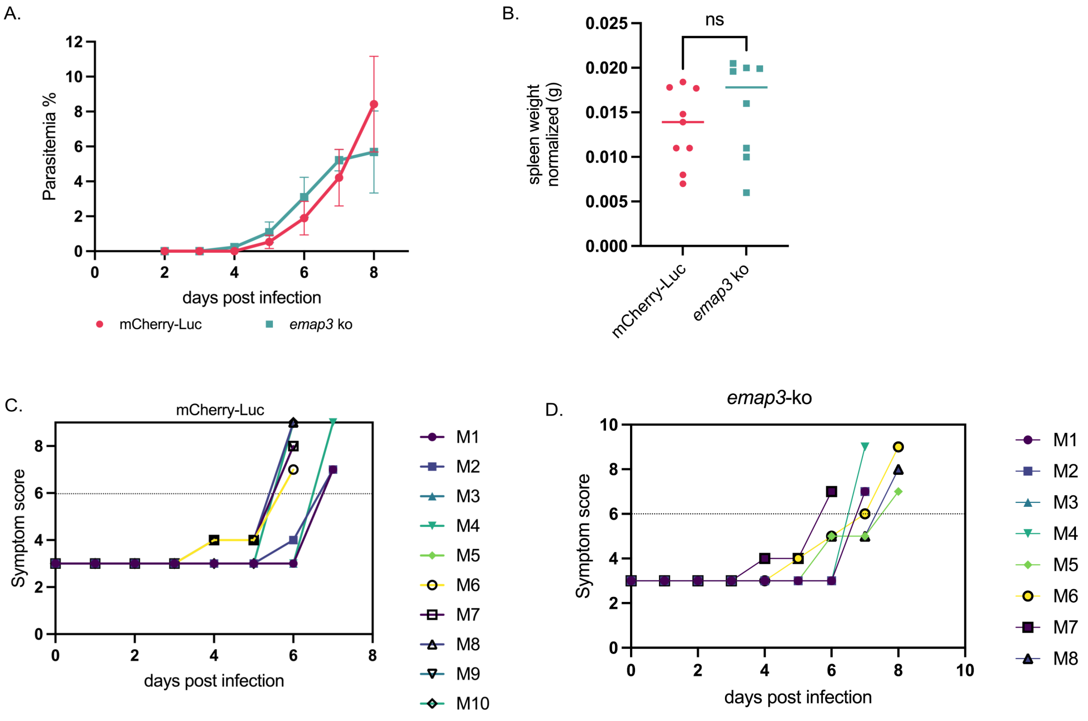


**Figure S6. Assessment of *emap3* ko virulence characteristics. (A)** *In vivo* blood stage growth assays of *emap3* ko compared to the mCherry-Luc *P. berghei* background line. C57BL/6 mice were injected with 1 x $10^6$ parasites intravenously (IV) on day 0. Parasitaemia was determined by counting parasites on Giemsa-stained thin blood smears. At each day the parasitaemia is given as an average from eight C57BL/6 mice. **(B)** Spleen weights of C57BL/6 mice infected with mCherry-Luc or *emap3* ko parasites (obtained day 7 post infection). Spleen weights were normalised to whole mouse body weights and calculated from eight (*emap3* ko) or nine (mCherry-Luc) C57BL/6 mice. **(C-D)** Symptom scoring forming basis of termination of C57BL/6 mice on the day the mice are reaching moderate symptoms (C) mCherry-Luc (D) *emap3* ko. Eight (emap3 ko) or ten (mCherry-Luc) C57BL/6 mice across two independent biological repeats were used to score symptoms of mice infected with *emap3* ko and mCherry-Luc lines. The data in panel A-D was obtained from the same C57BL/6 mice used to evaluate virulence (**main Fig. 4F**).
