## Supplemental Table 1 for "Erythrocyte membrane protein 3 (EMAP3) is exposed on the surface of the *Plasmodium berghei* infected red blood cell"

**Table S1. Supplementary tables.** Tables containing information on parasite lines, genotyping and cloning primers, antibodies for IFA, UEx-M and Western blotting, reagents used for experiments and raw data for Co-IP experiments

| Parasite line | Leiden LMRG reference | Description | Comment |
|---|---|---|---|
| mCherry.Luc | 1868Cl1 | Background line with hsp70p:mCherry eef1ap:Luciferase | |
| PbEMAP3 KO | 3076cl1 | *emap3* knockout in hsp70p:mCherry eef1ap:Luciferase background line | Generated in background line 1868Cl1 |
| PbCas9-FLAG_Tir1 -myc | N/A | Background line with hsp70p:Cas9-Flag and eef1ap:TIR-myc | Generated in background line PbCas9 |
| PbSMAC-myc | Line 1413 | PbSMAC::3xcMYC in eef1ap:GFP-Luciferase background line | Generated in background line eef1ap:GFP-Luciferase |
| PbEMAP1-myc | Line1583 | PbEMAP1::3xcMYC | |
| PbEMAP3-myc | Line 3084 | PbEMAP3::3xcMYC | |
| PbEMAP2-myc | Line 3296 | PbEMAP2::3xcMYC in hsp70p:mCherry eef1ap:Luciferase background line | Generated in background line 1868Cl1 |

| **Genotyping Primers** | | | | |
|---|---|---|---|---|
| **Line** | **Forward Primer (5' - 3')** | **Reverse Primer** | **Expected size** | **Description** |
| PbCas9-Flag Tir1-myc | GAGGCCATAGAAAATGATG TAGTTGG | CCTCATCATCTTCCTCATCATCT TCATC | 0.9 kb (WT), 3.7 kb (Tir1) | outer Fw x outer Rv |
| | GAGGCCATAGAAAATGATG TAGTTGG | CGTTGTTTGAGTCATGTGTCTT ATCTG | 0.5 kb (WT), n.d. (Tir1) | outer Fw x WT inner Rv |
| | GAGGCCATAGAAAATGATG TAGTTGG | CAGATCTTCTTCTGAAATCAAC | n.d. (WT), 2.7 kb (Tir1) | outer Fw x inner Rv |
| | GACAATGAGGAGCCTGTG GA | CCTCATCATCTTCCTCATCATCT TCATC | n.d. (WT), 1.3 kb (Tir1) | inner Fw x outer Rv |
| EMAP3-myc | CCTAGTTTAGTTTCTCTATTT | GTTACACGTATATTACGCATACA | 1,5 kb | 5' integration |

| | | | | |
|---|---|---|---|---|
| | TCTGG | ACG | | |
| | GCGAAAAACCGTCTATCAGG | CCTAGTTTAGTTTCTCTATTTTCTGG | 1,7 kb | 3' integration |
| | AATGAAGCGACGTATCGACC | GATCACATTCTTCAGCTGGTC | 0,7 kb | SM (Tgdhfr/ts) |
| | ccggatccGAACAAAAACTCATCTCAGAAG | GGGGCGGCCGCATGGAAGACGCCAAAAACATAAAG | 0,5 kb | 3xcmyc |
| | CGCAACTGAGCTATATGATGC | GGTGCACACTACTCATCTC | 0,7 kb | *emap3 orf* |
| EMAP2-myc | TATGGGATTTTCACATATTTACAAC | TTACATGTCACATGAATTATCCACCGG | 1,1 kb | 5' integration |
| | GCGAAAAACCGTCTATCAGG | gggggtaccCCACTGTTTCTTCGTAGACACC | 2,5 kb | 3' integration |
| | ccaagcttGAACAAAAACTCATCTCAGAAG | CGCTCCGCGGCTTTTTTCGTATTTTTCATTAACTTCC | 0,5 kb | 3xcmyc |
| | GATCACATTCTTCAGCTGGTC | ctcgagaagagaaggaagac | 0,6 kb | SM (Tgdhfr/ts) |
| EMAP1-myc | GCGATTAAGTTGGGTAACGC | GAACATTATGTGGGGTATACG | 1,4 kb | 3' integration |
| | CGGGATCCATGCATAAACCGGTGTGTC | CGGGATCCAAGCTTCTGTATTTCCGC | 1,9 kb | SM (Tgdhfr/ts) |
| SMAC-myc | GCCGGATCCCATTTATACATTCTAGCTTTTCAGAG | GTTGGATCCTTATATGCGTCTATTTATGTAGG | 1,8 kb | 5' integration |
| | TTTCCCAGTCACGACGTTG | ATGGTACCGGGTTGTATTATTAATATTTTATGGG | 1,4 kb | 3' integration |
| | CGGGATCCATGCATAAACCGGTGTGTC | CGGGATCCAAGCTTCTGTATTTCCGC | 1,9 kb | SM (Tgdhfr/ts) |
| EMAP3-KO | GAATTGTTCATATATTCGTAGAC | GATGTGTTATGTGATTAATTC | 0,6 kb | 5' integration |
| | CCAATACATATCTTGCATTCGTG | GTCTCTTCAATGATTCATAAATAGTTGG | 0,9 kb | 3' integration |
| | GGACAGATTGAACATCGTCG | GATCACATTCTTCAGCTGGTC | 1,0 kb | SM (Tgdhfr/ts) |
| | CGCAACTGAGCTATATGATGC | GGTGCACACTACTCATCTC | 0,7 kb | *emap3 orf* |

| **Cloning Primers** | | | | |
|---|---|---|---|---|
| **Line** | **Forward Primer (5' - 3')** | **Reverse Primer** | **Expected size** | **Description** |
| EMAP3-myc | ggACTAGTGGTATCTCAAATCGATTTACGAGC | ggGGATCCATTTTTGTATACGTCTTTACGAC | 0,7 kb | pL2302 - HR for PBANKA_0825900::3xcmyc (EMAP3) |
| *emap3*-ko | ggGGTACCAAATAAATAAATAACCATGAAAAAATC | ggAAGCTTTTTATTACAATGATATCGAC | 0,5 kb | pL2296 5' HR for PBANKA_0825900 KO (EMAP3) |
| | ggGAATTCTTTAATAAGAATGAATTTCCAG | gggGCGGCCGCTGGGTAAAATTTATTAACATATC | 0,5 kb | pL2296 3' HR for PBANKA_0825900 KO (EMAP3) |
| PbCas9-Flag Tir1-myc | ATGGCAGATCTCAATTGGATGACTCGAGGAATTCCTTAAGTTTAC | AGTATGTCATAGATCCCCCTATGTTTTATAAAATTTTTATTTATTTATAAG | | 5' ef-1α cloning primers |
| | AGGGGGATCTATGACATACTTTCCTGAAGAGGTCGTCG | TACCCCGCGGTTACAGATCTTCTTCTGAAATCAACTTTTGTTCCAGATCTTCTTCAGAGATGAGTTTCTGCTCACCACTAGCAGCAGAACCGGAGTCGACCAGAATCTTCACAAAGTTGGGAGCATCATC | | *osTir1 cloning primers* |
| | AGATCTGTAACCGCGGGGTACCCCATTAAATTTATTTAATAATAG | CGATCGCGTGGCCGGCCGATGCTCGGTACCTCTAGATTTCTCTGCGG | | *3' hrp2 cloning primers* |
| | ACTGGTTTAAAAAAGAAGGCGACTCGAGGAATTCCTTAAGTTTAC | ATGCAGAATCTTCTTGAGGCGCTCGGTACCTCTAGATTTCTCTGCGG | | osTir1 cloning into pYCs_PbU6_hDHFR/yFCU_NS_p230p plasmid |
| | CTATGACCATGATTACGCCACTGTTTTGAATATGTACAAATAGAAGGAACAG | TCTTCTTGAGGCGGCCGCCTTCTTTTTTAAACCAGTTATTTAAATC | | p230 5' HR |
| | AGGCGGCCGCCTCAAGAAGATTCTGCATCTACCG | ACCATGGCATGTTGGTACCACAATGTGTTCTAAATTTAGTAATGATATATTTTTTTC | | p230 3' HR |
| | TATTGAATCTTAACCTGTTTAATTG | AAACCAATTAAACAGGTTAAGATTC | | gRNA1 |
| | TATTGATAATATATATGCATACCAG | AAACCTGGTATGCATATATATTATC | | *gRNA2* |

**IFA antibodies**

| *Primary antibodies* | Working dilution | Species | Supplier |
|---|---|---|---|
| 3x myc (7D10) | 1:200 | Rabbit | Cell Signaling Technologies |
| BiP | 1:500 | Mouse | |
| EXP1 | 1:500 | Mouse | |

| *Secondary antibodies* | Working dilution | Supplier |
|---|---|---|
| TER-119 | 1:500 | Invitrogen |
| α-rabbit Alexa Fluor 405 | 1:500 | Invitrogen |
| α-mouse Alexa Fluor 594 | 1:500 | Invitrogen |
| α-rabbit Alexa Fluor 647 | 1:500 | Invitrogen |

| *Stains* | Working dilution | | Supplier |
|---|---|---|---|
| Atto 488 NHS-ester | 10 ug/mL | | Merck |
| BODIPY trc | 6 uM | | Thermofisher |

**Western blot antibodies**

| *Primary antibodies* | Working dilution | Species | Supplier |
|---|---|---|---|
| 3x myc (7D10) | 1:1,000 | rabbit | Cell Signaling Technology |
| FLAG tag | 1:1,000 | rabbit | Cell Signaling Technology |

| *Secondary antibodies* | Working dilution | Supplier |
|---|---|---|
| rabbit-HRP | 1:10,000 | Cell Signaling Technology |

**Chemicals**

| Name | Stock concentration | Supplier |
|---|---|---|
| Pyrmethamine | | Sigma |
| Giemsa | | Sigma |
| d-luciferin sodium salt | 80 mg/mL | Synchem OHG; cat. no. BC218 |
| Paraformaldehyde | | Pierce, Invitrogen |

| | | |
|---|---|---|
| Poly-d-lysine | 0.1 mg/mL | A3890401, GIBCO |
| Mounting media with DAPI | | VECTASHIELD |
| BSA | | Sigma |
| Protein G Dynabeads | | Invitrogen |
| Percoll | | Cytiva |
| Formaldehyde | 36.5-38% | F8775, Sigma |
| Acrylamide | 40% | A4058, Sigma |
| N,N0 -methylenbisacrylamide | 2% | M1533, Sigma |
| Sodium Acrylate | | R624, AK Scientific |
| Ammonium persulfate | | 17874, ThermoFisher |
| Tetramethylethylenediamine | | 17919, ThermoFisher |
| Brefeldin A | 5 mg/mL | 20350-15-6, Sigma |
| Histodenz | 27.6% w/v | Sigma |

**Crosslinked IP (without and with saponin) for EMAP1, EMAP2, EMAP3 and SMAC**

| Accession Number | EMAP1-myc + saponin N2 | EMAP1 +saponin | EMAP1-myc - saponin N2 | EMAP1 - saponin | EMAP2 +saponin | EMAP2-myc + saponin N2 | EMAP2 - saponin | EMAP2-myc - saponin N2 | EMAP3-myc + saponin N2 | EMAP3-myc + saponin | EMAP3-myc + saponin CST ab | EMAP3-myc - saponin | EMAP3-myc - saponin CST ab | EMAP3-myc - saponin N2 | SMAC-myc + saponin N2 | SMAC-myc + saponin N1 | SMAC-myc - saponin N2 | SMAC-myc - saponin N1 | TIRI-myc - saponin CST ab | TIRI-myc + saponin CST ab |
|---|---|---|---|---|---|---|---|---|---|---|---|---|---|---|---|---|---|---|---|---|
| PBANKA_0836800.1-p1 | 6 | 2 | 11 | 2 | 0 | 2 | 0 | 1 | 6 | 11 | 5 | 23 | 16 | 9 | 0 | 0 | 0 | 0 | 0 | 0 |
| PBANKA_1229000.1-p1 | 4 | 0 | 7 | 0 | 0 | 17 | 0 | 9 | 5 | 0 | 0 | 0 | 2 | 7 | 5 | 0 | 7 | 3 | 0 | 0 |
| PBANKA_0623300.1-p1 | 2 | 0 | 4 | 0 | 0 | 11 | 0 | 10 | 5 | 0 | 0 | 1 | 1 | 6 | 4 | 0 | 5 | 2 | 0 | 0 |
| PBANKA_0818900.1-p1 | 2 | 2 | 4 | 2 | 0 | 2 | 0 | 1 | 5 | 6 | 2 | 21 | 15 | 4 | 0 | 0 | 1 | 1 | 0 | 1 |
| PBANKA_0715500.1-p1 | 1 | 0 | 1 | 0 | 0 | 1 | 0 | 1 | 1 | 0 | 0 | 0 | 0 | 2 | 1 | 0 | 1 | 0 | 0 | 0 |
| PBANKA_0405300.1-p1 | 1 | 0 | 1 | 0 | 0 | 1 | 0 | 2 | 0 | 0 | 0 | 0 | 0 | 2 | 1 | 0 | 1 | 0 | 0 | 0 |
| PBANKA_0906500.1-p1 | 1 | 0 | 0 | 0 | 0 | 1 | 1 | 2 | 0 | 0 | 0 | 0 | 0 | 1 | 1 | 0 | 1 | 0 | 0 | 0 |
| PBANKA_0717800.1-p1 | 1 | 0 | 0 | 0 | 0 | 1 | 0 | 1 | 1 | 0 | 0 | 0 | 0 | 3 | 0 | 0 | 1 | 0 | 0 | 0 |
| PBANKA_0831000.1-p1 | 0 | 1 | 0 | 1 | 1 | 0 | 1 | 0 | 0 | 0 | 0 | 2 | 0 | 0 | 0 | 0 | 0 | 0 | 0 | 0 |
| PBANKA_0830200.1-p1 | 0 | 0 | 0 | 0 | 0 | 0 | 0 | 0 | 0 | 0 | 0 | 3 | 3 | 0 | 0 | 0 | 0 | 0 | 0 | 0 |
| PBANKA_0941900.1-p1 | 0 | 8 | 0 | 0 | 6 | 0 | 0 | 2 | 6 | 0 | 0 | 0 | 5 | 8 | 0 | 1 | 2 | 2 | 0 | 0 |
| PBANKA_1400600.1-p1 | 0 | 0 | 0 | 0 | 0 | 0 | 0 | 0 | 0 | 0 | 0 | 2 | 1 | 0 | 0 | 0 | 0 | 0 | 0 | 0 |
| PBANKA_1326400.1-p1 | 0 | 0 | 0 | 0 | 0 | 4 | 0 | 2 | 0 | 0 | 2 | 1 | 2 | 0 | 1 | 0 | 0 | 1 | 0 | 0 |
| PBANKA_1231000.1-p1 (+1) | 0 | 1 | 2 | 1 | 0 | 3 | 1 | 3 | 1 | 0 | 0 | 0 | 0 | 2 | 1 | 0 | 2 | 1 | 0 | 0 |
| PBANKA_0416000.1-p1 | 0 | 0 | 0 | 0 | 0 | 0 | 0 | 0 | 0 | 0 | 0 | 3 | 2 | 0 | 0 | 0 | 0 | 0 | 0 | 0 |
| PBANKA_1459300.1-p1 | 0 | 0 | 0 | 0 | 0 | 1 | 0 | 1 | 1 | 0 | 1 | 0 | 1 | 2 | 1 | 0 | 2 | 1 | 0 | 0 |
| PBANKA_1145800.1-p1 | 0 | 0 | 2 | 0 | 0 | 2 | 0 | 2 | 0 | 0 | 0 | 0 | 0 | 0 | 2 | 0 | 2 | 0 | 0 | 0 |
| PBANKA_1200600.1-p1 | 0 | 0 | 1 | 1 | 0 | 1 | 0 | 0 | 0 | 1 | 0 | 1 | 4 | 0 | 0 | 0 | 1 | 0 | 0 | 0 |
| PBANKA_1407600.1-p1 | 0 | 0 | 1 | 0 | 0 | 2 | 0 | 3 | 1 | 0 | 0 | 0 | 0 | 3 | 0 | 0 | 2 | 0 | 0 | 0 |
| PBANKA_1000600.1-p1 | 0 | 0 | 0 | 0 | 0 | 1 | 0 | 1 | 0 | 0 | 0 | 1 | 2 | 0 | 0 | 0 | 0 | 0 | 0 | 0 |
| PBANKA_1237800.1-p1 | 0 | 0 | 0 | 0 | 0 | 4 | 0 | 1 | 0 | 0 | 0 | 0 | 0 | 0 | 1 | 0 | 1 | 0 | 0 | 0 |
| PBANKA_1437300.1-p1 | 0 | 0 | 0 | 0 | 0 | 0 | 0 | 0 | 0 | 2 | 1 | 11 | 4 | 0 | 0 | 0 | 0 | 0 | 0 | 0 |
| PBANKA_1223800.1-p1 | 0 | 0 | 0 | 0 | 0 | 1 | 0 | 0 | 4 | 0 | 0 | 0 | 0 | 4 | 0 | 0 | 0 | 0 | 0 | 0 |
| PBANKA_0523700.1-p1 | 0 | 0 | 0 | 0 | 0 | 1 | 0 | 0 | 1 | 0 | 0 | 0 | 0 | 2 | 0 | 0 | 0 | 0 | 0 | 0 |
| PBANKA_0100600.1-p1 | 0 | 0 | 0 | 0 | 0 | 0 | 0 | 0 | 0 | 0 | 0 | 0 | 0 | 0 | 5 | 0 | 4 | 1 | 0 | 0 |
| PBANKA_0316800.1-p1 | 0 | 0 | 0 | 0 | 0 | 3 | 0 | 2 | 0 | 0 | 0 | 1 | 1 | 0 | 0 | 0 | 0 | 0 | 0 | 0 |
| PBANKA_0836200.1-p1 | 0 | 0 | 0 | 0 | 0 | 2 | 0 | 2 | 0 | 0 | 0 | 0 | 0 | 0 | 1 | 0 | 2 | 0 | 0 | 0 |
| PBANKA_0517000.1-p1 | 0 | 0 | 0 | 0 | 0 | 0 | 0 | 0 | 0 | 0 | 0 | 3 | 2 | 0 | 0 | 0 | 0 | 0 | 0 | 0 |
| PBANKA_0914300.1-p1 | 0 | 0 | 0 | 0 | 0 | 1 | 0 | 0 | 0 | 1 | 0 | 2 | 0 | 0 | 0 | 0 | 0 | 0 | 0 | 0 |
| PBANKA_0702800.1-p1 | 0 | 0 | 0 | 0 | 0 | 0 | 0 | 0 | 0 | 1 | 0 | 7 | 3 | 0 | 0 | 0 | 0 | 0 | 0 | 0 |
| PBANKA_1423300.1-p1 | 0 | 0 | 0 | 0 | 0 | 0 | 0 | 0 | 0 | 0 | 0 | 3 | 1 | 1 | 0 | 0 | 0 | 0 | 0 | 0 |
| PBANKA_1357200.1-p1 | 0 | 0 | 0 | 0 | 0 | 0 | 0 | 0 | 0 | 0 | 0 | 2 | 1 | 0 | 0 | 0 | 0 | 0 | 0 | 0 |

**Crosslinked IP mouse protein hits for EMAP3 and SMAC**

| **Identified proteins** | **EMAP3-myc + saponin N2** | **EMAP3-myc + saponin** | **EMAP3-myc - saponin** | **EMAP3-myc - saponin CST ab** | **EMAP3-myc - saponin N2** | **SMAC-myc - saponin N1** | **SMAC-myc + saponin N1** | **SMAC-myc - saponin N2** | **SMAC-myc + saponin N2** | **TIRI-myc - saponin CST ab** | **TIRI-myc + saponin CST ab** |
|---|---|---|---|---|---|---|---|---|---|---|---|
| Ig gamma-1 chain C region secreted form OS=Mus musculus OX=10090 GN=Ighg1 PE=1 SV=1 | 0 | 111 | 118 | 78 | 0 | 0 | 0 | 2 | 1 | 0 | 0 |
| Immunoglobulin kappa constant OS=Mus musculus OX=10090 GN=Igkc PE=1 SV=2 | 0 | 107 | 113 | 75 | 1 | 0 | 0 | 1 | 1 | 3 | 3 |
| Ig kappa chain V-III region PC 2880/PC 1229 OS=Mus musculus OX=10090 PE=1 SV=1 | 0 | 21 | 18 | 14 | 0 | 0 | 0 | 0 | 0 | 0 | 0 |
| Immunoglobulin heavy variable 5-4 (Fragment) OS=Mus musculus OX=10090 GN=Ighv5-4 PE=4 SV=1 | 0 | 11 | 11 | 8 | 0 | 0 | 0 | 0 | 0 | 0 | 0 |
| Progranulin OS=Mus musculus OX=10090 GN=Grn PE=1 SV=2 | 7 | 5 | 5 | 0 | 7 | 2 | 0 | 5 | 6 | 0 | 0 |
| Ankyrin-1 OS=Mus musculus OX=10090 GN=Ank1 PE=1 SV=2 | 3 | 4 | 33 | 57 | 5 | 1 | 0 | 6 | 7 | 0 | 0 |
| Band 3 anion transport protein OS=Mus musculus OX=10090 GN=Slc4a1 PE=1 SV=1 | 3 | 3 | 13 | 16 | 2 | 2 | 0 | 4 | 3 | 0 | 0 |
| Ig gamma-2B chain C region OS=Mus musculus OX=10090 GN=Igh-3 PE=1 SV=3 | 0 | 3 | 3 | 0 | 0 | 0 | 0 | 2 | 0 | 0 | 0 |
| Glyceraldehyde-3-phosphate dehydrogenase OS=Mus musculus OX=10090 GN=Gapdh PE=1 SV=2 | 0 | 2 | 14 | 8 | 0 | 0 | 0 | 4 | 0 | 3 | 0 |
| Antileukoproteinase OS=Mus musculus OX=10090 GN=Slpi PE=1 SV=1 | 0 | 2 | 1 | 0 | 0 | 0 | 0 | 0 | 0 | 0 | 0 |
| Hemoglobin subunit alpha OS=Mus musculus OX=10090 GN=Hba PE=1 SV=2 | 1 | 1 | 5 | 5 | 1 | 4 | 1 | 1 | 1 | 2 | 2 |
| Albumin OS=Mus musculus OX=10090 GN=Alb PE=1 SV=3 | 1 | 1 | 1 | 2 | 0 | 1 | 1 | 1 | 1 | 0 | 0 |
| Hemoglobin subunit beta-1 OS=Mus musculus OX=10090 GN=Hbb-b1 PE=1 SV=2 | 0 | 1 | 7 | 11 | 4 | 4 | 0 | 3 | 3 | 5 | 3 |
| Protein 4.2 OS=Mus musculus OX=10090 GN=Epb42 PE=1 SV=3 | 0 | 1 | 9 | 21 | 0 | 0 | 0 | 0 | 0 | 0 | 0 |
| Heat shock 70 kDa protein 1-like OS=Mus musculus OX=10090 GN=Hspa1l PE=1 SV=4 | 0 | 1 | 4 | 0 | 0 | 0 | 1 | 1 | 1 | 1 | 0 |
| Histone H3.3C OS=Mus musculus OX=10090 GN=H3-5 PE=3 SV=3 | 0 | 1 | 1 | 0 | 0 | 0 | 0 | 0 | 0 | 0 | 0 |
| Complement C4-B OS=Mus musculus OX=10090 GN=C4b PE=1 SV=3 | 0 | 1 | 1 | 0 | 0 | 0 | 0 | 0 | 0 | 0 | 0 |
| Keratin, type II cuticular Hb2 OS=Mus musculus OX=10090 GN=Krt82 PE=1 SV=2 | 0 | 1 | 0 | 2 | 0 | 1 | 0 | 0 | 0 | 0 | 0 |
| Rab GTPase-binding effector protein 2 OS=Mus musculus OX=10090 GN=Rabep2 PE=1 SV=3 | 0 | 1 | 5 | 2 | 0 | 0 | 0 | 0 | 0 | 0 | 0 |
| Spectrin beta chain, erythrocytic OS=Mus musculus OX=10090 GN=Sptb PE=1 SV=4 | 9 | 0 | 6 | 39 | 9 | 2 | 0 | 15 | 18 | 0 | 0 |
| Spectrin alpha chain, erythrocytic 1 OS=Mus musculus OX=10090 GN=Spta1 PE=1 SV=3 | 8 | 0 | 9 | 51 | 11 | 2 | 0 | 16 | 22 | 0 | 0 |
| Immunoglobulin heavy constant mu OS=Mus musculus OX=10090 GN=Ighm PE=1 SV=2 | 2 | 0 | 0 | 0 | 1 | 0 | 0 | 0 | 0 | 0 | 0 |
| Hemoglobin subunit beta-2 OS=Mus musculus OX=10090 GN=Hbb-b2 PE=1 SV=2 | 1 | 0 | 5 | 9 | 4 | 2 | 3 | 2 | 2 | 2 | 3 |
| Inter-alpha-trypsin inhibitor heavy chain H2 OS=Mus musculus OX=10090 GN=Itih2 PE=1 SV=1 | 1 | 0 | 0 | 0 | 2 | 3 | 2 | 3 | 0 | 0 | 0 |
| Ig gamma-3 chain C region OS=Mus musculus OX=10090 PE=1 SV=2 | 0 | 0 | 0 | 0 | 1 | 0 | 0 | 4 | 0 | 1 | 0 |
| Myosin-9 OS=Mus musculus OX=10090 GN=Myh9 PE=1 SV=4 | 0 | 0 | 0 | 38 | 1 | 0 | 0 | 3 | 0 | 0 | 0 |
| Talin-1 OS=Mus musculus OX=10090 GN=Tln1 PE=1 SV=2 | 0 | 0 | 0 | 45 | 0 | 0 | 0 | 1 | 0 | 0 | 0 |
| Keratin, type II cytoskeletal 1b OS=Mus musculus OX=10090 GN=Krt77 PE=1 SV=1 | 0 | 0 | 0 | 8 | 0 | 2 | 0 | 6 | 4 | 7 | 3 |
| Filamin-A OS=Mus musculus OX=10090 GN=Flna PE=1 SV=5 | 0 | 0 | 0 | 28 | 0 | 0 | 0 | 0 | 0 | 0 | 0 |
| Desmoplakin OS=Mus musculus OX=10090 GN=Dsp PE=1 SV=1 | 0 | 0 | 0 | 0 | 0 | 0 | 0 | 5 | 0 | 7 | 1 |
| Fibrinogen gamma chain OS=Mus musculus OX=10090 GN=Fgg PE=1 SV=1 | 0 | 0 | 0 | 14 | 0 | 0 | 0 | 0 | 0 | 0 | 0 |
| Thrombospondin-1 OS=Mus musculus OX=10090 GN=Thbs1 PE=1 SV=1 | 0 | 0 | 0 | 21 | 2 | 1 | 0 | 2 | 0 | 0 | 0 |
| Fibrinogen beta chain OS=Mus musculus OX=10090 GN=Fgb PE=1 SV=1 | 0 | 0 | 0 | 15 | 0 | 0 | 1 | 1 | 1 | 0 | 0 |
| Fibrinogen alpha chain OS=Mus musculus OX=10090 GN=Fga PE=1 SV=1 | 0 | 0 | 0 | 14 | 1 | 0 | 1 | 1 | 0 | 0 | 0 |
| Keratin, type I cytoskeletal 19 OS=Mus musculus OX=10090 GN=Krt19 PE=1 SV=1 | 0 | 0 | 0 | 0 | 0 | 0 | 0 | 10 | 0 | 6 | 0 |
| Pigment epithelium-derived factor OS=Mus musculus OX=10090 GN=Serpinf1 PE=1 SV=2 | 0 | 0 | 0 | 0 | 1 | 0 | 1 | 1 | 0 | 0 | 0 |
| Predicted pseudogene 8797 OS=Mus musculus OX=10090 GN=Gm8797 PE=4 SV=1 | 0 | 0 | 1 | 2 | 1 | 1 | 0 | 3 | 0 | 0 | 0 |

| | | | | | | | | | | | |
|---|---|---|---|---|---|---|---|---|---|---|---|
| Beta-globin OS=Mus musculus OX=10090 GN=Hbb-bs PE=1 SV=1 | 0 | 0 | 7 | 8 | 0 | 0 | 0 | 0 | 0 | 0 | 0 |
| Tubulin alpha-1B chain OS=Mus musculus OX=10090 GN=Tuba1b PE=1 SV=2 | 0 | 0 | 1 | 2 | 0 | 0 | 0 | 2 | 0 | 0 | 0 |
| Vinculin OS=Mus musculus OX=10090 GN=Vcl PE=1 SV=4 | 0 | 0 | 0 | 10 | 0 | 0 | 0 | 0 | 0 | 0 | 0 |
| L-lactate dehydrogenase C chain OS=Mus musculus OX=10090 GN=Ldhc PE=1 SV=2 | 0 | 0 | 1 | 1 | 0 | 0 | 0 | 2 | 0 | 0 | 0 |
| Tubulin alpha-4A chain OS=Mus musculus OX=10090 GN=Tuba4a PE=1 SV=1 | 0 | 0 | 1 | 5 | 0 | 0 | 0 | 0 | 0 | 0 | 0 |
| Heat shock cognate 71 kDa protein OS=Mus musculus OX=10090 GN=Hspa8 PE=1 SV=1 | 0 | 0 | 5 | 5 | 0 | 0 | 0 | 0 | 0 | 0 | 0 |
| Stomatin OS=Mus musculus OX=10090 GN=Stom PE=1 SV=3 | 0 | 0 | 2 | 3 | 0 | 0 | 0 | 0 | 0 | 0 | 0 |
| Apolipoprotein B-100 OS=Mus musculus OX=10090 GN=Apob PE=1 SV=1 | 0 | 0 | 0 | 0 | 0 | 2 | 2 | 0 | 0 | 0 | 0 |

IP raw data (without and with saponin, NP40) for EMAP1, EMAP2, EMAP3 and SMAC

| | EMAP2-myc - saponin N2 | EMAP1-myc + saponin N2 | EMAP1-myc - saponin N2 | EMAP2-myc + saponin N2 | EMAP3-myc - saponin N2 | EMAP1 +saponin | EMAP1 - saponin | EMAP2 +saponin | EMAP2 - saponin | EMAP3 0.25% NP40 (R1) | EMAP3 0.25% NP40 (R2) | EMAP3 0.25% NP40 (R3) | EMAP3 0.25% NP40 (R4) | EMAP3-myc + saponin | EMAP3-myc + saponin CST ab | EMAP3-myc - saponin | EMAP3-myc - saponin CST ab | EMAP3-myc + saponin N2 | SMAC-myc - saponin N1 | SMAC-myc + saponin N1 | SMAC-myc - saponin N2 | SMAC-myc + saponin N2 | TIRI-myc 0.25% NP40 (R1) | TIRI-myc 0.25% NP40 (R2) | TIRI-myc - saponin CST ab | TIRI-myc + saponin CST ab |
|---|---|---|---|---|---|---|---|---|---|---|---|---|---|---|---|---|---|---|---|---|---|---|---|---|---|---|
| Immunoglobulin G-binding protein G OS=Streptococcus sp. group G OX=1320 GN=spg PE=1 SV=1 | 187 | 165 | 193 | 178 | 191 | 127 | 132 | 119 | 86 | 243 | 168 | 339 | 305 | 83 | 104 | 87 | 119 | 191 | 181 | 125 | 191 | 180 | 354 | 407 | 557 | 580 |
| Trypsin OS=Sus scrofa PE=1 SV=1 | 46 | 38 | 64 | 47 | 71 | 29 | 24 | 30 | 15 | 235 | 274 | 157 | 152 | 19 | 19 | 18 | 20 | 64 | 70 | 49 | 51 | 45 | 173 | 119 | 270 | 296 |
| Keratin, type I cytoskeletal 10 OS=Homo sapiens GN=KRT10 PE=1 SV=6 | 39 | 27 | 58 | 36 | 22 | 36 | 33 | 23 | 18 | 31 | 52 | 48 | 24 | 41 | 24 | 37 | 53 | 18 | 26 | 18 | 62 | 47 | 31 | 40 | 59 | 44 |
| Keratin, type II cytoskeletal 1 OS=Homo sapiens GN=KRT1 PE=1 SV=6 | 36 | 34 | 51 | 52 | 22 | 25 | 28 | 22 | 15 | 48 | 66 | 45 | 25 | 25 | 26 | 32 | 40 | 15 | 26 | 22 | 44 | 37 | 27 | 27 | 42 | 44 |
| Keratin, type I cytoskeletal 9 OS=Homo sapiens GN=KRT9 PE=1 SV=3 | 33 | 19 | 31 | 42 | 14 | 17 | 18 | 23 | 9 | 46 | 80 | 42 | 27 | 20 | 19 | 32 | 32 | 6 | 19 | 16 | 34 | 27 | 27 | 27 | 36 | 46 |
| Keratin, type II cytoskeletal 2 epidermal OS=Homo sapiens GN=KRT2 PE=1 SV=2 | 27 | 13 | 30 | 21 | 10 | 23 | 27 | 14 | 13 | 31 | 28 | 28 | 21 | 16 | 17 | 20 | 31 | 11 | 12 | 13 | 28 | 18 | 23 | 35 | 36 | 35 |
| Spectrin alpha chain, erythrocytic 1 OS=Mus musculus OX=10090 GN=Spta1 PE=1 SV=3 | 36 | 1 | 15 | 52 | 11 | 5 | 4 | 15 | 12 | 11 | 1 | 146 | 4 | 0 | 1 | 9 | 51 | 8 | 2 | 0 | 16 | 22 | 12 | 51 | 0 | 0 |
| Ig gamma-1 chain C region secreted form OS=Mus musculus OX=10090 GN=Ighg1 PE=1 SV=1 | 0 | 1 | 2 | 1 | 0 | 0 | 1 | 0 | 1 | 2 | 2 | 21 | 5 | 111 | 57 | 118 | 78 | 0 | 0 | 0 | 2 | 1 | 8 | 18 | 0 | 0 |
| Immunoglobulin kappa constant OS=Mus musculus OX=10090 GN=Igkc PE=1 SV=2 | 1 | 0 | 1 | 2 | 1 | 0 | 0 | 0 | 0 | 9 | 8 | 44 | 6 | 107 | 50 | 113 | 75 | 0 | 0 | 0 | 1 | 1 | 7 | 38 | 3 | 3 |
| Spectrin beta chain, erythrocytic OS=Mus musculus OX=10090 GN=Sptb PE=1 SV=4 | 34 | 0 | 11 | 47 | 9 | 3 | 1 | 14 | 11 | 4 | 0 | 103 | 1 | 0 | 0 | 6 | 39 | 9 | 2 | 0 | 15 | 18 | 4 | 40 | 0 | 0 |
| Serum albumin OS=Bos taurus GN=ALB PE=1 SV=4 | 28 | 16 | 14 | 41 | 19 | 12 | 0 | 0 | 2 | 16 | 11 | 25 | 5 | 24 | 5 | 25 | 4 | 12 | 14 | 8 | 16 | 11 | 11 | 37 | 4 | 7 |
| Ankyrin-1 OS=Mus musculus OX=10090 GN=Ank1 PE=1 SV=2 | 20 | 1 | 5 | 22 | 5 | 2 | 5 | 12 | 7 | 8 | 3 | 57 | 3 | 4 | 4 | 33 | 57 | 3 | 1 | 0 | 6 | 7 | 0 | 11 | 0 | 0 |
| Keratin, type II cytoskeletal 5 OS=Mus musculus OX=10090 GN=Krt5 PE=1 SV=1 | 3 | 2 | 6 | 7 | 0 | 5 | 7 | 3 | 3 | 16 | 14 | 18 | 10 | 3 | 0 | 4 | 3 | 0 | 1 | 0 | 11 | 6 | 9 | 12 | 20 | 10 |
| Actin, cytoplasmic 1 OS=Mus musculus OX=10090 GN=Actb PE=1 SV=1 | 7 | 3 | 5 | 11 | 8 | 7 | 0 | 7 | 4 | 6 | 6 | 22 | 6 | 1 | 0 | 2 | 8 | 5 | 1 | 4 | 12 | 4 | 5 | 9 | 6 | 9 |
| Keratin, type II cytoskeletal 6A OS=Mus musculus OX=10090 GN=Krt6a PE=1 SV=3 | 0 | 0 | 17 | 7 | 3 | 0 | 8 | 0 | 0 | 16 | 12 | 12 | 8 | 5 | 2 | 5 | 6 | 0 | 0 | 0 | 22 | 7 | 5 | 10 | 18 | 9 |
| \| transcript=PBANKA_0836800.1 \| gene=PBANKA_0836800 \| organism=Plasmodium_berghei_ANKA \| gene_product=erythrocyte membrane associated protein 1 \| transcript_product=erythrocyte membrane associated protein 1 \| location=PbANKA_08_v3:1349533-1351266(-) \| protei | 1 | 6 | 11 | 2 | 9 | 2 | 2 | 0 | 0 | 11 | 9 | 12 | 10 | 11 | 5 | 23 | 16 | 6 | 0 | 0 | 0 | 0 | 0 | 2 | 0 | 0 |
| Band 3 anion transport protein OS=Mus musculus OX=10090 GN=Slc4a1 PE=1 SV=1 | 6 | 0 | 4 | 7 | 2 | 1 | 3 | 6 | 3 | 2 | 1 | 10 | 1 | 3 | 4 | 13 | 16 | 3 | 2 | 0 | 4 | 3 | 1 | 1 | 0 | 0 |
| Ig gamma-3 chain C region OS=Mus musculus OX=10090 PE=1 SV=2 | 2 | 1 | 6 | 3 | 1 | 0 | 0 | 0 | 0 | 11 | 6 | 38 | 12 | 0 | 0 | 0 | 0 | 0 | 0 | 0 | 4 | 0 | 10 | 17 | 1 | 0 |
| Keratin, type I cytoskeletal 42 OS=Mus musculus OX=10090 GN=Krt42 PE=1 SV=1 | 7 | 0 | 12 | 7 | 0 | 0 | 0 | 0 | 0 | 9 | 10 | 10 | 6 | 5 | 0 | 7 | 10 | 0 | 0 | 0 | 15 | 5 | 0 | 6 | 11 | 7 |
| Hemoglobin subunit beta-1 OS=Mus musculus OX=10090 GN=Hbb-b1 PE=1 SV=2 | 8 | 2 | 2 | 7 | 4 | 2 | 1 | 1 | 1 | 4 | 4 | 19 | 5 | 1 | 0 | 7 | 11 | 0 | 4 | 0 | 3 | 3 | 13 | 14 | 5 | 3 |
| Myosin-9 OS=Mus musculus OX=10090 GN=Myh9 PE=1 SV=4 | 0 | 0 | 0 | 1 | 1 | 10 | 0 | 8 | 0 | 0 | 0 | 62 | 0 | 0 | 0 | 0 | 38 | 0 | 0 | 0 | 3 | 0 | 0 | 4 | 0 | 0 |
| \| transcript=PBANKA_0831000.1 \| gene=PBANKA_0831000 \| organism=Plasmodium_berghei_ANKA \| gene_product=merozoite surface protein 1 \| transcript_product=merozoite surface protein 1 \| location=PbANKA_08_v3:1170535-1175910(+) \| protein_length=1791 \| sequence_SO= | 0 | 0 | 0 | 0 | 0 | 1 | 1 | 1 | 1 | 0 | 0 | 51 | 28 | 0 | 0 | 2 | 0 | 0 | 0 | 0 | 0 | 0 | 17 | 22 | 0 | 0 |
| Hemoglobin subunit alpha OS=Mus musculus OX=10090 GN=Hba PE=1 SV=2 | 2 | 0 | 0 | 5 | 1 | 2 | 2 | 3 | 3 | 4 | 3 | 14 | 5 | 1 | 0 | 5 | 5 | 1 | 4 | 1 | 1 | 1 | 8 | 14 | 2 | 2 |
| \| transcript=PBANKA_1229000.1 \| gene=PBANKA_1229000 \| organism=Plasmodium_berghei_ANKA \| gene_product=Plasmodium exported protein (PHIST), unknown function \| transcript_product=Plasmodium exported protein (PHIST), unknown function \| location=PbANKA_12_v3:112 | 9 | 4 | 7 | 17 | 7 | 0 | 0 | 0 | 0 | 15 | 6 | 17 | 1 | 0 | 0 | 0 | 2 | 5 | 3 | 0 | 7 | 5 | 0 | 9 | 0 | 0 |
| Keratin, type I cytoskeletal 14 OS=Mus musculus OX=10090 GN=Krt14 PE=1 SV=2 | 0 | 0 | 15 | 6 | 3 | 0 | 0 | 0 | 0 | 14 | 12 | 12 | 8 | 0 | 0 | 0 | 0 | 0 | 4 | 0 | 0 | 5 | 5 | 8 | 15 | 8 |
| \| transcript=PBANKA_0623300.1 \| gene=PBANKA_0623300 \| organism=Plasmodium_berghei_ANKA \| gene_product=tryptophan-rich protein \| transcript_product=tryptophan-rich protein \| location=PbANKA_06_v3:951400-952526(-) \| protein_length=342 \| sequence_SO=chromosome | 10 | 2 | 4 | 11 | 6 | 0 | 0 | 0 | 0 | 2 | 6 | 11 | 1 | 0 | 0 | 1 | 1 | 5 | 2 | 0 | 5 | 4 | 0 | 8 | 0 | 0 |
| Keratin, type I cytoskeletal 17 OS=Mus musculus OX=10090 GN=Krt17 PE=1 SV=3 | 7 | 5 | 0 | 0 | 0 | 7 | 8 | 4 | 4 | 0 | 0 | 0 | 0 | 8 | 6 | 12 | 11 | 0 | 0 | 0 | 16 | 0 | 0 | 0 | 13 | 8 |
| Talin-1 OS=Mus musculus OX=10090 GN=Tln1 PE=1 SV=2 | 0 | 0 | 0 | 12 | 0 | 2 | 0 | 0 | 1 | 0 | 0 | 5 | 0 | 0 | 0 | 0 | 45 | 0 | 0 | 0 | 1 | 0 | 0 | 12 | 0 | 0 |
| Hemoglobin subunit beta-2 OS=Mus musculus OX=10090 GN=Hbb-b2 PE=1 SV=2 | 6 | 0 | 0 | 6 | 4 | 0 | 0 | 0 | 0 | 4 | 3 | 14 | 3 | 0 | 0 | 5 | 9 | 1 | 2 | 3 | 2 | 2 | 6 | 9 | 2 | 3 |
| \| transcript=PBANKA_0818900.1 \| gene=PBANKA_0818900 \| organism=Plasmodium_berghei_ANKA \| gene_product=endoplasmic reticulum chaperone BiP, putative \| transcript_product=endoplasmic reticulum chaperone BiP, putative \| location=PbANKA_08_v3:755408-757677(+) \| | 1 | 2 | 4 | 2 | 4 | 2 | 2 | 0 | 0 | 1 | 0 | 2 | 1 | 6 | 2 | 21 | 15 | 5 | 1 | 0 | 1 | 0 | 1 | 0 | 0 | 1 |
| Progranulin OS=Mus musculus OX=10090 GN=Grn PE=1 SV=2 | 3 | 3 | 3 | 5 | 7 | 2 | 3 | 1 | 3 | 0 | 0 | 0 | 0 | 5 | 0 | 5 | 0 | 7 | 2 | 0 | 5 | 6 | 0 | 0 | 0 | 0 |
| Keratin, type I cytoskeletal 16 OS=Mus musculus OX=10090 GN=Krt16 PE=1 SV=3 | 0 | 0 | 10 | 5 | 0 | 0 | 0 | 0 | 0 | 5 | 4 | 5 | 4 | 3 | 0 | 6 | 10 | 0 | 0 | 0 | 13 | 0 | 2 | 3 | 7 | 4 |

| | | | | | | | | | | | | | | | | | | | | | | | | | | |
|---|---|---|---|---|---|---|---|---|---|---|---|---|---|---|---|---|---|---|---|---|---|---|---|---|---|---|
| \| transcript=PBANKA_1133300.1 \| gene=PBANKA_1133300 \| organism=Plasmodium_berghei_ANKA \| gene_product=elongation factor 1-alpha \| transcript_product=elongation factor 1-alpha \| location=PbANKA_11_v3:1253535-1254866(-) \| protein_length=443 \| sequence_SO=chrom | 5 | 1 | 3 | 6 | 5 | 1 | 1 | 0 | 1 | 3 | 0 | 6 | 1 | 0 | 0 | 2 | 2 | 2 | 1 | 0 | 2 | 2 | 5 | 3 | 1 | 1 |
| \| transcript=PBANKA_0711900.1 \| gene=PBANKA_0711900 \| organism=Plasmodium_berghei_ANKA \| gene_product=heat shock protein 70 \| transcript_product=heat shock protein 70 \| location=PbANKA_07_v3:454582-456663(-) \| protein_length=693 \| sequence_SO=chromosome \| SO | 1 | 0 | 0 | 5 | 2 | 1 | 0 | 0 | 2 | 4 | 0 | 10 | 3 | 0 | 1 | 2 | 5 | 1 | 0 | 0 | 2 | 1 | 6 | 5 | 1 | 1 |
| Ig kappa chain V-III region PC 2880/PC 1229 OS=Mus musculus OX=10090 PE=1 SV=1 | 0 | 0 | 0 | 0 | 0 | 0 | 0 | 0 | 0 | 0 | 0 | 0 | 0 | 21 | 9 | 18 | 14 | 0 | 0 | 0 | 0 | 0 | 0 | 0 | 0 | 0 |
| \| transcript=PBANKA_0830200.1 \| gene=PBANKA_0830200 \| organism=Plasmodium_berghei_ANKA \| gene_product=high molecular weight rhoptry protein 2 \| transcript_product=high molecular weight rhoptry protein 2 \| location=PbANKA_08_v3:1145085-1149997(+) \| protein_le | 0 | 0 | 0 | 0 | 0 | 0 | 0 | 0 | 0 | 3 | 0 | 40 | 0 | 0 | 0 | 3 | 3 | 0 | 0 | 0 | 0 | 0 | 2 | 7 | 0 | 0 |
| Keratin, type II cytoskeletal 1b OS=Mus musculus OX=10090 GN=Krt77 PE=1 SV=1 | 3 | 0 | 5 | 4 | 0 | 8 | 8 | 0 | 0 | 4 | 3 | 4 | 0 | 0 | 0 | 0 | 8 | 0 | 2 | 0 | 6 | 4 | 0 | 0 | 7 | 3 |
| Histone H4 OS=Mus musculus OX=10090 GN=H4c1 PE=1 SV=2 | 0 | 1 | 0 | 0 | 5 | 9 | 2 | 4 | 2 | 0 | 0 | 5 | 1 | 1 | 3 | 3 | 5 | 2 | 0 | 0 | 4 | 3 | 1 | 0 | 4 | 3 |
| Protein 4.2 OS=Mus musculus OX=10090 GN=Epb42 PE=1 SV=3 | 6 | 0 | 0 | 4 | 0 | 0 | 0 | 1 | 0 | 0 | 0 | 5 | 1 | 1 | 2 | 9 | 21 | 0 | 0 | 0 | 0 | 0 | 0 | 0 | 0 | 0 |
| Ig gamma-2A chain C region, A allele OS=Mus musculus OX=10090 GN=Ighg PE=1 SV=1 | 1 | 0 | 0 | 0 | 0 | 0 | 0 | 0 | 0 | 1 | 1 | 20 | 4 | 1 | 0 | 1 | 0 | 0 | 0 | 0 | 1 | 0 | 4 | 10 | 0 | 0 |
| Glyceraldehyde-3-phosphate dehydrogenase OS=Mus musculus OX=10090 GN=Gapdh PE=1 SV=2 | 1 | 2 | 1 | 2 | 0 | 1 | 0 | 0 | 0 | 0 | 0 | 4 | 1 | 2 | 1 | 14 | 8 | 0 | 0 | 0 | 4 | 0 | 0 | 2 | 3 | 0 |
| Filamin-A OS=Mus musculus OX=10090 GN=Flna PE=1 SV=5 | 0 | 0 | 0 | 0 | 0 | 8 | 0 | 8 | 1 | 0 | 0 | 1 | 0 | 0 | 0 | 0 | 28 | 0 | 0 | 0 | 0 | 0 | 0 | 5 | 0 | 0 |
| \| transcript=PBANKA_0941900.1 \| gene=PBANKA_0941900 \| organism=Plasmodium_berghei_ANKA \| gene_product=histone H4, putative \| transcript_product=histone H4, putative \| location=PbANKA_09_v3:1513619-1513930(+) \| protein_length=103 \| sequence_SO=chromosome \| SO | 2 | 0 | 0 | 0 | 8 | 8 | 0 | 6 | 0 | 2 | 0 | 3 | 0 | 0 | 0 | 0 | 5 | 6 | 2 | 1 | 2 | 0 | 1 | 1 | 0 | 0 |
| \| transcript=PBANKA_1423600.1 \| gene=PBANKA_1423600 \| organism=Plasmodium_berghei_ANKA \| gene_product=40S ribosomal protein S16, putative \| transcript_product=40S ribosomal protein S16, putative \| location=PbANKA_14_v3: 913169-914008(+) \| protein_length=144 \| | 4 | 2 | 2 | 1 | 3 | 0 | 0 | 0 | 0 | 3 | 2 | 3 | 2 | 0 | 0 | 1 | 0 | 3 | 1 | 0 | 3 | 4 | 2 | 1 | 3 | 4 |
| Ig gamma-2B chain C region OS=Mus musculus OX=10090 GN=Igh-3 PE=1 SV=3 | 0 | 0 | 0 | 1 | 0 | 0 | 0 | 0 | 0 | 5 | 2 | 12 | 2 | 3 | 0 | 3 | 0 | 0 | 0 | 0 | 2 | 0 | 3 | 9 | 0 | 0 |
| Immunoglobulin heavy constant mu OS=Mus musculus OX=10090 GN=Ighm PE=1 SV=2 | 0 | 0 | 0 | 1 | 1 | 0 | 0 | 0 | 0 | 3 | 1 | 11 | 8 | 0 | 0 | 0 | 0 | 2 | 0 | 0 | 0 | 0 | 5 | 8 | 0 | 0 |
| Inter-alpha-trypsin inhibitor heavy chain H2 OS=Mus musculus OX=10090 GN=Itih2 PE=1 SV=1 | 3 | 3 | 3 | 4 | 2 | 0 | 0 | 0 | 0 | 2 | 1 | 7 | 3 | 0 | 0 | 0 | 0 | 1 | 3 | 2 | 3 | 0 | 2 | 9 | 0 | 0 |
| Desmoplakin OS=Mus musculus OX=10090 GN=Dsp PE=1 SV=1 | 1 | 0 | 0 | 2 | 0 | 0 | 0 | 0 | 0 | 3 | 11 | 2 | 2 | 0 | 0 | 0 | 0 | 0 | 0 | 0 | 5 | 0 | 0 | 1 | 7 | 1 |
| Fibrinogen gamma chain OS=Mus musculus OX=10090 GN=Fgg PE=1 SV=1 | 0 | 0 | 0 | 2 | 0 | 1 | 0 | 0 | 1 | 0 | 0 | 9 | 0 | 0 | 0 | 0 | 14 | 0 | 0 | 0 | 0 | 0 | 0 | 5 | 0 | 0 |
| Thrombospondin-1 OS=Mus musculus OX=10090 GN=Thbs1 PE=1 SV=1 | 1 | 0 | 2 | 2 | 2 | 1 | 0 | 0 | 0 | 0 | 0 | 4 | 0 | 0 | 0 | 0 | 21 | 0 | 1 | 0 | 2 | 0 | 0 | 6 | 0 | 0 |
| \| transcript=PBANKA_1400600.1 \| gene=PBANKA_1400600 \| organism=Plasmodium_berghei_ANKA \| gene_product=cytoadherence linked asexual protein, putative \| transcript_product=cytoadherence linked asexual protein, putative \| location=PbANKA_14_v3:75956-80552(-) \| | 0 | 0 | 0 | 0 | 0 | 0 | 0 | 0 | 0 | 0 | 0 | 36 | 0 | 0 | 0 | 2 | 1 | 0 | 0 | 0 | 0 | 0 | 1 | 2 | 0 | 0 |
| \| transcript=PBANKA_1326400.1 \| gene=PBANKA_1326400 \| organism=Plasmodium_berghei_ANKA \| gene_product=glyceraldehyde-3-phosphate dehydrogenase \| transcript_product=glyceraldehyde-3-phosphate dehydrogenase \| location=PbANKA_13_v3: 1039995-1041245(+) \| protein_ | 2 | 0 | 0 | 4 | 0 | 0 | 0 | 0 | 0 | 0 | 0 | 5 | 1 | 0 | 2 | 1 | 2 | 0 | 1 | 0 | 0 | 1 | 2 | 4 | 0 | 0 |
| Fibrinogen beta chain OS=Mus musculus OX=10090 GN=Fgb PE=1 SV=1 | 0 | 0 | 0 | 4 | 0 | 4 | 0 | 0 | 0 | 0 | 0 | 8 | 0 | 0 | 0 | 0 | 15 | 0 | 0 | 1 | 1 | 1 | 0 | 7 | 0 | 0 |
| Immunoglobulin heavy variable 5-4 (Fragment) OS=Mus musculus OX=10090 GN=Ighv5-4 PE=4 SV=1 | 0 | 0 | 0 | 0 | 0 | 0 | 0 | 0 | 0 | 1 | 0 | 4 | 1 | 11 | 6 | 11 | 8 | 0 | 0 | 0 | 0 | 0 | 0 | 3 | 0 | 0 |
| Fibrinogen alpha chain OS=Mus musculus OX=10090 GN=Fga PE=1 SV=1 | 0 | 0 | 0 | 1 | 1 | 2 | 0 | 1 | 0 | 1 | 0 | 3 | 1 | 0 | 0 | 0 | 14 | 0 | 0 | 1 | 1 | 0 | 0 | 2 | 0 | 0 |
| Complement C1q subcomponent subunit B OS=Mus musculus OX=10090 GN=C1qb PE=1 SV=2 | 0 | 0 | 0 | 0 | 0 | 0 | 0 | 0 | 0 | 6 | 2 | 17 | 4 | 1 | 0 | 1 | 0 | 0 | 0 | 0 | 0 | 0 | 7 | 7 | 0 | 0 |
| \| transcript=PBANKA_1102200.1 \| gene=PBANKA_1102200 \| organism=Plasmodium_berghei_ANKA \| gene_product=merozoite surface protein 8 \| transcript_product=merozoite surface protein 8 \| location=PbANKA_11_v3:111532-112761(-) \| protein_length=409 \| sequence_SO=chr | 0 | 0 | 0 | 0 | 0 | 0 | 0 | 0 | 0 | 0 | 0 | 13 | 6 | 0 | 0 | 0 | 0 | 0 | 0 | 0 | 0 | 0 | 5 | 9 | 0 | 0 |
| Histone H2B type 1-F/J/L OS=Mus musculus OX=10090 GN=H2bc7 PE=1 SV=2 | 1 | 0 | 0 | 0 | 1 | 5 | 0 | 2 | 0 | 0 | 0 | 3 | 2 | 0 | 0 | 0 | 5 | 0 | 0 | 0 | 1 | 0 | 1 | 0 | 3 | 3 |
| \| transcript=PBANKA_1224200.1 \| gene=PBANKA_1224200 \| organism=Plasmodium_berghei_ANKA \| gene_product=DnaJ protein, putative \| transcript_product=DnaJ protein, putative \| location=PbANKA_12_v3:916254-918704(+) \| protein_length=816 \| sequence_SO=chromosome \| | 4 | 0 | 0 | 5 | 1 | 0 | 0 | 0 | 0 | 3 | 1 | 6 | 1 | 0 | 0 | 0 | 0 | 0 | 0 | 0 | 0 | 1 | 1 | 2 | 0 | 0 |

| | | | | | | | | | | | | | | | | | | | | | | | | | | |
|---|---|---|---|---|---|---|---|---|---|---|---|---|---|---|---|---|---|---|---|---|---|---|---|---|---|---|
| Ig heavy chain V-III region J606 OS=Mus musculus OX=10090 PE=1 SV=1 | 0 | 0 | 0 | 0 | 0 | 0 | 0 | 0 | 0 | 5 | 3 | 11 | 3 | 0 | 0 | 0 | 0 | 0 | 0 | 0 | 0 | 0 | 2 | 8 | 0 | 0 |
| \| transcript=PBANKA_0930300.1 \| gene=PBANKA_0930300 \| organism=Plasmodium_berghei_ANKA \| gene_product=GTP-binding nuclear protein RAN/TC4, putative \| transcript_product=GTP-binding nuclear protein RAN/TC4, putative \| location=PbANKA_09_v3:1111332-1112130 (+) | 1 | 0 | 2 | 2 | 2 | 0 | 0 | 0 | 0 | 0 | 0 | 5 | 1 | 0 | 0 | 1 | 2 | 0 | 0 | 0 | 0 | 0 | 3 | 4 | 1 | 0 |
| \| transcript=PBANKA_1365500.1 \| gene=PBANKA_1365500 \| organism=Plasmodium_berghei_ANKA \| gene_product=exported protein IBIS1 \| transcript_product=exported protein IBIS1 \| location=PbANKA_13_v3:2490490-2491687(+) \| protein_length=327 \| sequence_SO=chromosome | 2 | 0 | 0 | 1 | 0 | 0 | 0 | 0 | 0 | 2 | 2 | 8 | 4 | 0 | 0 | 0 | 0 | 0 | 0 | 0 | 1 | 0 | 5 | 6 | 0 | 0 |
| Complement C3 OS=Mus musculus OX=10090 GN=C3 PE=1 SV=3 | 4 | 1 | 0 | 5 | 1 | 0 | 0 | 0 | 0 | 0 | 0 | 3 | 0 | 0 | 0 | 0 | 0 | 0 | 0 | 0 | 0 | 0 | 0 | 5 | 0 | 0 |
| \| transcript=PBANKA_0108800.1 \| gene=PBANKA_0108800 \| organism=Plasmodium_berghei_ANKA \| gene_product=histone H3, putative \| transcript_product=histone H3, putative \| location=PbANKA_01_v3:363203-363613(+) \| protein_length=136 \| sequence_SO=chromosome \| SO=p | 1 | 0 | 1 | 0 | 1 | 2 | 0 | 1 | 0 | 1 | 0 | 1 | 0 | 0 | 0 | 0 | 1 | 0 | 0 | 0 | 0 | 0 | 0 | 0 | 1 | 2 |
| \| transcript=PBANKA_1231000.1 \| gene=PBANKA_1231000 \| organism=Plasmodium_berghei_ANKA \| gene_product=40S ribosomal protein S11, putative \| transcript_product=40S ribosomal protein S11, putative \| location=PbANKA_12_v3: 1200699-1201422(-) \| protein_length=151 | 3 | 0 | 2 | 3 | 2 | 1 | 1 | 0 | 1 | 2 | 0 | 0 | 0 | 0 | 0 | 0 | 0 | 1 | 1 | 0 | 2 | 1 | 0 | 0 | 0 | 0 |
| Gelsolin OS=Mus musculus OX=10090 GN=Gsn PE=1 SV=3 | 3 | 0 | 0 | 5 | 0 | 0 | 0 | 0 | 0 | 1 | 0 | 5 | 0 | 0 | 0 | 0 | 0 | 0 | 0 | 0 | 0 | 0 | 1 | 4 | 0 | 0 |
| \| transcript=PBANKA_0416000.1 \| gene=PBANKA_0416000 \| organism=Plasmodium_berghei_ANKA \| gene_product=high molecular weight rhoptry protein 3, putative \| transcript_product=high molecular weight rhoptry protein 3, putative \| location=PbANKA_04_v3:574017-5775 | 0 | 0 | 0 | 0 | 0 | 0 | 0 | 0 | 0 | 1 | 0 | 18 | 0 | 0 | 0 | 3 | 2 | 0 | 0 | 0 | 0 | 0 | 0 | 1 | 0 | 0 |
| Keratin, type I cytoskeletal 19 OS=Mus musculus OX=10090 GN=Krt19 PE=1 SV=1 | 0 | 0 | 7 | 5 | 0 | 0 | 0 | 0 | 0 | 4 | 0 | 0 | 0 | 0 | 0 | 0 | 0 | 0 | 0 | 0 | 10 | 0 | 0 | 0 | 6 | 0 |
| \| transcript=PBANKA_0941800.1 \| gene=PBANKA_0941800 \| organism=Plasmodium_berghei_ANKA \| gene_product=histone H2B, putative \| transcript_product=histone H2B, putative \| location=PbANKA_09_v3:1510191-1510547(-) \| protein_length=118 \| sequence_SO=chromosome \| | 2 | 0 | 0 | 1 | 3 | 1 | 0 | 0 | 1 | 3 | 2 | 2 | 1 | 0 | 0 | 0 | 0 | 1 | 0 | 1 | 1 | 0 | 2 | 1 | 1 | 0 |
| Protein 4.1 OS=Mus musculus OX=10090 GN=Epb41 PE=1 SV=2 | 2 | 0 | 0 | 2 | 0 | 0 | 0 | 0 | 0 | 1 | 1 | 17 | 0 | 0 | 0 | 0 | 0 | 0 | 0 | 0 | 0 | 1 | 0 | 0 | 0 | 0 |
| Complement C1q subcomponent subunit A OS=Mus musculus OX=10090 GN=C1qa PE=1 SV=2 | 0 | 0 | 0 | 0 | 0 | 0 | 0 | 0 | 0 | 7 | 3 | 10 | 1 | 0 | 0 | 0 | 0 | 0 | 0 | 0 | 0 | 0 | 2 | 4 | 0 | 0 |
| \| transcript=PBANKA_0941500.1 \| gene=PBANKA_0941500 \| organism=Plasmodium_berghei_ANKA \| gene_product=40S ribosomal protein S4, putative \| transcript_product=40S ribosomal protein S4, putative \| location=PbANKA_09_v3: 1499958-1501019(+) \| protein_length=261 \| | 1 | 0 | 0 | 0 | 0 | 0 | 0 | 0 | 0 | 5 | 3 | 2 | 2 | 0 | 0 | 0 | 0 | 0 | 0 | 0 | 2 | 0 | 1 | 1 | 1 | 0 |
| \| transcript=PBANKA_1459300.1 \| gene=PBANKA_1459300 \| organism=Plasmodium_berghei_ANKA \| gene_product=actin I \| transcript_product=actin I \| location=PbANKA_14_v3: 2267380-2268510(+) \| protein_length=376 \| sequence_SO=chromosome \| SO=protein_coding \| is_pseud | 1 | 0 | 0 | 1 | 2 | 0 | 0 | 0 | 0 | 0 | 0 | 6 | 1 | 0 | 1 | 0 | 1 | 1 | 1 | 0 | 2 | 1 | 0 | 0 | 0 | 0 |
| \| transcript=PBANKA_1145800.1 \| gene=PBANKA_1145800 \| organism=Plasmodium_berghei_ANKA \| gene_product=membrane associated histidine-rich protein 1a \| transcript_product=membrane associated histidine-rich protein 1a \| location=PbANKA_11_v3:1714536-1715328(+) | 2 | 0 | 2 | 2 | 0 | 0 | 0 | 0 | 0 | 2 | 2 | 5 | 3 | 0 | 0 | 0 | 0 | 0 | 0 | 0 | 2 | 2 | 3 | 4 | 0 | 0 |
| Periostin OS=Mus musculus OX=10090 GN=Postn PE=1 SV=2 | 2 | 0 | 2 | 8 | 4 | 0 | 0 | 0 | 0 | 0 | 0 | 0 | 0 | 0 | 0 | 0 | 0 | 0 | 0 | 0 | 1 | 0 | 0 | 1 | 0 | 0 |
| Keratin, type I cytoskeletal 15 OS=Ovis aries GN=KRT15 PE=2 SV=1 | 0 | 0 | 0 | 0 | 0 | 0 | 9 | 4 | 0 | 0 | 0 | 0 | 4 | 0 | 0 | 0 | 0 | 0 | 0 | 0 | 0 | 0 | 0 | 0 | 0 | 0 |
| Keratin, type I microfibrillar 48 kDa, component 8C-1 OS=Ovis aries PE=1 SV=2 | 0 | 0 | 0 | 0 | 0 | 0 | 0 | 0 | 0 | 0 | 0 | 0 | 0 | 0 | 0 | 0 | 0 | 0 | 0 | 0 | 23 | 0 | 0 | 0 | 0 | 0 |
| Junction plakoglobin OS=Mus musculus OX=10090 GN=Jup PE=1 SV=3 | 0 | 0 | 0 | 0 | 0 | 0 | 0 | 0 | 0 | 1 | 5 | 3 | 1 | 0 | 0 | 0 | 1 | 0 | 0 | 0 | 4 | 0 | 0 | 3 | 1 | 0 |
| Heat shock 70 kDa protein 1-like OS=Mus musculus OX=10090 GN=Hspa1l PE=1 SV=4 | 3 | 0 | 0 | 0 | 0 | 0 | 0 | 0 | 0 | 0 | 1 | 4 | 2 | 1 | 0 | 4 | 0 | 0 | 0 | 1 | 1 | 1 | 3 | 3 | 1 | 0 |
| \| transcript=PBANKA_1210200.1 \| gene=PBANKA_1210200 \| organism=Plasmodium_berghei_ANKA \| gene_product=PRE-binding protein, putative \| transcript_product=PRE-binding protein, putative \| location=PbANKA_12_v3:379958-382870 (+) \| protein_length=970 \| sequence_SO | 0 | 0 | 0 | 0 | 0 | 0 | 0 | 0 | 0 | 11 | 8 | 0 | 0 | 0 | 0 | 0 | 0 | 0 | 0 | 0 | 0 | 0 | 0 | 5 | 0 | 0 |
| \| transcript=PBANKA_1200600.1 \| gene=PBANKA_1200600 \| organism=Plasmodium_berghei_ANKA \| gene_product=Plasmodium exported protein, unknown function \| transcript_product=Plasmodium exported protein, unknown function \| location=PbANKA_12_v3:49909-56662 (-) \| pr | 0 | 0 | 1 | 1 | 0 | 0 | 1 | 0 | 0 | 1 | 1 | 6 | 1 | 1 | 0 | 1 | 4 | 0 | 0 | 0 | 1 | 0 | 1 | 1 | 0 | 0 |
| von Willebrand factor OS=Mus musculus OX=10090 GN=Vwf PE=1 SV=2 | 0 | 0 | 0 | 0 | 0 | 5 | 0 | 3 | 0 | 0 | 0 | 0 | 0 | 0 | 0 | 0 | 12 | 0 | 0 | 0 | 0 | 0 | 0 | 0 | 0 | 0 |
| Keratin, type I cuticular Ha3-II OS=Homo sapiens GN=KRT33B PE=2 SV=3 | 0 | 0 | 0 | 0 | 0 | 4 | 0 | 0 | 0 | 0 | 0 | 0 | 0 | 0 | 0 | 3 | 0 | 0 | 0 | 0 | 17 | 0 | 0 | 0 | 0 | 0 |
| Keratin, type I microfibrillar, 47.6 kDa OS=Ovis aries PE=3 SV=2 | 0 | 0 | 0 | 0 | 0 | 0 | 0 | 0 | 0 | 0 | 0 | 0 | 0 | 0 | 0 | 0 | 0 | 0 | 0 | 0 | 22 | 0 | 0 | 0 | 0 | 0 |

| | | | | | | | | | | | | | | | | | | | | | | | | | | |
|---|---|---|---|---|---|---|---|---|---|---|---|---|---|---|---|---|---|---|---|---|---|---|---|---|---|---|
| Histone H2A type 1-B OS=Mus musculus OX=10090 GN=H2ac4 PE=1 SV=1 | 0 | 0 | 0 | 1 | 0 | 2 | 0 | 1 | 0 | 0 | 0 | 0 | 0 | 0 | 0 | 0 | 0 | 0 | 0 | 0 | 1 | 0 | 0 | 0 | 0 | 0 |
| \| transcript=PBANKA_1407600.1 \| gene=PBANKA_1407600 \| organism=Plasmodium_berghei_ANKA \| gene_product=60S ribosomal protein L24, putative \| transcript_product=60S ribosomal protein L24, putative \| location=PbANKA_14_v3:358003-359152(+) \| protein_length=163 \| | 3 | 0 | 1 | 2 | 3 | 0 | 0 | 0 | 0 | 0 | 0 | 0 | 0 | 0 | 0 | 0 | 0 | 1 | 0 | 0 | 2 | 0 | 0 | 0 | 0 | 0 |
| Inter alpha-trypsin inhibitor, heavy chain 4 OS=Mus musculus OX=10090 GN=Itih4 PE=1 SV=2 | 0 | 0 | 0 | 3 | 0 | 0 | 0 | 0 | 0 | 0 | 0 | 0 | 0 | 0 | 0 | 0 | 0 | 0 | 0 | 0 | 1 | 0 | 0 | 4 | 0 | 0 |
| \| transcript=PBANKA_0511400.1 \| gene=PBANKA_0511400 \| organism=Plasmodium_berghei_ANKA \| gene_product=peroxiredoxin, putative \| transcript_product=peroxiredoxin, putative \| location=PbANKA_05_v3:457605-459183(-) \| protein_length=389 \| sequence_SO=chromosome | 0 | 0 | 0 | 0 | 1 | 0 | 0 | 0 | 0 | 4 | 3 | 5 | 3 | 0 | 0 | 0 | 0 | 0 | 0 | 0 | 0 | 0 | 2 | 3 | 0 | 0 |
| Complement C1q subcomponent subunit C OS=Mus musculus OX=10090 GN=C1qc PE=1 SV=2 | 0 | 0 | 0 | 0 | 0 | 0 | 0 | 0 | 0 | 1 | 2 | 8 | 0 | 0 | 0 | 1 | 0 | 0 | 0 | 0 | 0 | 0 | 3 | 3 | 0 | 0 |
| Ig kappa chain V19-17 OS=Mus musculus OX=10090 GN=Igk-V19-17 PE=1 SV=1 | 0 | 0 | 0 | 0 | 0 | 0 | 0 | 0 | 0 | 3 | 0 | 6 | 2 | 0 | 0 | 0 | 0 | 0 | 0 | 0 | 0 | 0 | 1 | 1 | 0 | 1 |
| Actin, aortic smooth muscle OS=Mus musculus OX=10090 GN=Acta2 PE=1 SV=1 | 0 | 0 | 0 | 0 | 0 | 0 | 1 | 0 | 0 | 0 | 0 | 0 | 0 | 0 | 15 | 0 | 0 | 0 | 0 | 0 | 0 | 0 | 0 | 0 | 0 | 0 |
| Albumin OS=Mus musculus OX=10090 GN=Alb PE=1 SV=3 | 0 | 0 | 1 | 2 | 0 | 0 | 0 | 0 | 0 | 0 | 0 | 4 | 1 | 1 | 1 | 1 | 2 | 1 | 1 | 1 | 1 | 1 | 1 | 1 | 0 | 0 |
| Histone H3.3C OS=Mus musculus OX=10090 GN=H3-5 PE=3 SV=3 | 0 | 0 | 0 | 0 | 0 | 2 | 0 | 0 | 0 | 0 | 0 | 4 | 0 | 1 | 0 | 1 | 0 | 0 | 0 | 0 | 0 | 0 | 0 | 0 | 0 | 0 |
| \| transcript=PBANKA_1000600.1 \| gene=PBANKA_1000600 \| organism=Plasmodium_berghei_ANKA \| gene_product=erythrocyte membrane antigen 1 \| transcript_product=erythrocyte membrane antigen 1 \| location=PbANKA_10_v3:65304-66403(+) \| protein_length=319 \| sequence_SO | 1 | 0 | 0 | 1 | 0 | 0 | 0 | 0 | 0 | 1 | 0 | 10 | 1 | 0 | 0 | 1 | 2 | 0 | 0 | 0 | 0 | 0 | 0 | 2 | 0 | 0 |
| \| transcript=PBANKA_1237800.1 \| gene=PBANKA_1237800 \| organism=Plasmodium_berghei_ANKA \| gene_product=multidrug resistance protein 1, putative \| transcript_product=multidrug resistance protein 1, putative \| location=PbANKA_12_v3:1452481-1456740(+) \| protein_ | 1 | 0 | 0 | 4 | 0 | 0 | 0 | 0 | 0 | 0 | 0 | 7 | 1 | 0 | 0 | 0 | 0 | 0 | 0 | 0 | 1 | 1 | 0 | 1 | 0 | 0 |
| ATP synthase subunit beta, mitochondrial OS=Mus musculus OX=10090 GN=Atp5f1b PE=1 SV=2 | 0 | 0 | 0 | 0 | 0 | 0 | 0 | 0 | 0 | 3 | 4 | 2 | 1 | 0 | 0 | 0 | 0 | 0 | 0 | 0 | 1 | 0 | 4 | 2 | 0 | 0 |
| \| transcript=PBANKA_0918100.1 \| gene=PBANKA_0918100 \| organism=Plasmodium_berghei_ANKA \| gene_product=60S ribosomal protein L38, putative \| transcript_product=60S ribosomal protein L38, putative \| location=PbANKA_09_v3:679335-679580(+) \| protein_length=81 \| | 0 | 0 | 1 | 0 | 0 | 0 | 0 | 0 | 0 | 4 | 5 | 0 | 1 | 0 | 0 | 0 | 0 | 0 | 0 | 0 | 1 | 0 | 2 | 0 | 0 | 0 |
| \| transcript=PBANKA_1437300.1 \| gene=PBANKA_1437300 \| organism=Plasmodium_berghei_ANKA \| gene_product=endoplasmin, putative \| transcript_product=endoplasmin, putative \| location=PbANKA_14_v3:1403252-1405678(-) \| protein_length=808 \| sequence_SO=chromosome \| | 0 | 0 | 0 | 0 | 0 | 0 | 0 | 0 | 0 | 0 | 0 | 0 | 0 | 2 | 1 | 11 | 4 | 0 | 0 | 0 | 0 | 0 | 0 | 0 | 0 | 0 |
| Keratin, type II microfibrillar, component 7C OS=Ovis aries PE=1 SV=1 | 0 | 0 | 0 | 0 | 0 | 0 | 0 | 0 | 0 | 0 | 0 | 0 | 0 | 0 | 0 | 0 | 0 | 0 | 0 | 0 | 18 | 0 | 0 | 0 | 0 | 0 |
| Inter-alpha-trypsin inhibitor heavy chain H3 OS=Mus musculus OX=10090 GN=Itih3 PE=1 SV=3 | 1 | 0 | 0 | 1 | 0 | 0 | 0 | 0 | 0 | 1 | 1 | 3 | 0 | 0 | 0 | 0 | 0 | 0 | 0 | 0 | 2 | 0 | 0 | 1 | 0 | 0 |
| \| transcript=PBANKA_1028400.1 \| gene=PBANKA_1028400 \| organism=Plasmodium_berghei_ANKA \| gene_product=60S ribosomal protein L10, putative \| transcript_product=60S ribosomal protein L10, putative \| location=PbANKA_10_v3:1159709-1160368(-) \| protein_length=219 | 1 | 1 | 0 | 1 | 2 | 0 | 0 | 0 | 0 | 0 | 0 | 3 | 2 | 0 | 0 | 0 | 1 | 0 | 0 | 0 | 1 | 0 | 2 | 3 | 1 | 0 |
| Ig heavy chain V region 441 OS=Mus musculus OX=10090 PE=4 SV=1 | 0 | 0 | 0 | 0 | 0 | 0 | 0 | 0 | 0 | 2 | 4 | 3 | 3 | 0 | 0 | 0 | 0 | 0 | 0 | 0 | 0 | 0 | 2 | 3 | 0 | 0 |
| \| transcript=PBANKA_1420300.1 \| gene=PBANKA_1420300 \| organism=Plasmodium_berghei_ANKA \| gene_product=conserved Plasmodium protein, unknown function \| transcript_product=conserved Plasmodium protein, unknown function \| location=PbANKA_14_v3:806316-807930(-) | 1 | 1 | 1 | 3 | 0 | 0 | 0 | 0 | 0 | 1 | 0 | 2 | 1 | 0 | 0 | 0 | 0 | 3 | 0 | 0 | 1 | 1 | 0 | 1 | 0 | 1 |
| Elongation factor 1-alpha 1 OS=Mus musculus OX=10090 GN=Eef1a1 PE=1 SV=3 | 0 | 0 | 0 | 0 | 0 | 0 | 0 | 0 | 0 | 0 | 1 | 4 | 0 | 0 | 1 | 0 | 0 | 0 | 0 | 0 | 2 | 0 | 1 | 2 | 4 | 1 |
| \| transcript=PBANKA_1223800.1 \| gene=PBANKA_1223800 \| organism=Plasmodium_berghei_ANKA \| gene_product=V-type proton ATPase subunit a, putative \| transcript_product=V-type proton ATPase subunit a, putative \| location=PbANKA_12_v3:898046-901101(+) \| protein_le | 0 | 0 | 0 | 1 | 4 | 0 | 0 | 0 | 0 | 0 | 0 | 1 | 0 | 0 | 0 | 0 | 0 | 4 | 0 | 0 | 0 | 0 | 1 | 0 | 0 | 0 |
| Alpha-enolase OS=Mus musculus OX=10090 GN=Eno1 PE=1 SV=3 | 0 | 0 | 0 | 2 | 0 | 0 | 0 | 0 | 0 | 0 | 0 | 0 | 0 | 0 | 2 | 1 | 0 | 0 | 0 | 0 | 2 | 0 | 0 | 1 | 0 | 0 |
| Ig lambda-1 chain C region OS=Mus musculus OX=10090 PE=1 SV=1 | 0 | 0 | 0 | 0 | 0 | 0 | 0 | 0 | 0 | 2 | 2 | 6 | 1 | 0 | 0 | 0 | 0 | 0 | 0 | 0 | 0 | 0 | 1 | 4 | 0 | 0 |
| Pigment epithelium-derived factor OS=Mus musculus OX=10090 GN=Serpinf1 PE=1 SV=2 | 2 | 1 | 1 | 3 | 1 | 0 | 0 | 0 | 0 | 0 | 0 | 2 | 0 | 0 | 0 | 0 | 0 | 0 | 0 | 1 | 1 | 0 | 1 | 4 | 0 | 0 |
| Ig heavy chain V region MOPC 47A OS=Mus musculus OX=10090 PE=1 SV=1 | 1 | 0 | 1 | 1 | 1 | 0 | 0 | 0 | 0 | 0 | 1 | 3 | 1 | 0 | 0 | 0 | 0 | 0 | 0 | 0 | 1 | 0 | 1 | 2 | 0 | 0 |
| \| transcript=PBANKA_1135100.1 \| gene=PBANKA_1135100 \| organism=Plasmodium_berghei_ANKA \| gene_product=40S ribosomal protein S15, putative \| transcript_product=40S ribosomal protein S15, putative \| location=PbANKA_11_v3:1315094-1315549(+) \| protein_length=151 | 4 | 0 | 0 | 0 | 1 | 0 | 0 | 0 | 0 | 1 | 0 | 1 | 1 | 0 | 0 | 0 | 0 | 0 | 0 | 0 | 0 | 0 | 0 | 0 | 0 | 0 |

<table>
<tr><td>| transcript=PBANKA_1401300.1 | gene=PBANKA_1401300 | organism=Plasmodium_berghei_ANKA | gene_product=40S ribosomal protein S7, putative | transcript_product=40S ribosomal protein S7, putative | location=PbANKA_14_v3: 108464-109231(-) | protein_length=194 | s</td><td>0</td><td>0</td><td>0</td><td>0</td><td>0</td><td>0</td><td>0</td><td>0</td><td>0</td><td>5</td><td>4</td><td>0</td><td>0</td><td>0</td><td>0</td><td>1</td><td>0</td><td>0</td><td>0</td><td>0</td><td>0</td><td>0</td><td>2</td><td>0</td><td>0</td><td>0</td></tr>
<tr><td>| transcript=PBANKA_0523700.1 | gene=PBANKA_0523700 | organism=Plasmodium_berghei_ANKA | gene_product=conserved Plasmodium protein, unknown function | transcript_product=conserved Plasmodium protein, unknown function | location=PbANKA_05_v3: 874814-875812(-)</td><td>0</td><td>0</td><td>0</td><td>1</td><td>2</td><td>0</td><td>0</td><td>0</td><td>0</td><td>2</td><td>0</td><td>1</td><td>2</td><td>0</td><td>0</td><td>0</td><td>0</td><td>1</td><td>0</td><td>0</td><td>0</td><td>0</td><td>0</td><td>1</td><td>0</td><td>0</td></tr>
<tr><td>| transcript=PBANKA_1423500.1 | gene=PBANKA_1423500 | organism=Plasmodium_berghei_ANKA | gene_product=60S ribosomal protein L13-2, putative | transcript_product=60S ribosomal protein L13-2, putative | location=PbANKA_14_v3: 910790-911537(-) | protein_length=2</td><td>3</td><td>0</td><td>0</td><td>1</td><td>0</td><td>0</td><td>0</td><td>0</td><td>0</td><td>0</td><td>0</td><td>0</td><td>0</td><td>0</td><td>0</td><td>0</td><td>0</td><td>0</td><td>1</td><td>0</td><td>0</td><td>0</td><td>0</td><td>0</td><td>1</td><td>0</td></tr>
<tr><td>Keratin, type II microfibrillar, component 5 OS=Ovis aries PE=1 SV=1</td><td>0</td><td>0</td><td>0</td><td>0</td><td>0</td><td>0</td><td>0</td><td>0</td><td>0</td><td>0</td><td>0</td><td>0</td><td>0</td><td>0</td><td>0</td><td>0</td><td>0</td><td>0</td><td>0</td><td>0</td><td>15</td><td>0</td><td>0</td><td>0</td><td>0</td><td>0</td></tr>
<tr><td>| transcript=PBANKA_0100600.1 | gene=PBANKA_0100600 | organism=Plasmodium_berghei_ANKA | gene_product=schizont membrane associated cytoadherence protein | transcript_product=schizont membrane associated cytoadherence protein | location=PbANKA_01_v3:42789-432</td><td>0</td><td>0</td><td>0</td><td>0</td><td>0</td><td>0</td><td>0</td><td>0</td><td>0</td><td>0</td><td>0</td><td>0</td><td>0</td><td>0</td><td>0</td><td>0</td><td>0</td><td>0</td><td>1</td><td>0</td><td>4</td><td>5</td><td>0</td><td>0</td><td>0</td><td>0</td></tr>
<tr><td>Keratin, type II cuticular Hb6 OS=Homo sapiens GN=KRT86 PE=1 SV=1</td><td>1</td><td>0</td><td>0</td><td>0</td><td>0</td><td>0</td><td>0</td><td>0</td><td>0</td><td>0</td><td>0</td><td>0</td><td>0</td><td>0</td><td>0</td><td>0</td><td>0</td><td>0</td><td>0</td><td>0</td><td>13</td><td>0</td><td>0</td><td>0</td><td>0</td><td>0</td></tr>
<tr><td>Keratin, type I cuticular Ha3-II OS=Mus musculus OX=10090 GN=Krt33b PE=1 SV=2</td><td>0</td><td>0</td><td>0</td><td>0</td><td>0</td><td>0</td><td>0</td><td>0</td><td>0</td><td>0</td><td>0</td><td>0</td><td>0</td><td>0</td><td>0</td><td>0</td><td>0</td><td>0</td><td>0</td><td>0</td><td>15</td><td>0</td><td>0</td><td>0</td><td>0</td><td>0</td></tr>
<tr><td>Prothrombin OS=Mus musculus OX=10090 GN=F2 PE=1 SV=1</td><td>1</td><td>1</td><td>1</td><td>2</td><td>1</td><td>0</td><td>0</td><td>0</td><td>0</td><td>0</td><td>0</td><td>1</td><td>0</td><td>0</td><td>0</td><td>0</td><td>0</td><td>0</td><td>0</td><td>0</td><td>0</td><td>0</td><td>0</td><td>1</td><td>0</td><td>0</td></tr>
<tr><td>| transcript=PBANKA_0709100.1 | gene=PBANKA_0709100 | organism=Plasmodium_berghei_ANKA | gene_product=60S ribosomal protein L22, putative | transcript_product=60S ribosomal protein L22, putative | location=PbANKA_07_v3: 355109-355741(-) | protein_length=138 |</td><td>0</td><td>0</td><td>0</td><td>0</td><td>0</td><td>0</td><td>0</td><td>0</td><td>0</td><td>3</td><td>2</td><td>0</td><td>2</td><td>0</td><td>0</td><td>0</td><td>0</td><td>0</td><td>0</td><td>0</td><td>0</td><td>0</td><td>2</td><td>0</td><td>1</td><td>0</td></tr>
<tr><td>Pregnancy zone protein OS=Mus musculus OX=10090 GN=Pzp PE=1 SV=3</td><td>0</td><td>0</td><td>0</td><td>0</td><td>0</td><td>0</td><td>0</td><td>0</td><td>0</td><td>0</td><td>0</td><td>8</td><td>0</td><td>0</td><td>0</td><td>0</td><td>0</td><td>0</td><td>0</td><td>0</td><td>0</td><td>0</td><td>0</td><td>1</td><td>0</td><td>0</td></tr>
<tr><td>Complement C4-B OS=Mus musculus OX=10090 GN=C4b PE=1 SV=3</td><td>1</td><td>0</td><td>0</td><td>1</td><td>0</td><td>0</td><td>0</td><td>0</td><td>0</td><td>1</td><td>0</td><td>2</td><td>1</td><td>1</td><td>0</td><td>1</td><td>0</td><td>0</td><td>0</td><td>0</td><td>0</td><td>0</td><td>1</td><td>1</td><td>0</td><td>0</td></tr>
<tr><td>| transcript=PBANKA_0715500.1 | gene=PBANKA_0715500 | organism=Plasmodium_berghei_ANKA | gene_product=phosphoglucomutase-2 | transcript_product=phosphoglucomutase-2 | location=PbANKA_07_v3:574761-575657(+) | protein_length=298 | sequence_SO=chromosome | SO=p</td><td>1</td><td>1</td><td>1</td><td>1</td><td>2</td><td>0</td><td>0</td><td>0</td><td>0</td><td>0</td><td>0</td><td>0</td><td>0</td><td>0</td><td>0</td><td>0</td><td>0</td><td>1</td><td>0</td><td>0</td><td>1</td><td>1</td><td>0</td><td>0</td><td>0</td><td>0</td></tr>
<tr><td>| transcript=PBANKA_0405300.1 | gene=PBANKA_0405300 | organism=Plasmodium_berghei_ANKA | gene_product=40S ribosomal protein S23, putative | transcript_product=40S ribosomal protein S23, putative | location=PbANKA_04_v3: 193055-194284(+) | protein_length=145 |</td><td>2</td><td>1</td><td>1</td><td>1</td><td>2</td><td>0</td><td>0</td><td>0</td><td>0</td><td>1</td><td>0</td><td>0</td><td>0</td><td>0</td><td>0</td><td>0</td><td>0</td><td>0</td><td>0</td><td>0</td><td>1</td><td>1</td><td>0</td><td>0</td><td>0</td><td>0</td></tr>
<tr><td>Immunoglobulin kappa chain variable 12-41 (Fragment) OS=Mus musculus OX=10090 GN=Igkv12-41 PE=1 SV=1</td><td>0</td><td>0</td><td>0</td><td>1</td><td>0</td><td>0</td><td>0</td><td>0</td><td>0</td><td>1</td><td>1</td><td>3</td><td>1</td><td>0</td><td>0</td><td>0</td><td>0</td><td>0</td><td>0</td><td>0</td><td>0</td><td>0</td><td>1</td><td>3</td><td>0</td><td>0</td></tr>
<tr><td>| transcript=PBANKA_1329300.1 | gene=PBANKA_1329300 | organism=Plasmodium_berghei_ANKA | gene_product=40S ribosomal protein S3, putative | transcript_product=40S ribosomal protein S3, putative | location=PbANKA_13_v3: 1185082-1186355(+) | protein_length=222 |</td><td>0</td><td>0</td><td>0</td><td>0</td><td>0</td><td>0</td><td>0</td><td>0</td><td>0</td><td>1</td><td>1</td><td>4</td><td>1</td><td>0</td><td>0</td><td>0</td><td>0</td><td>0</td><td>0</td><td>0</td><td>0</td><td>0</td><td>1</td><td>1</td><td>0</td><td>0</td></tr>
<tr><td>Predicted pseudogene 8797 OS=Mus musculus OX=10090 GN=Gm8797 PE=4 SV=1</td><td>0</td><td>0</td><td>0</td><td>2</td><td>1</td><td>0</td><td>0</td><td>0</td><td>0</td><td>0</td><td>0</td><td>1</td><td>0</td><td>0</td><td>0</td><td>1</td><td>2</td><td>0</td><td>1</td><td>0</td><td>3</td><td>0</td><td>0</td><td>1</td><td>0</td><td>0</td></tr>
<tr><td>| transcript=PBANKA_1324400.1 | gene=PBANKA_1324400 | organism=Plasmodium_berghei_ANKA | gene_product=60S ribosomal protein L27, putative | transcript_product=60S ribosomal protein L27, putative | location=PbANKA_13_v3: 979146-980299(-) | protein_length=146 |</td><td>0</td><td>0</td><td>0</td><td>1</td><td>0</td><td>0</td><td>0</td><td>0</td><td>0</td><td>1</td><td>1</td><td>1</td><td>2</td><td>0</td><td>0</td><td>0</td><td>0</td><td>0</td><td>0</td><td>0</td><td>0</td><td>0</td><td>1</td><td>2</td><td>0</td><td>0</td></tr>
<tr><td>| transcript=PBANKA_0316800.1 | gene=PBANKA_0316800 | organism=Plasmodium_berghei_ANKA | gene_product=erythrocyte membrane associated protein 2 | transcript_product=erythrocyte membrane associated protein 2 | location=PbANKA_03_v3:585675-586962(+) | protein_</td><td>2</td><td>0</td><td>0</td><td>3</td><td>0</td><td>0</td><td>0</td><td>0</td><td>0</td><td>0</td><td>0</td><td>1</td><td>0</td><td>0</td><td>0</td><td>1</td><td>1</td><td>0</td><td>0</td><td>0</td><td>0</td><td>0</td><td>0</td><td>0</td><td>0</td><td>0</td></tr>
<tr><td>| transcript=PBANKA_0414600.1 | gene=PBANKA_0414600 | organism=Plasmodium_berghei_ANKA | gene_product=40S ribosomal protein S15A, putative | transcript_product=40S ribosomal protein S15A, putative | location=PbANKA_04_v3: 520129-520638(+) | protein_length=130</td><td>1</td><td>0</td><td>0</td><td>0</td><td>0</td><td>0</td><td>0</td><td>0</td><td>0</td><td>2</td><td>1</td><td>1</td><td>1</td><td>0</td><td>0</td><td>0</td><td>0</td><td>0</td><td>0</td><td>0</td><td>0</td><td>0</td><td>0</td><td>0</td><td>0</td><td>0</td></tr>
<tr><td>| transcript=PBANKA_1234200.1 | gene=PBANKA_1234200 | organism=Plasmodium_berghei_ANKA | gene_product=40S ribosomal protein S24, putative | transcript_product=40S ribosomal protein S24, putative | location=PbANKA_12_v3: 1309151-1309654(-) | protein_length=133</td><td>2</td><td>0</td><td>0</td><td>0</td><td>1</td><td>0</td><td>0</td><td>0</td><td>0</td><td>2</td><td>2</td><td>0</td><td>0</td><td>0</td><td>0</td><td>0</td><td>0</td><td>0</td><td>0</td><td>0</td><td>1</td><td>0</td><td>0</td><td>0</td><td>0</td><td>0</td></tr>
<tr><td>Immunoglobulin G-binding protein A OS=Staphylococcus aureus (strain NCTC 8325) OX=93061 GN=spa PE=1 SV=3</td><td>0</td><td>0</td><td>0</td><td>0</td><td>0</td><td>0</td><td>0</td><td>0</td><td>0</td><td>1</td><td>1</td><td>0</td><td>0</td><td>0</td><td>0</td><td>0</td><td>0</td><td>0</td><td>0</td><td>0</td><td>0</td><td>0</td><td>0</td><td>1</td><td>7</td><td>3</td></tr>
<tr><td>| transcript=PBANKA_0836200.1 | gene=PBANKA_0836200 | organism=Plasmodium_berghei_ANKA | gene_product=conserved Plasmodium protein, unknown function | transcript_product=conserved Plasmodium protein, unknown function | location=PbANKA_08_v3: 1326359-1328673(+</td><td>2</td><td>0</td><td>0</td><td>2</td><td>0</td><td>0</td><td>0</td><td>0</td><td>0</td><td>0</td><td>0</td><td>3</td><td>0</td><td>0</td><td>0</td><td>0</td><td>0</td><td>0</td><td>0</td><td>0</td><td>2</td><td>1</td><td>0</td><td>0</td><td>0</td><td>0</td></tr>
</table>

| | | | | | | | | | | | | | | | | | | | | | | | | | | |
|---|---|---|---|---|---|---|---|---|---|---|---|---|---|---|---|---|---|---|---|---|---|---|---|---|---|---|
| Ig kappa chain V-V region HP R16.7 OS=Mus musculus OX=10090 PE=1 SV=1 | 0 | 0 | 0 | 0 | 0 | 0 | 0 | 0 | 0 | 0 | 0 | 5 | 1 | 0 | 0 | 0 | 0 | 0 | 0 | 0 | 0 | 0 | 0 | 3 | 0 | 0 |
| \| transcript=PBANKA_0300600.1 \| gene=PBANKA_0300600 \| organism=Plasmodium_berghei_ANKA \| gene_product=Plasmodium exported protein, unknown function \| transcript_product=Plasmodium exported protein, unknown function \| location=PbANKA_03_v3:19615-21889(-) \| pr | 0 | 0 | 0 | 0 | 0 | 0 | 0 | 0 | 0 | 0 | 0 | 6 | 1 | 0 | 0 | 0 | 0 | 0 | 0 | 0 | 0 | 0 | 1 | 4 | 0 | 0 |
| Beta-globin OS=Mus musculus OX=10090 GN=Hbb-bs PE=1 SV=1 | 0 | 0 | 0 | 0 | 0 | 0 | 0 | 0 | 0 | 0 | 0 | 0 | 0 | 0 | 1 | 7 | 8 | 0 | 0 | 0 | 0 | 0 | 0 | 0 | 0 | 0 |
| \| transcript=PBANKA_0906500.1 \| gene=PBANKA_0906500 \| organism=Plasmodium_berghei_ANKA \| gene_product=60S ribosomal protein L28, putative \| transcript_product=60S ribosomal protein L28, putative \| location=PbANKA_09_v3:212735-213540(+) \| protein_length=126 \| | 2 | 1 | 0 | 1 | 1 | 0 | 0 | 0 | 1 | 1 | 1 | 1 | 0 | 0 | 0 | 0 | 0 | 0 | 0 | 0 | 1 | 1 | 0 | 0 | 0 | 0 |
| \| transcript=PBANKA_0511900.1 \| gene=PBANKA_0511900 \| organism=Plasmodium_berghei_ANKA \| gene_product=60S ribosomal protein L3, putative \| transcript_product=60S ribosomal protein L3, putative \| location=PbANKA_05_v3: 468739-469899(-) \| protein_length=386 \| s | 0 | 0 | 0 | 0 | 0 | 0 | 0 | 0 | 0 | 1 | 0 | 2 | 1 | 0 | 0 | 0 | 0 | 0 | 0 | 0 | 0 | 0 | 1 | 0 | 0 | 0 |
| Vimentin OS=Mus musculus OX=10090 GN=Vim PE=1 SV=3 | 0 | 0 | 0 | 0 | 0 | 1 | 0 | 0 | 0 | 0 | 0 | 3 | 0 | 0 | 0 | 0 | 0 | 0 | 0 | 0 | 0 | 0 | 0 | 0 | 0 | 0 |
| \| transcript=PBANKA_0517000.1 \| gene=PBANKA_0517000 \| organism=Plasmodium_berghei_ANKA \| gene_product=early transcribed membrane protein \| transcript_product=early transcribed membrane protein \| location=PbANKA_05_v3: 628632-629573(+) \| protein_length=313 \| s | 0 | 0 | 0 | 0 | 0 | 0 | 0 | 0 | 0 | 3 | 1 | 0 | 0 | 0 | 0 | 3 | 2 | 0 | 0 | 0 | 0 | 0 | 0 | 0 | 0 | 0 |
| Ig heavy chain V region RF OS=Mus musculus OX=10090 PE=1 SV=1 | 0 | 0 | 0 | 0 | 0 | 0 | 0 | 0 | 0 | 1 | 1 | 4 | 2 | 0 | 0 | 0 | 0 | 0 | 0 | 0 | 0 | 0 | 2 | 3 | 0 | 0 |
| Dematin OS=Mus musculus OX=10090 GN=Dmtn PE=1 SV=1 | 1 | 0 | 0 | 1 | 0 | 0 | 0 | 0 | 0 | 0 | 0 | 5 | 0 | 0 | 0 | 0 | 1 | 0 | 0 | 0 | 0 | 0 | 0 | 0 | 0 | 0 |
| \| transcript=PBANKA_1117200.1 \| gene=PBANKA_1117200 \| organism=Plasmodium_berghei_ANKA \| gene_product=conserved Plasmodium protein, unknown function \| transcript_product=conserved Plasmodium protein, unknown function \| location=PbANKA_11_v3:651041-653497(+) | 0 | 0 | 0 | 0 | 0 | 0 | 0 | 0 | 0 | 0 | 0 | 2 | 0 | 0 | 0 | 0 | 0 | 0 | 0 | 0 | 0 | 0 | 0 | 0 | 0 | 0 |
| Histone H1.2 OS=Mus musculus OX=10090 GN=H1-2 PE=1 SV=2 | 0 | 0 | 0 | 0 | 0 | 8 | 0 | 0 | 0 | 1 | 0 | 0 | 0 | 0 | 0 | 0 | 1 | 0 | 0 | 0 | 0 | 0 | 0 | 0 | 2 | 1 |
| \| transcript=PBANKA_1101400.1 \| gene=PBANKA_1101400 \| organism=Plasmodium_berghei_ANKA \| gene_product=rhoptry-associated protein 2/3 \| transcript_product=rhoptry-associated protein 2/3 \| location=PbANKA_11_v3:86661-87827(-) \| protein_length=388 \| sequence_SO | 0 | 0 | 0 | 0 | 0 | 0 | 0 | 0 | 0 | 0 | 0 | 6 | 4 | 0 | 0 | 0 | 0 | 0 | 0 | 0 | 0 | 0 | 1 | 0 | 0 | 0 |
| Tropomyosin alpha-3 chain OS=Mus musculus OX=10090 GN=Tpm3 PE=1 SV=3 | 0 | 0 | 0 | 0 | 0 | 0 | 0 | 0 | 0 | 0 | 0 | 3 | 0 | 0 | 4 | 0 | 2 | 0 | 0 | 0 | 1 | 0 | 0 | 0 | 0 | 0 |
| \| transcript=PBANKA_1022000.1 \| gene=PBANKA_1022000 \| organism=Plasmodium_berghei_ANKA \| gene_product=40S ribosomal protein S25, putative \| transcript_product=40S ribosomal protein S25, putative \| location=PbANKA_10_v3: 891797-892591(+) \| protein_length=105 \| | 2 | 1 | 0 | 0 | 0 | 0 | 0 | 0 | 0 | 2 | 1 | 1 | 0 | 0 | 0 | 1 | 0 | 0 | 0 | 0 | 1 | 0 | 0 | 0 | 0 | 0 |
| Peroxiredoxin-2 OS=Mus musculus OX=10090 GN=Prdx2 PE=1 SV=3 | 0 | 0 | 0 | 0 | 0 | 0 | 0 | 0 | 0 | 0 | 1 | 1 | 0 | 0 | 0 | 4 | 2 | 0 | 0 | 0 | 1 | 0 | 0 | 0 | 2 | 0 |
| Heparin cofactor 2 OS=Mus musculus OX=10090 GN=Serpind1 PE=1 SV=1 | 0 | 0 | 0 | 0 | 0 | 0 | 0 | 0 | 0 | 0 | 0 | 0 | 0 | 0 | 0 | 0 | 0 | 0 | 0 | 0 | 0 | 0 | 0 | 2 | 0 | 0 |
| Myosin light polypeptide 6 OS=Mus musculus OX=10090 GN=Myl6 PE=1 SV=3 | 0 | 0 | 0 | 0 | 0 | 0 | 0 | 0 | 0 | 1 | 0 | 5 | 0 | 0 | 0 | 0 | 5 | 0 | 0 | 0 | 0 | 0 | 1 | 2 | 0 | 0 |
| Tubulin alpha-1B chain OS=Mus musculus OX=10090 GN=Tuba1b PE=1 SV=2 | 0 | 0 | 0 | 0 | 0 | 0 | 0 | 0 | 0 | 0 | 0 | 4 | 0 | 0 | 0 | 1 | 2 | 0 | 0 | 0 | 2 | 0 | 0 | 2 | 0 | 0 |
| Antithrombin-III OS=Mus musculus OX=10090 GN=Serpinc1 PE=1 SV=1 | 0 | 0 | 0 | 1 | 1 | 0 | 0 | 0 | 0 | 0 | 0 | 2 | 0 | 0 | 0 | 0 | 0 | 0 | 0 | 0 | 0 | 0 | 0 | 1 | 0 | 0 |
| \| transcript=PBANKA_0619700.1 \| gene=PBANKA_0619700 \| organism=Plasmodium_berghei_ANKA \| gene_product=rhoptry-associated leucine zipper-like protein 1, putative \| transcript_product=rhoptry-associated leucine zipper-like protein 1, putative \| location=PbANKA | 0 | 0 | 0 | 0 | 0 | 0 | 0 | 0 | 0 | 0 | 0 | 6 | 2 | 0 | 0 | 0 | 0 | 0 | 0 | 0 | 0 | 0 | 0 | 1 | 0 | 0 |
| Histone H1.4 OS=Mus musculus OX=10090 GN=H1-4 PE=1 SV=2 | 0 | 0 | 0 | 0 | 0 | 7 | 0 | 5 | 0 | 0 | 0 | 0 | 0 | 0 | 0 | 0 | 0 | 0 | 0 | 0 | 0 | 0 | 1 | 0 | 0 | 0 |
| Vinculin OS=Mus musculus OX=10090 GN=Vcl PE=1 SV=4 | 0 | 0 | 0 | 0 | 0 | 0 | 0 | 0 | 0 | 0 | 0 | 0 | 0 | 0 | 0 | 0 | 10 | 0 | 0 | 0 | 0 | 0 | 0 | 0 | 0 | 0 |
| \| transcript=PBANKA_0914300.1 \| gene=PBANKA_0914300 \| organism=Plasmodium_berghei_ANKA \| gene_product=protein disulfide-isomerase, putative \| transcript_product=protein disulfide-isomerase, putative \| location=PbANKA_09_v3:549526-550797(+) \| protein_length=4 | 0 | 0 | 0 | 1 | 0 | 0 | 0 | 0 | 0 | 0 | 0 | 2 | 0 | 1 | 0 | 2 | 0 | 0 | 0 | 0 | 0 | 0 | 0 | 2 | 0 | 0 |
| \| transcript=PBANKA_0926700.1 \| gene=PBANKA_0926700 \| organism=Plasmodium_berghei_ANKA \| gene_product=exported protein 1 \| transcript_product=exported protein 1 \| location=PbANKA_09_v3:991568-992293(+) \| protein_length=166 \| sequence_SO=chromosome \| SO=prote | 0 | 0 | 0 | 0 | 0 | 0 | 0 | 0 | 0 | 3 | 0 | 1 | 3 | 0 | 0 | 0 | 0 | 0 | 0 | 0 | 0 | 0 | 1 | 2 | 0 | 0 |

<table>
<tr><th></th><th></th><th></th><th></th><th></th><th></th><th></th><th></th><th></th><th></th><th></th><th></th><th></th><th></th><th></th><th></th><th></th><th></th><th></th><th></th><th></th><th></th><th></th><th></th><th></th><th></th><th></th></tr>
<tr><td>| transcript=PBANKA_1106700.1 | gene=PBANKA_1106700 | organism=Plasmodium_berghei_ANKA | gene_product=60S ribosomal protein L4, putative | transcript_product=60S ribosomal protein L4, putative | location=PbANKA_11_v3: 264086-265321(-) | protein_length=411 | s</td><td>1</td><td>0</td><td>0</td><td>1</td><td>1</td><td>0</td><td>0</td><td>0</td><td>0</td><td>0</td><td>0</td><td>3</td><td>0</td><td>0</td><td>0</td><td>0</td><td>0</td><td>0</td><td>0</td><td>0</td><td>0</td><td>0</td><td>2</td><td>0</td><td>0</td><td>0</td></tr>
<tr><td>Ig kappa chain V-III region CBPC 101 OS=Mus musculus OX=10090 PE=1 SV=1</td><td>0</td><td>0</td><td>0</td><td>0</td><td>0</td><td>0</td><td>0</td><td>0</td><td>0</td><td>0</td><td>0</td><td>4</td><td>2</td><td>0</td><td>0</td><td>0</td><td>0</td><td>0</td><td>0</td><td>0</td><td>0</td><td>0</td><td>1</td><td>3</td><td>0</td><td>0</td></tr>
<tr><td>| transcript=PBANKA_1032100.1 | gene=PBANKA_1032100 | organism=Plasmodium_berghei_ANKA | gene_product=rhoptry-associated protein 1 | transcript_product=rhoptry-associated protein 1 | location=PbANKA_10_v3:1292452-1294266(-) | protein_length=604 | sequence_SO</td><td>0</td><td>0</td><td>0</td><td>0</td><td>0</td><td>0</td><td>0</td><td>0</td><td>0</td><td>0</td><td>0</td><td>6</td><td>2</td><td>0</td><td>0</td><td>0</td><td>0</td><td>0</td><td>0</td><td>0</td><td>0</td><td>0</td><td>0</td><td>0</td><td>0</td><td>0</td></tr>
<tr><td>Protein 4.1</td><td>0</td><td>0</td><td>0</td><td>0</td><td>0</td><td>0</td><td>0</td><td>1</td><td>0</td><td>0</td><td>0</td><td>0</td><td>0</td><td>0</td><td>0</td><td>0</td><td>6</td><td>0</td><td>0</td><td>0</td><td>0</td><td>0</td><td>0</td><td>0</td><td>0</td><td>0</td></tr>
<tr><td>| transcript=PBANKA_0702800.1 | gene=PBANKA_0702800 | organism=Plasmodium_berghei_ANKA | gene_product=protein disulfide-isomerase, putative | transcript_product=protein disulfide-isomerase, putative | location=PbANKA_07_v3:143845-145402(+) | protein_length=4</td><td>0</td><td>0</td><td>0</td><td>0</td><td>0</td><td>0</td><td>0</td><td>0</td><td>0</td><td>0</td><td>0</td><td>0</td><td>0</td><td>1</td><td>0</td><td>7</td><td>3</td><td>0</td><td>0</td><td>0</td><td>0</td><td>0</td><td>0</td><td>0</td><td>0</td><td>0</td></tr>
<tr><td>| transcript=PBANKA_1416300.1 | gene=PBANKA_1416300 | organism=Plasmodium_berghei_ANKA | gene_product=40S ribosomal protein S19, putative | transcript_product=40S ribosomal protein S19, putative | location=PbANKA_14_v3: 653704-654794(+) | protein_length=145 |</td><td>1</td><td>0</td><td>1</td><td>0</td><td>1</td><td>0</td><td>0</td><td>0</td><td>0</td><td>2</td><td>0</td><td>0</td><td>0</td><td>0</td><td>0</td><td>0</td><td>0</td><td>0</td><td>0</td><td>0</td><td>1</td><td>0</td><td>0</td><td>0</td><td>1</td><td>0</td></tr>
<tr><td>Keratin, type II cuticular Hb2 OS=Mus musculus OX=10090 GN=Krt82 PE=1 SV=2</td><td>1</td><td>0</td><td>2</td><td>1</td><td>0</td><td>1</td><td>0</td><td>0</td><td>0</td><td>0</td><td>0</td><td>1</td><td>0</td><td>1</td><td>1</td><td>0</td><td>2</td><td>0</td><td>1</td><td>0</td><td>0</td><td>0</td><td>0</td><td>0</td><td>0</td><td>0</td></tr>
<tr><td>| transcript=PBANKA_0712600.1 | gene=PBANKA_0712600 | organism=Plasmodium_berghei_ANKA | gene_product=14-3-3 protein | transcript_product=14-3-3 protein | location=PbANKA_07_v3:480414-481905(-) | protein_length=262 | sequence_SO=chromosome | SO=protein_codin</td><td>0</td><td>0</td><td>0</td><td>0</td><td>0</td><td>0</td><td>0</td><td>0</td><td>0</td><td>1</td><td>0</td><td>4</td><td>0</td><td>0</td><td>0</td><td>0</td><td>0</td><td>0</td><td>0</td><td>0</td><td>0</td><td>0</td><td>1</td><td>0</td><td>0</td><td>0</td></tr>
<tr><td>Keratin, type I cuticular Ha6 OS=Homo sapiens GN=KRT36 PE=1 SV=1</td><td>0</td><td>0</td><td>0</td><td>0</td><td>0</td><td>0</td><td>0</td><td>0</td><td>0</td><td>0</td><td>0</td><td>0</td><td>0</td><td>0</td><td>0</td><td>0</td><td>0</td><td>0</td><td>0</td><td>0</td><td>10</td><td>0</td><td>0</td><td>0</td><td>0</td><td>0</td></tr>
<tr><td>| transcript=PBANKA_1204400.1 | gene=PBANKA_1204400 | organism=Plasmodium_berghei_ANKA | gene_product=DNA/RNA-binding protein Alba 3, putative | transcript_product=DNA/RNA-binding protein Alba 3, putative | location=PbANKA_12_v3:192634-193274(-) | protein_le</td><td>0</td><td>0</td><td>0</td><td>0</td><td>0</td><td>0</td><td>0</td><td>0</td><td>0</td><td>1</td><td>0</td><td>2</td><td>0</td><td>0</td><td>0</td><td>1</td><td>0</td><td>0</td><td>0</td><td>0</td><td>0</td><td>0</td><td>1</td><td>2</td><td>1</td><td>1</td></tr>
<tr><td>| transcript=PBANKA_1101300.1 | gene=PBANKA_1101300 | organism=Plasmodium_berghei_ANKA | gene_product=skeleton-binding protein 1 | transcript_product=skeleton-binding protein 1 | location=PbANKA_11_v3:83394-85785(+) | protein_length=762 | sequence_SO=chromos</td><td>0</td><td>0</td><td>0</td><td>0</td><td>0</td><td>0</td><td>0</td><td>0</td><td>0</td><td>1</td><td>0</td><td>2</td><td>0</td><td>0</td><td>0</td><td>0</td><td>0</td><td>0</td><td>0</td><td>0</td><td>1</td><td>0</td><td>1</td><td>1</td><td>0</td><td>0</td></tr>
<tr><td>L-lactate dehydrogenase C chain OS=Mus musculus OX=10090 GN=Ldhc PE=1 SV=2</td><td>0</td><td>0</td><td>0</td><td>1</td><td>0</td><td>0</td><td>0</td><td>0</td><td>0</td><td>0</td><td>0</td><td>0</td><td>0</td><td>0</td><td>0</td><td>1</td><td>1</td><td>0</td><td>0</td><td>0</td><td>2</td><td>0</td><td>0</td><td>0</td><td>0</td><td>0</td></tr>
<tr><td>| transcript=PBANKA_0938300.1 | gene=PBANKA_0938300 | organism=Plasmodium_berghei_ANKA | gene_product=heat shock protein J2, putative | transcript_product=heat shock protein J2, putative | location=PbANKA_09_v3:1375279-1376940(+) | protein_length=553 | seque</td><td>0</td><td>0</td><td>0</td><td>0</td><td>0</td><td>0</td><td>0</td><td>0</td><td>0</td><td>4</td><td>0</td><td>1</td><td>0</td><td>0</td><td>0</td><td>2</td><td>0</td><td>0</td><td>0</td><td>0</td><td>0</td><td>0</td><td>0</td><td>1</td><td>0</td><td>0</td></tr>
<tr><td>Tubulin alpha-4A chain OS=Mus musculus OX=10090 GN=Tuba4a PE=1 SV=1</td><td>0</td><td>0</td><td>0</td><td>2</td><td>0</td><td>0</td><td>0</td><td>0</td><td>0</td><td>0</td><td>0</td><td>0</td><td>0</td><td>0</td><td>0</td><td>1</td><td>5</td><td>0</td><td>0</td><td>0</td><td>0</td><td>0</td><td>0</td><td>1</td><td>0</td><td>0</td></tr>
<tr><td>| transcript=PBANKA_0915000.1 | gene=PBANKA_0915000 | organism=Plasmodium_berghei_ANKA | gene_product=apical membrane antigen 1 | transcript_product=apical membrane antigen 1 | location=PbANKA_09_v3:573540-575210(-) | protein_length=556 | sequence_SO=chromos</td><td>0</td><td>0</td><td>0</td><td>0</td><td>0</td><td>0</td><td>0</td><td>0</td><td>0</td><td>0</td><td>0</td><td>3</td><td>2</td><td>0</td><td>0</td><td>0</td><td>0</td><td>0</td><td>0</td><td>0</td><td>0</td><td>0</td><td>1</td><td>0</td><td>0</td><td>0</td></tr>
<tr><td>| transcript=PBANKA_1146000.1 | gene=PBANKA_1146000 | organism=Plasmodium_berghei_ANKA | gene_product=tryptophan-rich protein | transcript_product=tryptophan-rich protein | location=PbANKA_11_v3:1719210-1721298(-) | protein_length=642 | sequence_SO=chromosom</td><td>0</td><td>0</td><td>0</td><td>0</td><td>0</td><td>0</td><td>0</td><td>0</td><td>0</td><td>0</td><td>0</td><td>3</td><td>0</td><td>0</td><td>0</td><td>0</td><td>0</td><td>0</td><td>0</td><td>0</td><td>0</td><td>0</td><td>0</td><td>1</td><td>0</td><td>0</td></tr>
<tr><td>Ig kappa chain V-III region PC 7175 OS=Mus musculus OX=10090 PE=1 SV=1</td><td>0</td><td>0</td><td>0</td><td>0</td><td>0</td><td>0</td><td>0</td><td>0</td><td>0</td><td>0</td><td>0</td><td>4</td><td>2</td><td>0</td><td>0</td><td>0</td><td>0</td><td>0</td><td>0</td><td>0</td><td>0</td><td>0</td><td>0</td><td>3</td><td>0</td><td>0</td></tr>
<tr><td>Heat shock cognate 71 kDa protein OS=Mus musculus OX=10090 GN=Hspa8 PE=1 SV=1</td><td>0</td><td>0</td><td>0</td><td>0</td><td>0</td><td>0</td><td>0</td><td>0</td><td>0</td><td>0</td><td>0</td><td>0</td><td>0</td><td>0</td><td>0</td><td>5</td><td>5</td><td>0</td><td>0</td><td>0</td><td>0</td><td>0</td><td>0</td><td>0</td><td>0</td><td>0</td></tr>
<tr><td>Prelamin-A/C OS=Mus musculus OX=10090 GN=Lmna PE=1 SV=2</td><td>0</td><td>0</td><td>0</td><td>0</td><td>0</td><td>0</td><td>0</td><td>0</td><td>0</td><td>0</td><td>0</td><td>0</td><td>0</td><td>0</td><td>0</td><td>0</td><td>0</td><td>0</td><td>0</td><td>0</td><td>7</td><td>0</td><td>0</td><td>0</td><td>0</td><td>0</td></tr>
<tr><td>Ig kappa chain V-V region MOPC 149 OS=Mus musculus OX=10090 PE=1 SV=1</td><td>0</td><td>0</td><td>0</td><td>0</td><td>0</td><td>0</td><td>0</td><td>0</td><td>0</td><td>1</td><td>1</td><td>2</td><td>1</td><td>0</td><td>0</td><td>0</td><td>0</td><td>0</td><td>0</td><td>0</td><td>0</td><td>0</td><td>1</td><td>2</td><td>0</td><td>0</td></tr>
<tr><td>PBANKA_0925400.1-p1-DECOY</td><td>0</td><td>0</td><td>0</td><td>1</td><td>0</td><td>0</td><td>0</td><td>0</td><td>0</td><td>0</td><td>0</td><td>2</td><td>0</td><td>0</td><td>0</td><td>0</td><td>0</td><td>0</td><td>0</td><td>0</td><td>0</td><td>0</td><td>0</td><td>1</td><td>0</td><td>0</td></tr>
<tr><td>Immunoglobulin kappa chain variable 9-120 OS=Mus musculus OX=10090 GN=Igkv9-120 PE=1 SV=1</td><td>0</td><td>0</td><td>0</td><td>0</td><td>0</td><td>0</td><td>0</td><td>0</td><td>0</td><td>1</td><td>1</td><td>3</td><td>1</td><td>0</td><td>0</td><td>0</td><td>0</td><td>0</td><td>0</td><td>0</td><td>0</td><td>0</td><td>1</td><td>2</td><td>0</td><td>0</td></tr>
<tr><td>| transcript=PBANKA_0717800.1 | gene=PBANKA_0717800 | organism=Plasmodium_berghei_ANKA | gene_product=60S ribosomal protein L15, putative | transcript_product=60S ribosomal protein L15, putative | location=PbANKA_07_v3: 641432-642319(-) | protein_length=205 |</td><td>1</td><td>1</td><td>0</td><td>1</td><td>3</td><td>0</td><td>0</td><td>0</td><td>0</td><td>0</td><td>0</td><td>0</td><td>0</td><td>0</td><td>0</td><td>0</td><td>0</td><td>1</td><td>0</td><td>0</td><td>1</td><td>0</td><td>0</td><td>0</td><td>0</td><td>0</td></tr>
</table>

| | | | | | | | | | | | | | | | | | | | | | | | | | | |
|---|---|---|---|---|---|---|---|---|---|---|---|---|---|---|---|---|---|---|---|---|---|---|---|---|---|---|
| \| transcript=PBANKA_1331900.1 \| gene=PBANKA_1331900 \| organism=Plasmodium_berghei_ANKA \| gene_product=eukaryotic initiation factor 4a, putative \| transcript_product=eukaryotic initiation factor 4a, putative \| location=PbANKA_13_v3:1297871-1299452(+) \| protei | 0 | 0 | 0 | 0 | 0 | 0 | 0 | 0 | 0 | 0 | 0 | 3 | 0 | 0 | 0 | 0 | 1 | 0 | 0 | 0 | 0 | 0 | 1 | 1 | 0 | 0 |
| \| transcript=PBANKA_1355100.1 \| gene=PBANKA_1355100 \| organism=Plasmodium_berghei_ANKA \| gene_product=40S ribosomal protein S6, putative \| transcript_product=40S ribosomal protein S6, putative \| location=PbANKA_13_v3:2075849-2076781(-) \| protein_length=310 \| | 1 | 0 | 0 | 0 | 1 | 0 | 0 | 0 | 0 | 3 | 1 | 0 | 0 | 0 | 0 | 0 | 0 | 0 | 0 | 0 | 0 | 0 | 0 | 0 | 0 | 0 |
| Histone H2A type 2-C OS=Mus musculus OX=10090 GN=H2ac20 PE=1 SV=3 | 0 | 0 | 0 | 0 | 0 | 0 | 0 | 0 | 0 | 0 | 0 | 4 | 1 | 0 | 0 | 0 | 1 | 0 | 0 | 0 | 0 | 0 | 0 | 0 | 0 | 0 |
| Myosin-3 OS=Mus musculus OX=10090 GN=Myh3 PE=2 SV=2 | 0 | 0 | 0 | 0 | 0 | 0 | 0 | 0 | 0 | 0 | 0 | 0 | 0 | 0 | 6 | 0 | 0 | 0 | 0 | 0 | 0 | 0 | 0 | 0 | 0 | 0 |
| \| transcript=PBANKA_0520200.1 \| gene=PBANKA_0520200 \| organism=Plasmodium_berghei_ANKA \| gene_product=ADP, ATP carrier protein 1, putative \| transcript_product=ADP,ATP carrier protein 1, putative \| location=PbANKA_05_v3:731281-732186(-) \| protein_length=301 \| | 2 | 0 | 0 | 0 | 0 | 0 | 0 | 0 | 0 | 0 | 0 | 0 | 1 | 0 | 0 | 0 | 0 | 0 | 0 | 0 | 0 | 0 | 0 | 0 | 0 | 0 |
| High affinity immunoglobulin gamma Fc receptor I OS=Mus musculus OX=10090 GN=Fcgr1 PE=1 SV=1 | 0 | 0 | 0 | 0 | 0 | 0 | 2 | 1 | 0 | 0 | 0 | 0 | 0 | 0 | 0 | 1 | 0 | 0 | 0 | 0 | 0 | 0 | 0 | 0 | 0 | 0 |
| Pyruvate kinase PKM OS=Mus musculus OX=10090 GN=Pkm PE=1 SV=4 | 0 | 0 | 0 | 1 | 0 | 0 | 0 | 0 | 0 | 0 | 0 | 0 | 0 | 0 | 2 | 0 | 1 | 0 | 0 | 0 | 1 | 0 | 0 | 0 | 0 | 0 |
| \| transcript=PBANKA_1206900.1 \| gene=PBANKA_1206900 \| organism=Plasmodium_berghei_ANKA \| gene_product=tubulin beta chain, putative \| transcript_product=tubulin beta chain, putative \| location=PbANKA_12_v3:282779-284936(-) \| protein_length=445 \| sequence_SO=c | 0 | 0 | 0 | 0 | 0 | 0 | 0 | 0 | 0 | 0 | 0 | 3 | 1 | 0 | 0 | 0 | 0 | 0 | 0 | 0 | 0 | 0 | 2 | 1 | 0 | 0 |
| Stomatin OS=Mus musculus OX=10090 GN=Stom PE=1 SV=3 | 2 | 0 | 0 | 0 | 0 | 0 | 0 | 0 | 0 | 0 | 0 | 0 | 0 | 0 | 0 | 2 | 3 | 0 | 0 | 0 | 0 | 0 | 0 | 0 | 0 | 0 |
| Ig heavy chain V region MOPC 141 OS=Mus musculus OX=10090 PE=4 SV=1 | 0 | 0 | 0 | 0 | 0 | 0 | 0 | 0 | 0 | 0 | 0 | 4 | 1 | 0 | 0 | 0 | 0 | 0 | 0 | 0 | 0 | 0 | 0 | 2 | 0 | 0 |
| Apolipoprotein B-100 OS=Mus musculus OX=10090 GN=Apob PE=1 SV=1 | 0 | 0 | 0 | 0 | 0 | 0 | 0 | 0 | 0 | 0 | 0 | 0 | 0 | 0 | 0 | 0 | 0 | 0 | 2 | 2 | 0 | 0 | 0 | 2 | 0 | 0 |
| Rab GTPase-binding effector protein 2 OS=Mus musculus OX=10090 GN=Rabep2 PE=1 SV=3 | 0 | 0 | 0 | 0 | 0 | 0 | 0 | 0 | 0 | 0 | 0 | 0 | 0 | 1 | 0 | 5 | 2 | 0 | 0 | 0 | 0 | 0 | 0 | 0 | 0 | 0 |
| Ig kappa chain V-III region PC 2880/PC 1229 OS=Mus musculus OX=10090 PE=1 SV=1 | 0 | 0 | 0 | 0 | 0 | 0 | 0 | 0 | 0 | 0 | 0 | 3 | 0 | 0 | 0 | 0 | 0 | 0 | 0 | 0 | 0 | 0 | 0 | 2 | 0 | 0 |
| Annexin A2 OS=Mus musculus OX=10090 GN=Anxa2 PE=1 SV=2 | 0 | 0 | 0 | 0 | 0 | 0 | 0 | 0 | 0 | 0 | 0 | 0 | 0 | 0 | 0 | 0 | 0 | 0 | 0 | 0 | 5 | 0 | 0 | 0 | 4 | 0 |
| Protein-glutamine gamma-glutamyltransferase K OS=Mus musculus OX=10090 GN=Tgm1 PE=1 SV=2 | 0 | 0 | 0 | 0 | 0 | 0 | 0 | 0 | 0 | 0 | 0 | 0 | 0 | 0 | 0 | 0 | 0 | 0 | 0 | 0 | 3 | 0 | 0 | 0 | 1 | 0 |
| \| transcript=PBANKA_0623100.1 \| gene=PBANKA_0623100 \| organism=Plasmodium_berghei_ANKA \| gene_product=tryptophan-rich protein IPIS3 \| transcript_product=tryptophan-rich protein IPIS3 \| location=PbANKA_06_v3:940354-942905(-) \| protein_length=793 \| sequence_SO | 0 | 0 | 0 | 0 | 0 | 0 | 0 | 0 | 0 | 0 | 0 | 5 | 0 | 0 | 0 | 0 | 0 | 0 | 0 | 0 | 0 | 0 | 0 | 2 | 0 | 0 |
| \| transcript=PBANKA_0524300.1 \| gene=PBANKA_0524300 \| organism=Plasmodium_berghei_ANKA \| gene_product=tryptophan-rich protein IPIS2 \| transcript_product=tryptophan-rich protein IPIS2 \| location=PbANKA_05_v3:894066-895250(-) \| protein_length=354 \| sequence_SO | 0 | 0 | 0 | 0 | 0 | 0 | 0 | 0 | 0 | 0 | 0 | 4 | 0 | 0 | 0 | 0 | 0 | 0 | 0 | 0 | 0 | 0 | 0 | 2 | 0 | 0 |
| Keratin 83 OS=Mus musculus OX=10090 GN=Krt83 PE=1 SV=1 | 0 | 0 | 0 | 0 | 0 | 0 | 0 | 0 | 0 | 0 | 0 | 0 | 0 | 0 | 0 | 1 | 0 | 0 | 0 | 0 | 8 | 0 | 0 | 0 | 0 | 0 |
| Myosin-7 OS=Mus musculus OX=10090 GN=Myh7 PE=2 SV=1 | 0 | 0 | 0 | 0 | 0 | 0 | 0 | 0 | 0 | 0 | 0 | 0 | 0 | 0 | 6 | 0 | 0 | 0 | 0 | 0 | 0 | 0 | 0 | 0 | 0 | 0 |
| Ig kappa chain V-II region 7S34.1 OS=Mus musculus OX=10090 PE=1 SV=1 | 0 | 0 | 0 | 0 | 0 | 0 | 0 | 0 | 0 | 1 | 1 | 3 | 1 | 0 | 0 | 0 | 0 | 0 | 0 | 0 | 0 | 0 | 0 | 2 | 0 | 0 |
| \| transcript=PBANKA_1400700.1 \| gene=PBANKA_1400700 \| organism=Plasmodium_berghei_ANKA \| gene_product=Plasmodium exported protein, unknown function \| transcript_product=Plasmodium exported protein, unknown function \| location=PbANKA_14_v3:83081-83763(-) \| pr | 0 | 0 | 0 | 0 | 0 | 0 | 0 | 0 | 0 | 0 | 0 | 2 | 1 | 0 | 0 | 0 | 0 | 0 | 0 | 0 | 0 | 0 | 1 | 0 | 0 | 0 |
| \| transcript=PBANKA_1234800.1 \| gene=PBANKA_1234800 \| organism=Plasmodium_berghei_ANKA \| gene_product=40S ribosomal protein S9, putative \| transcript_product=40S ribosomal protein S9, putative \| location=PbANKA_12_v3:1336256-1337356(+) \| protein_length=189 \| | 2 | 0 | 0 | 0 | 1 | 0 | 0 | 0 | 0 | 0 | 0 | 0 | 0 | 0 | 0 | 1 | 0 | 0 | 0 | 0 | 0 | 0 | 0 | 0 | 1 | 0 |
| \| transcript=PBANKA_0407700.1 \| gene=PBANKA_0407700 \| organism=Plasmodium_berghei_ANKA \| gene_product=60S acidic ribosomal protein P2, putative \| transcript_product=60S acidic ribosomal protein P2, putative \| location=PbANKA_04_v3:282316-282651(+) \| protein_ | 0 | 0 | 0 | 0 | 0 | 0 | 0 | 0 | 0 | 1 | 0 | 3 | 0 | 0 | 0 | 0 | 0 | 0 | 0 | 0 | 0 | 0 | 0 | 1 | 0 | 0 |
| \| transcript=PBANKA_1238800.1 \| gene=PBANKA_1238800 \| organism=Plasmodium_berghei_ANKA \| gene_product=karyopherin beta, putative \| transcript_product=karyopherin beta, putative \| location=PbANKA_12_v3:1489931-1493311(+) \| protein_length=1126 \| sequence_SO=ch | 0 | 0 | 0 | 0 | 0 | 0 | 0 | 0 | 0 | 0 | 0 | 3 | 1 | 0 | 0 | 0 | 0 | 0 | 0 | 0 | 0 | 0 | 1 | 1 | 0 | 0 |
| Fibronectin OS=Mus musculus OX=10090 GN=Fn1 PE=1 SV=4 | 0 | 0 | 0 | 0 | 0 | 1 | 0 | 0 | 0 | 0 | 0 | 2 | 0 | 0 | 0 | 0 | 0 | 0 | 0 | 0 | 0 | 0 | 0 | 1 | 0 | 0 |

<table>
<tr><td>| transcript=PBANKA_1423300.1 | gene=PBANKA_1423300 | organism=Plasmodium_berghei_ANKA | gene_product=DNA/RNA-binding protein Alba 1, putative | transcript_product=DNA/RNA-binding protein Alba 1, putative | location=PbANKA_14_v3:905727-906455(-) | protein_le</td><td>0</td><td>0</td><td>0</td><td>0</td><td>1</td><td>0</td><td>0</td><td>0</td><td>0</td><td>0</td><td>0</td><td>0</td><td>0</td><td>0</td><td>0</td><td>3</td><td>1</td><td>0</td><td>0</td><td>0</td><td>0</td><td>0</td><td>0</td><td>0</td><td>0</td><td>0</td></tr>
<tr><td>Tubulin beta-4B chain OS=Mus musculus OX=10090 GN=Tubb4b PE=1 SV=1</td><td>0</td><td>0</td><td>0</td><td>0</td><td>0</td><td>0</td><td>0</td><td>0</td><td>0</td><td>0</td><td>0</td><td>4</td><td>1</td><td>0</td><td>0</td><td>0</td><td>1</td><td>0</td><td>0</td><td>0</td><td>0</td><td>0</td><td>0</td><td>0</td><td>0</td><td>0</td></tr>
<tr><td>Profilin-1 OS=Mus musculus OX=10090 GN=Pfn1 PE=1 SV=2</td><td>0</td><td>0</td><td>0</td><td>1</td><td>0</td><td>0</td><td>0</td><td>0</td><td>0</td><td>0</td><td>0</td><td>0</td><td>0</td><td>0</td><td>0</td><td>0</td><td>1</td><td>0</td><td>0</td><td>0</td><td>0</td><td>0</td><td>0</td><td>2</td><td>0</td><td>0</td></tr>
<tr><td>| transcript=PBANKA_1334300.1 | gene=PBANKA_1334300 | organism=Plasmodium_berghei_ANKA | gene_product=exported protein 2 | transcript_product=exported protein 2 | location=PbANKA_13_v3:1374381-1375409(-) | protein_length=269 | sequence_SO=chromosome | SO=pro</td><td>0</td><td>0</td><td>0</td><td>0</td><td>0</td><td>0</td><td>0</td><td>0</td><td>0</td><td>0</td><td>0</td><td>2</td><td>2</td><td>0</td><td>0</td><td>0</td><td>0</td><td>0</td><td>0</td><td>0</td><td>0</td><td>0</td><td>0</td><td>1</td><td>0</td><td>0</td></tr>
<tr><td>| transcript=PBANKA_1215600.1 | gene=PBANKA_1215600 | organism=Plasmodium_berghei_ANKA | gene_product=40S ribosomal protein S3A, putative | transcript_product=40S ribosomal protein S3A, putative | location=PbANKA_12_v3:586716-587998(+) | protein_length=262 |</td><td>0</td><td>0</td><td>0</td><td>0</td><td>0</td><td>0</td><td>0</td><td>0</td><td>0</td><td>1</td><td>0</td><td>2</td><td>0</td><td>0</td><td>0</td><td>0</td><td>0</td><td>0</td><td>0</td><td>0</td><td>0</td><td>0</td><td>1</td><td>0</td><td>0</td><td>0</td></tr>
<tr><td>14-3-3 protein theta OS=Mus musculus OX=10090 GN=Ywhaq PE=1 SV=1</td><td>0</td><td>0</td><td>0</td><td>0</td><td>0</td><td>0</td><td>0</td><td>0</td><td>0</td><td>0</td><td>0</td><td>0</td><td>0</td><td>0</td><td>0</td><td>0</td><td>0</td><td>0</td><td>0</td><td>0</td><td>2</td><td>0</td><td>0</td><td>2</td><td>0</td><td>0</td></tr>
<tr><td>| transcript=PBANKA_1456100.1 | gene=PBANKA_1456100 | organism=Plasmodium_berghei_ANKA | gene_product=40S ribosomal protein S17, putative | transcript_product=40S ribosomal protein S17, putative | location=PbANKA_14_v3:2163552-2164337(+) | protein_length=137</td><td>1</td><td>0</td><td>0</td><td>0</td><td>0</td><td>0</td><td>0</td><td>0</td><td>0</td><td>2</td><td>2</td><td>0</td><td>0</td><td>0</td><td>0</td><td>0</td><td>0</td><td>0</td><td>0</td><td>0</td><td>0</td><td>0</td><td>0</td><td>0</td><td>0</td><td>0</td></tr>
<tr><td>Immunoglobulin J chain OS=Mus musculus OX=10090 GN=Jchain PE=1 SV=4</td><td>0</td><td>0</td><td>0</td><td>0</td><td>0</td><td>0</td><td>0</td><td>0</td><td>0</td><td>0</td><td>0</td><td>2</td><td>1</td><td>0</td><td>0</td><td>0</td><td>0</td><td>0</td><td>0</td><td>0</td><td>0</td><td>0</td><td>0</td><td>2</td><td>0</td><td>0</td></tr>
<tr><td>Alpha-adducin OS=Mus musculus OX=10090 GN=Add1 PE=1 SV=2</td><td>0</td><td>0</td><td>0</td><td>0</td><td>0</td><td>0</td><td>0</td><td>0</td><td>0</td><td>0</td><td>0</td><td>4</td><td>0</td><td>0</td><td>0</td><td>0</td><td>0</td><td>0</td><td>0</td><td>0</td><td>0</td><td>0</td><td>0</td><td>0</td><td>0</td><td>0</td></tr>
<tr><td>| transcript=PBANKA_1116800.1 | gene=PBANKA_1116800 | organism=Plasmodium_berghei_ANKA | gene_product=BFR1 domain-containing protein, putative | transcript_product=BFR1 domain-containing protein, putative | location=PbANKA_11_v3:636950-638728(+) | protein_le</td><td>0</td><td>0</td><td>0</td><td>0</td><td>0</td><td>0</td><td>0</td><td>0</td><td>0</td><td>4</td><td>0</td><td>0</td><td>0</td><td>0</td><td>0</td><td>0</td><td>0</td><td>0</td><td>0</td><td>0</td><td>0</td><td>0</td><td>0</td><td>0</td><td>0</td><td>0</td></tr>
<tr><td>| transcript=PBANKA_1137800.1 | gene=PBANKA_1137800 | organism=Plasmodium_berghei_ANKA | gene_product=glideosome-associated connector | transcript_product=glideosome-associated connector | location=PbANKA_11_v3:1426696-1434816(+) | protein_length=2605 | sequ</td><td>0</td><td>0</td><td>0</td><td>0</td><td>0</td><td>0</td><td>0</td><td>0</td><td>0</td><td>0</td><td>0</td><td>1</td><td>0</td><td>0</td><td>0</td><td>0</td><td>0</td><td>0</td><td>0</td><td>0</td><td>0</td><td>0</td><td>3</td><td>0</td><td>0</td><td>0</td></tr>
<tr><td>Keratin, type II cytoskeletal 79 OS=Mus musculus OX=10090 GN=Krt79 PE=1 SV=2</td><td>0</td><td>0</td><td>0</td><td>0</td><td>0</td><td>0</td><td>0</td><td>0</td><td>0</td><td>0</td><td>0</td><td>0</td><td>0</td><td>0</td><td>0</td><td>0</td><td>0</td><td>0</td><td>0</td><td>0</td><td>0</td><td>0</td><td>0</td><td>0</td><td>7</td><td>0</td></tr>
<tr><td>Keratin, type II cytoskeletal 80 OS=Mus musculus OX=10090 GN=Krt80 PE=1 SV=1</td><td>0</td><td>0</td><td>0</td><td>0</td><td>0</td><td>0</td><td>0</td><td>0</td><td>0</td><td>0</td><td>2</td><td>0</td><td>1</td><td>0</td><td>0</td><td>0</td><td>0</td><td>0</td><td>0</td><td>0</td><td>0</td><td>0</td><td>0</td><td>0</td><td>1</td><td>0</td></tr>
<tr><td>| transcript=PBANKA_1125600.1 | gene=PBANKA_1125600 | organism=Plasmodium_berghei_ANKA | gene_product=pyruvate kinase, putative | transcript_product=pyruvate kinase, putative | location=PbANKA_11_v3:940783-943039(+) | protein_length=511 | sequence_SO=chromos</td><td>0</td><td>0</td><td>0</td><td>0</td><td>0</td><td>0</td><td>0</td><td>0</td><td>0</td><td>0</td><td>0</td><td>1</td><td>0</td><td>0</td><td>0</td><td>1</td><td>0</td><td>0</td><td>0</td><td>0</td><td>0</td><td>0</td><td>2</td><td>1</td><td>0</td><td>0</td></tr>
<tr><td>| transcript=PBANKA_0713100.1 | gene=PBANKA_0713100 | organism=Plasmodium_berghei_ANKA | gene_product=rhoptry neck protein 5 | transcript_product=rhoptry neck protein 5 | location=PbANKA_07_v3:496794-503841(+) | protein_length=1148 | sequence_SO=chromosome |</td><td>0</td><td>0</td><td>0</td><td>0</td><td>0</td><td>0</td><td>0</td><td>0</td><td>0</td><td>0</td><td>0</td><td>3</td><td>0</td><td>0</td><td>0</td><td>0</td><td>0</td><td>0</td><td>0</td><td>0</td><td>0</td><td>0</td><td>1</td><td>0</td><td>0</td><td>0</td></tr>
<tr><td>| transcript=PBANKA_0922100.1 | gene=PBANKA_0922100 | organism=Plasmodium_berghei_ANKA | gene_product=40S ribosomal protein S18, putative | transcript_product=40S ribosomal protein S18, putative | location=PbANKA_09_v3:808973-809654(+) | protein_length=156 |</td><td>1</td><td>0</td><td>0</td><td>0</td><td>0</td><td>0</td><td>0</td><td>0</td><td>0</td><td>2</td><td>0</td><td>0</td><td>0</td><td>0</td><td>0</td><td>0</td><td>0</td><td>0</td><td>0</td><td>0</td><td>0</td><td>0</td><td>0</td><td>0</td><td>0</td><td>0</td></tr>
<tr><td>Clathrin heavy chain 1 OS=Mus musculus OX=10090 GN=Cltc PE=1 SV=3</td><td>0</td><td>0</td><td>0</td><td>0</td><td>0</td><td>0</td><td>0</td><td>0</td><td>0</td><td>0</td><td>0</td><td>0</td><td>0</td><td>0</td><td>0</td><td>0</td><td>2</td><td>0</td><td>0</td><td>0</td><td>1</td><td>0</td><td>0</td><td>0</td><td>0</td><td>0</td></tr>
<tr><td>| transcript=PBANKA_0522800.1 | gene=PBANKA_0522800 | organism=Plasmodium_berghei_ANKA | gene_product=40S ribosomal protein S19, putative | transcript_product=40S ribosomal protein S19, putative | location=PbANKA_05_v3:851991-852951(+) | protein_length=172 |</td><td>0</td><td>0</td><td>0</td><td>0</td><td>0</td><td>0</td><td>0</td><td>0</td><td>0</td><td>1</td><td>0</td><td>0</td><td>0</td><td>0</td><td>0</td><td>0</td><td>0</td><td>0</td><td>0</td><td>0</td><td>0</td><td>0</td><td>2</td><td>0</td><td>0</td><td>0</td></tr>
<tr><td>Ig heavy chain V regions TEPC 15/S107/HPCM1/HPCM2/HPCM3 OS=Mus musculus OX=10090 PE=1 SV=1</td><td>0</td><td>0</td><td>0</td><td>0</td><td>0</td><td>0</td><td>0</td><td>0</td><td>0</td><td>2</td><td>0</td><td>3</td><td>0</td><td>0</td><td>0</td><td>0</td><td>0</td><td>0</td><td>0</td><td>0</td><td>0</td><td>0</td><td>0</td><td>0</td><td>0</td><td>0</td></tr>
<tr><td>Ig heavy chain V region 108A OS=Mus musculus OX=10090 GN=Igh-VJ558 PE=4 SV=1</td><td>0</td><td>0</td><td>0</td><td>0</td><td>0</td><td>0</td><td>0</td><td>0</td><td>0</td><td>0</td><td>0</td><td>3</td><td>0</td><td>0</td><td>0</td><td>0</td><td>0</td><td>0</td><td>0</td><td>0</td><td>0</td><td>0</td><td>0</td><td>3</td><td>0</td><td>0</td></tr>
<tr><td>E3 ubiquitin-protein ligase TRIM21 OS=Mus musculus OX=10090 GN=Trim21 PE=1 SV=1</td><td>0</td><td>0</td><td>0</td><td>0</td><td>0</td><td>0</td><td>0</td><td>0</td><td>0</td><td>3</td><td>2</td><td>0</td><td>0</td><td>0</td><td>0</td><td>0</td><td>0</td><td>0</td><td>0</td><td>0</td><td>0</td><td>0</td><td>0</td><td>0</td><td>0</td><td>0</td></tr>
<tr><td>Triosephosphate isomerase OS=Mus musculus OX=10090 GN=Tpi1 PE=1 SV=5</td><td>0</td><td>0</td><td>0</td><td>0</td><td>0</td><td>0</td><td>0</td><td>0</td><td>0</td><td>0</td><td>0</td><td>0</td><td>0</td><td>0</td><td>3</td><td>0</td><td>0</td><td>0</td><td>0</td><td>0</td><td>1</td><td>0</td><td>0</td><td>0</td><td>0</td><td>0</td></tr>
<tr><td>Myosin regulatory light chain 12B OS=Mus musculus OX=10090 GN=Myl12b PE=1 SV=2</td><td>0</td><td>0</td><td>0</td><td>0</td><td>0</td><td>0</td><td>0</td><td>0</td><td>0</td><td>0</td><td>0</td><td>4</td><td>0</td><td>0</td><td>0</td><td>0</td><td>1</td><td>0</td><td>0</td><td>0</td><td>0</td><td>0</td><td>0</td><td>0</td><td>0</td><td>0</td></tr>
<tr><td>Fructose-bisphosphate aldolase OS=Mus musculus OX=10090 GN=Aldoart1 PE=1 SV=1</td><td>0</td><td>0</td><td>0</td><td>0</td><td>0</td><td>0</td><td>0</td><td>0</td><td>0</td><td>0</td><td>0</td><td>0</td><td>0</td><td>0</td><td>0</td><td>0</td><td>1</td><td>0</td><td>0</td><td>0</td><td>2</td><td>0</td><td>0</td><td>0</td><td>0</td><td>0</td></tr>
<tr><td>Histone H1.5 OS=Mus musculus OX=10090 GN=H1-5 PE=1 SV=2</td><td>0</td><td>0</td><td>0</td><td>0</td><td>0</td><td>4</td><td>0</td><td>1</td><td>0</td><td>0</td><td>0</td><td>0</td><td>0</td><td>0</td><td>0</td><td>0</td><td>0</td><td>0</td><td>0</td><td>0</td><td>0</td><td>0</td><td>0</td><td>0</td><td>0</td><td>0</td></tr>
</table>

| | | | | | | | | | | | | | | | | | | | | | | | | | | |
|---|---|---|---|---|---|---|---|---|---|---|---|---|---|---|---|---|---|---|---|---|---|---|---|---|---|---|
| Tropomodulin-1 OS=Mus musculus OX=10090 GN=Tmod1 PE=1 SV=2 | 0 | 0 | 0 | 0 | 0 | 0 | 0 | 0 | 0 | 0 | 0 | 4 | 0 | 0 | 0 | 0 | 1 | 0 | 0 | 0 | 0 | 0 | 0 | 0 | 0 | 0 |
| \| transcript=PBANKA_1426900.1 \| gene=PBANKA_1426900 \| organism=Plasmodium_berghei_ANKA \| gene_product=ATP-dependent RNA helicase DBP1, putative \| transcript_product=ATP-dependent RNA helicase DBP1, putative \| location=PbANKA_14_v3:1035477-1038206(-) \| protei | 0 | 0 | 0 | 0 | 0 | 0 | 0 | 0 | 0 | 5 | 0 | 0 | 0 | 0 | 0 | 0 | 0 | 0 | 0 | 0 | 0 | 0 | 0 | 0 | 0 | 0 |
| Fermitin family homolog 3 OS=Mus musculus OX=10090 GN=Fermt3 PE=1 SV=1 | 0 | 0 | 0 | 0 | 0 | 0 | 0 | 0 | 0 | 0 | 0 | 0 | 0 | 0 | 0 | 0 | 4 | 0 | 0 | 0 | 0 | 0 | 0 | 0 | 0 | 0 |
| Delta-aminolevulinic acid dehydratase OS=Mus musculus OX=10090 GN=Alad PE=1 SV=1 | 0 | 0 | 0 | 0 | 0 | 0 | 0 | 0 | 0 | 0 | 0 | 0 | 0 | 0 | 0 | 4 | 0 | 0 | 0 | 0 | 0 | 0 | 0 | 0 | 0 | 0 |
| Alpha-actinin-4 OS=Mus musculus OX=10090 GN=Actn4 PE=1 SV=1 | 0 | 0 | 0 | 0 | 0 | 0 | 0 | 0 | 0 | 0 | 0 | 0 | 0 | 0 | 1 | 0 | 1 | 0 | 0 | 0 | 2 | 0 | 0 | 0 | 0 | 0 |
| Peroxiredoxin-4 OS=Mus musculus OX=10090 GN=Prdx4 PE=1 SV=1 | 0 | 0 | 0 | 0 | 0 | 0 | 0 | 0 | 0 | 0 | 0 | 1 | 0 | 0 | 0 | 0 | 0 | 0 | 0 | 0 | 2 | 0 | 0 | 0 | 0 | 0 |
| Ig heavy chain V region BCL1 OS=Mus musculus OX=10090 PE=4 SV=1 | 0 | 0 | 0 | 0 | 0 | 0 | 0 | 0 | 0 | 0 | 0 | 3 | 0 | 0 | 0 | 0 | 0 | 0 | 0 | 0 | 0 | 0 | 0 | 1 | 0 | 0 |
| SEC23-interacting protein OS=Mus musculus OX=10090 GN=Sec23ip PE=1 SV=2 | 0 | 0 | 0 | 0 | 0 | 0 | 0 | 0 | 0 | 0 | 0 | 0 | 0 | 0 | 0 | 0 | 3 | 0 | 0 | 0 | 0 | 0 | 0 | 0 | 0 | 0 |
| Ig lambda-3 chain C region OS=Mus musculus OX=10090 GN=Iglc3 PE=1 SV=1 | 0 | 0 | 0 | 0 | 0 | 0 | 0 | 0 | 0 | 0 | 0 | 3 | 0 | 0 | 0 | 0 | 0 | 0 | 0 | 0 | 0 | 0 | 0 | 2 | 0 | 0 |
| \| transcript=PBANKA_1357200.1 \| gene=PBANKA_1357200 \| organism=Plasmodium_berghei_ANKA \| gene_product=endoplasmic reticulum chaperone GRP170, putative \| transcript_product=endoplasmic reticulum chaperone GRP170, putative \| location=PbANKA_13_v3: 2179879-21826 | 0 | 0 | 0 | 0 | 0 | 0 | 0 | 0 | 0 | 0 | 0 | 0 | 0 | 0 | 0 | 2 | 1 | 0 | 0 | 0 | 0 | 0 | 0 | 0 | 0 | 0 |
| \| transcript=PBANKA_0311800.1 \| gene=PBANKA_0311800 \| organism=Plasmodium_berghei_ANKA \| gene_product=acyl-CoA synthetase, putative \| transcript_product=acyl-CoA synthetase, putative \| location=PbANKA_03_v3:411074-413335(-) \| protein_length=753 \| sequence_SO | 0 | 0 | 0 | 0 | 0 | 0 | 0 | 0 | 0 | 0 | 0 | 2 | 0 | 0 | 0 | 0 | 0 | 0 | 0 | 0 | 0 | 0 | 0 | 0 | 0 | 0 |
| \| transcript=PBANKA_0403800.1 \| gene=PBANKA_0403800 \| organism=Plasmodium_berghei_ANKA \| gene_product=transporter, putative \| transcript_product=transporter, putative \| location=PbANKA_04_v3:144333-146745(-) \| protein_length=735 \| sequence_SO=chromosome \| SO | 0 | 0 | 0 | 0 | 0 | 0 | 0 | 0 | 0 | 0 | 0 | 2 | 0 | 0 | 0 | 0 | 0 | 0 | 0 | 0 | 0 | 0 | 0 | 1 | 0 | 0 |
| \| transcript=PBANKA_1315700.1 \| gene=PBANKA_1315700 \| organism=Plasmodium_berghei_ANKA \| gene_product=rhoptry neck protein 2 \| transcript_product=rhoptry neck protein 2 \| location=PbANKA_13_v3:664125-670370(+) \| protein_length=2081 \| sequence_SO=chromosome \| | 0 | 0 | 0 | 0 | 0 | 0 | 0 | 0 | 0 | 0 | 0 | 4 | 0 | 0 | 0 | 0 | 0 | 0 | 0 | 0 | 0 | 0 | 0 | 0 | 0 | 0 |
| \| transcript=PBANKA_1103100.1 \| gene=PBANKA_1103100 \| organism=Plasmodium_berghei_ANKA \| gene_product=actin-depolymerizing factor 1 \| transcript_product=actin-depolymerizing factor 1 \| location=PbANKA_11_v3:137069-137623(+) \| protein_length=122 \| sequence_SO | 0 | 0 | 0 | 0 | 0 | 0 | 0 | 0 | 0 | 0 | 0 | 2 | 0 | 0 | 0 | 0 | 0 | 0 | 0 | 0 | 0 | 0 | 1 | 1 | 0 | 0 |
| \| transcript=PBANKA_0617100.1 \| gene=PBANKA_0617100 \| organism=Plasmodium_berghei_ANKA \| gene_product=60S ribosomal protein L11a, putative \| transcript_product=60S ribosomal protein L11a, putative \| location=PbANKA_06_v3: 726658-727475(-) \| protein_length=173 | 0 | 0 | 0 | 2 | 0 | 0 | 0 | 0 | 0 | 0 | 0 | 0 | 0 | 0 | 0 | 0 | 0 | 0 | 0 | 0 | 0 | 0 | 0 | 0 | 0 | 0 |
| Antileukoproteinase OS=Mus musculus OX=10090 GN=Slpi PE=1 SV=1 | 0 | 0 | 0 | 0 | 0 | 0 | 0 | 0 | 0 | 0 | 0 | 0 | 0 | 2 | 0 | 1 | 0 | 0 | 0 | 0 | 0 | 0 | 0 | 0 | 0 | 0 |
| Alpha-mannosidase 2C1 OS=Mus musculus OX=10090 GN=Man2c1 PE=1 SV=1 | 0 | 0 | 0 | 0 | 0 | 0 | 0 | 0 | 0 | 0 | 0 | 0 | 0 | 0 | 0 | 2 | 0 | 0 | 0 | 0 | 0 | 0 | 0 | 0 | 0 | 0 |
| Low affinity immunoglobulin gamma Fc region receptor III-A OS=Mus musculus OX=10090 GN=Fcgr4 PE=1 SV=1 | 0 | 0 | 0 | 0 | 0 | 0 | 0 | 0 | 0 | 0 | 0 | 2 | 0 | 0 | 0 | 0 | 0 | 0 | 0 | 0 | 0 | 0 | 0 | 0 | 0 | 0 |
| Glycogen phosphorylase, brain form OS=Mus musculus OX=10090 GN=Pygb PE=1 SV=3 | 0 | 0 | 0 | 0 | 0 | 0 | 0 | 0 | 0 | 0 | 0 | 0 | 0 | 0 | 2 | 0 | 0 | 0 | 0 | 0 | 0 | 0 | 0 | 0 | 0 | 0 |
| \| transcript=PBANKA_1349100.1 \| gene=PBANKA_1349100 \| organism=Plasmodium_berghei_ANKA \| gene_product=MSP7-like protein \| transcript_product=MSP7-like protein \| location=PbANKA_13_v3:1889651-1890658(-) \| protein_length=335 \| sequence_SO=chromosome \| SO=prote | 0 | 0 | 0 | 0 | 0 | 0 | 0 | 0 | 0 | 0 | 0 | 2 | 0 | 0 | 0 | 0 | 0 | 0 | 0 | 0 | 0 | 0 | 0 | 0 | 0 | 0 |
| Tubulin beta-5 chain OS=Mus musculus OX=10090 GN=Tubb5 PE=1 SV=1 | 0 | 0 | 0 | 0 | 0 | 0 | 0 | 0 | 0 | 0 | 0 | 0 | 0 | 0 | 0 | 0 | 0 | 0 | 0 | 0 | 0 | 0 | 0 | 3 | 0 | 0 |
| \| transcript=PBANKA_1127800.1 \| gene=PBANKA_1127800 \| organism=Plasmodium_berghei_ANKA \| gene_product=DnaJ protein, putative \| transcript_product=DnaJ protein, putative \| location=PbANKA_11_v3:1041667-1042806(-) \| protein_length=379 \| sequence_SO=chromosome | 0 | 0 | 0 | 0 | 0 | 0 | 0 | 0 | 0 | 0 | 0 | 0 | 0 | 0 | 0 | 2 | 0 | 0 | 0 | 0 | 0 | 0 | 0 | 0 | 0 | 0 |
| \| transcript=PBANKA_1301300.1 \| gene=PBANKA_1301300 \| organism=Plasmodium_berghei_ANKA \| gene_product=trailer hitch homolog \| transcript_product=trailer hitch homolog \| location=PbANKA_13_v3:88173-89186(-) \| protein_length=337 \| sequence_SO=chromosome \| SO=p | 0 | 0 | 0 | 0 | 0 | 0 | 0 | 0 | 0 | 2 | 0 | 0 | 0 | 0 | 0 | 0 | 0 | 0 | 0 | 0 | 0 | 0 | 0 | 0 | 0 | 0 |

| | | | | | | | | | | | | | | | | | | | | | | | | | | |
|---|---|---|---|---|---|---|---|---|---|---|---|---|---|---|---|---|---|---|---|---|---|---|---|---|---|---|
| \| transcript=PBANKA_1354500.1 \| gene=PBANKA_1354500 \| organism=Plasmodium_berghei_ANKA \| gene_product=60S ribosomal protein L18-2, putative \| transcript_product=60S ribosomal protein L18-2, putative \| location=PbANKA_13_v3: 2060347-2061576(+) \| protein_length | 1 | 0 | 0 | 0 | 2 | 0 | 0 | 0 | 0 | 0 | 0 | 0 | 0 | 0 | 0 | 0 | 0 | 0 | 0 | 0 | 0 | 0 | 0 | 0 | 0 | 0 |
| \| transcript=PBANKA_1434700.1 \| gene=PBANKA_1434700 \| organism=Plasmodium_berghei_ANKA \| gene_product=clathrin heavy chain, putative \| transcript_product=clathrin heavy chain, putative \| location=PbANKA_14_v3:1303281-1309241(+) \| protein_length=1986 \| sequen | 0 | 0 | 0 | 0 | 0 | 0 | 0 | 0 | 0 | 0 | 0 | 3 | 0 | 0 | 0 | 0 | 0 | 0 | 0 | 0 | 0 | 0 | 0 | 0 | 0 | 0 |
| Adenosylhomocysteinase OS=Mus musculus OX=10090 GN=Ahcy PE=1 SV=3 | 0 | 0 | 0 | 0 | 0 | 0 | 0 | 0 | 0 | 0 | 0 | 1 | 0 | 0 | 0 | 0 | 0 | 0 | 0 | 0 | 0 | 0 | 0 | 2 | 0 | 0 |
| Transferrin receptor protein 1 OS=Mus musculus OX=10090 GN=Tfrc PE=1 SV=1 | 0 | 0 | 0 | 0 | 0 | 0 | 0 | 0 | 0 | 0 | 0 | 0 | 0 | 0 | 0 | 0 | 0 | 0 | 0 | 0 | 0 | 0 | 0 | 0 | 2 | 0 |
| Lysozyme C OS=Gallus gallus GN=LYZ PE=1 SV=1 | 0 | 0 | 0 | 0 | 0 | 0 | 0 | 0 | 0 | 0 | 0 | 0 | 0 | 0 | 0 | 0 | 0 | 0 | 0 | 3 | 0 | 0 | 0 | 0 | 0 | 0 |
| Beta-adducin OS=Mus musculus OX=10090 GN=Add2 PE=1 SV=4 | 0 | 0 | 0 | 0 | 0 | 0 | 0 | 0 | 0 | 0 | 0 | 3 | 0 | 0 | 0 | 0 | 0 | 0 | 0 | 0 | 0 | 0 | 0 | 0 | 0 | 0 |
| Sarcoplasmic/endoplasmic reticulum calcium ATPase 1 OS=Mus musculus OX=10090 GN=Atp2a1 PE=1 SV=1 | 0 | 0 | 0 | 0 | 0 | 0 | 0 | 0 | 0 | 0 | 0 | 0 | 0 | 0 | 3 | 0 | 0 | 0 | 0 | 0 | 0 | 0 | 0 | 0 | 0 | 0 |
| Tropomyosin 3, related sequence 7 OS=Mus musculus OX=10090 GN=Tpm3-rs7 PE=3 SV=1 | 0 | 0 | 0 | 0 | 0 | 0 | 0 | 0 | 0 | 0 | 0 | 0 | 0 | 0 | 0 | 0 | 2 | 0 | 0 | 0 | 0 | 0 | 0 | 0 | 0 | 0 |
| \| transcript=PBANKA_0614500.1 \| gene=PBANKA_0614500 \| organism=Plasmodium_berghei_ANKA \| gene_product=eukaryotic translation initiation factor 3 subunit I, putative \| transcript_product=eukaryotic translation initiation factor 3 subunit I, putative \| locatio | 0 | 0 | 0 | 0 | 0 | 0 | 0 | 0 | 0 | 2 | 0 | 0 | 0 | 0 | 0 | 0 | 0 | 0 | 0 | 0 | 0 | 0 | 0 | 0 | 0 | 0 |
| Integrin-linked protein kinase OS=Mus musculus OX=10090 GN=Ilk PE=1 SV=2 | 0 | 0 | 0 | 0 | 0 | 0 | 0 | 0 | 0 | 0 | 0 | 0 | 0 | 0 | 0 | 0 | 2 | 0 | 0 | 0 | 0 | 0 | 0 | 0 | 0 | 0 |
| \| transcript=PBANKA_0932000.1 \| gene=PBANKA_0932000 \| organism=Plasmodium_berghei_ANKA \| gene_product=rhoptry neck protein 4 \| transcript_product=rhoptry neck protein 4 \| location=PbANKA_09_v3:1166868-1169639(+) \| protein_length=786 \| sequence_SO=chromosome | 0 | 0 | 0 | 0 | 0 | 0 | 0 | 0 | 0 | 0 | 0 | 2 | 0 | 0 | 0 | 0 | 0 | 0 | 0 | 0 | 0 | 0 | 0 | 0 | 0 | 0 |
| \| transcript=PBANKA_1360300.1 \| gene=PBANKA_1360300 \| organism=Plasmodium_berghei_ANKA \| gene_product=DNA/RNA-binding protein Alba 4, putative \| transcript_product=DNA/RNA-binding protein Alba 4, putative \| location=PbANKA_13_v3:2295446-2296682(-) \| protein_ | 0 | 0 | 0 | 0 | 0 | 0 | 0 | 0 | 0 | 0 | 0 | 0 | 0 | 0 | 0 | 2 | 0 | 0 | 0 | 0 | 0 | 0 | 0 | 0 | 0 | 0 |
| Beta-parvin OS=Mus musculus OX=10090 GN=Parvb PE=1 SV=1 | 0 | 0 | 0 | 0 | 0 | 0 | 0 | 0 | 0 | 0 | 0 | 0 | 0 | 0 | 0 | 0 | 2 | 0 | 0 | 0 | 0 | 0 | 0 | 0 | 0 | 0 |
| Phospholipid transfer protein OS=Mus musculus OX=10090 GN=Pltp PE=1 SV=1 | 0 | 0 | 0 | 0 | 0 | 0 | 0 | 0 | 0 | 0 | 0 | 0 | 0 | 0 | 0 | 0 | 0 | 0 | 0 | 0 | 0 | 0 | 0 | 2 | 0 | 0 |
| Kappa-casein OS=Bos taurus GN=CSN3 PE=1 SV=1 | 0 | 0 | 0 | 0 | 0 | 0 | 0 | 0 | 0 | 0 | 0 | 0 | 0 | 0 | 2 | 0 | 0 | 0 | 0 | 0 | 0 | 0 | 0 | 0 | 0 | 0 |
